# Assessing computational predictions of antimicrobial resistance phenotypes from microbial genomes

**DOI:** 10.1101/2024.01.31.578169

**Authors:** Kaixin Hu, Fernando Meyer, Zhi-Luo Deng, Ehsaneddin Asgari, Tzu-Hao Kuo, Philipp C. Münch, Alice C. McHardy

## Abstract

The advent of rapid whole-genome sequencing has created new opportunities for computational prediction of antimicrobial resistance (AMR) phenotypes from genomic data. Both rule-based and machine learning (ML) approaches have been explored for this task, but systematic benchmarking is still needed. Here, we evaluated four state-of-the-art ML methods (Kover, PhenotypeSeeker, Seq2Geno2Pheno, and Aytan-Aktug), an ML baseline, and the rule-based ResFinder by training and testing each of them across 78 species–antibiotic datasets, using a rigorous benchmarking workflow that integrates three evaluation approaches, each paired with three distinct sample splitting methods. Our analysis revealed considerable variation in the performance across techniques and datasets. Whereas ML methods generally excelled for closely related strains, ResFinder excelled for handling divergent genomes. Overall, Kover most frequently ranked top among the ML approaches, followed by PhenotypeSeeker and Seq2Geno2Pheno. AMR phenotypes for antibiotic classes such as macrolides and sulfonamides were predicted with the highest accuracies. The quality of predictions varied substantially across species–antibiotic combinations, particularly for beta-lactams; across species, resistance phenotyping of the beta-lactams compound, aztreonam, amox-clav, cefoxitin, ceftazidime, and piperacillin/tazobactam, alongside tetracyclines demonstrated more variable performance than the other benchmarked antibiotics. By organism, *C. jejuni* and *E. faecium* phenotypes were more robustly predicted than those of *Escherichia coli*, *Staphylococcus aureus*, *Salmonella enterica*, *Neisseria gonorrhoeae*, *Klebsiella pneumoniae*, *Pseudomonas aeruginosa*, *Acinetobacter baumannii*, *Streptococcus pneumoniae*, and Mycobacterium tuberculos*is*. In addition, our study provides software recommendations for each species–antibiotic combination. It furthermore highlights the need for optimization for robust clinical applications, particularly for strains that diverge substantially from those used for training.

## Introduction

The efficacy of antimicrobials, compounds that can eliminate or inhibit microorganisms, is increasingly undermined by the problem of antimicrobial resistance (AMR) [1–3]. As one of the top 10 public health threats identified by the World Health Organization [4,5], bacterial AMR was responsible for an alarming 1.27 million deaths worldwide in 2019 [6]. The conventional culture-based antimicrobial susceptibility testing currently guiding therapy [7] is time-consuming and limited to pure bacterial cultures [1,7–10]. Marker-based molecular diagnostics offer a faster alternative but cover only a few known loci, making them prone to false negative results [9,11,12]. In contrast, whole-genome sequencing, by providing access to all genes, offers an approach to determining the AMR phenotype that is potentially more accurate than the current molecular methods [8,13,14] and, in combination, with metagenomic analytics, more rapid than culture-based testing for biomedical applications [15–17], which, in future, could complement or replace the standard antimicrobial susceptibility testing for AMR phenotyping in clinical diagnostics, surveillance, and containment [1,8,13,14].

Two main computational approaches have been developed for predicting AMR phenotypes from genomic data: rule-based AMR catalog mapping (referred to as rule-based) and machine learning (ML) approaches. Rule-based approaches determine AMR phenotypes using catalogs of known single nucleotide polymorphisms (SNPs), indels, and/or genes associated with AMR, comparing the sample with reference catalogs [10,18–34]. Although this approach can indicate the underlying AMR mechanisms by pinpointing SNPs and genes, it is inherently constrained by the scope, quality, and interpretation of the catalog or database [35].

The accumulation of genomes with AMR metadata [36] provides a rich resource for developing ML-based AMR phenotyping approaches, offering a strategy that is potentially more versatile and accurate for predicting AMR. These ML approaches use various classification algorithms for identifying AMR patterns from the data (**Table S1**). ML models utilizing feature representation of k-mers [37–42], pan-genomes [43–46], and genome-wide or common mutations [43,47–52] can predict AMR phenotype without an understanding of the molecular mechanism. Additionally, some ML-based approaches also encode information on the determinants of AMR into features [45,48,50,51,53–65], and some research has utilized ML techniques for detecting AMR determinants [43,46,66–71].

A direct consequence of the diverse landscape of AMR prediction methodologies is the inherent difficulty in benchmarking and comparing their effectiveness. Although newly developed methods often highlight their advances compared with existing methods, the evaluations are not standardized, leading to the use of different datasets, different yet limited combinations of species and antibiotics, and a lack of comprehensive comparisons with the full spectrum of available methods. ML-based AMR phenotyping is assessed by different approaches (**Table S1**). Most straightforwardly, performance can be compared on the basis of the same set of features across various classifiers [37,38,40,43,47,50,53,57,58,72]. Further, benchmarks can be compared using features derived from various types of information, such as different AMR reference databases [73], complete sets and subsets of features (SNPs, genes, or k-mers) [41,48,49,51,61–63,65,69,70,74], and combinations of features (pan-genomes, SNPs, population or non-genomic information) [43–45]. Other studies have compared ML-based software with rule-based software [44,48,55,61–63,65], or with another type of ML-based software on only one species [52,54,59,64,75], or have assessed a larger group of methods only on *M. tuberculosis* [51,59,60,64]. Of these studies, all except two, used up to five pathogen species for benchmarking (**Supplemental Methods**). The field also lacks a more comprehensive comparison covering both a wide range of pathogens and the wide spectrum of ML-based AMR phenotyping software to survey whether ML methods are generally more competitive than the conventional ones. Therefore, establishing standardized benchmarking procedures for a systematic comparison of ML-based methods on a wide range of species remains a significant research gap in the field of computational predictions of AMR phenotypes from microbial genomes.

To establish a comprehensive benchmark for evaluating AMR phenotyping, we developed a rigorous workflow in terms of datasets, guidelines, and metrics. We curated 78 datasets from the PATRIC database, representing different combinations of species and antibiotics. We then evaluated the AMR phenotyping software using one of three defined approaches (nested cross-validation (CV), iterative bound selection [38] with CV, or iterative evaluation) to accommodate different techniques, each in conjunction with three methods of partitioning the sample (random folds, phylogeny-aware folds, homology-aware folds). We evaluated the ML-based methods in both single-species or single-antibiotic models and models with multiple species or antibiotics. We measured the performance with a set of common metrics, prioritizing the macro F1-score for a comprehensive analysis, and the precision and F1-score for the susceptible samples for a clinically oriented analysis. Within this framework, we conducted a thorough evaluation of state-of-the-art AMR phenotyping methods, namely Aytan-Aktug [55], Seq2Geno2Pheno [43], PhenotypeSeeker [40], and Kover [38], alongside the rule-based ResFinder 4.0 [21] and ML baseline [38]. Our study aims to guide researchers in selecting the most suitable software for specific pathogens and antimicrobials, and to indicate the strengths and weaknesses of the current techniques in the field.

## Results

### Development of a rigorous transparent benchmarking workflow for predicting AMR

To perform an unbiased and thorough comparison of AMR prediction software, we developed a benchmarking workflow using three distinct sample splitting methods and providing common performance metrics (**Fig. 1A**). Within this workflow, the methods under evaluation were trained and tested on the predefined splits. This comprehensive approach, in terms of software, antimicrobial, and species coverage, combined with our rigorous workflow, provides a balanced assessment, thoroughly examining the capacity for generalization across a spectrum of scenarios. To promote transparency and enable the replication of our results, we have made the workflow available under an open-source license (http://github.com/hzi-bifo/AMR_benchmarking).

**Figure 1:**
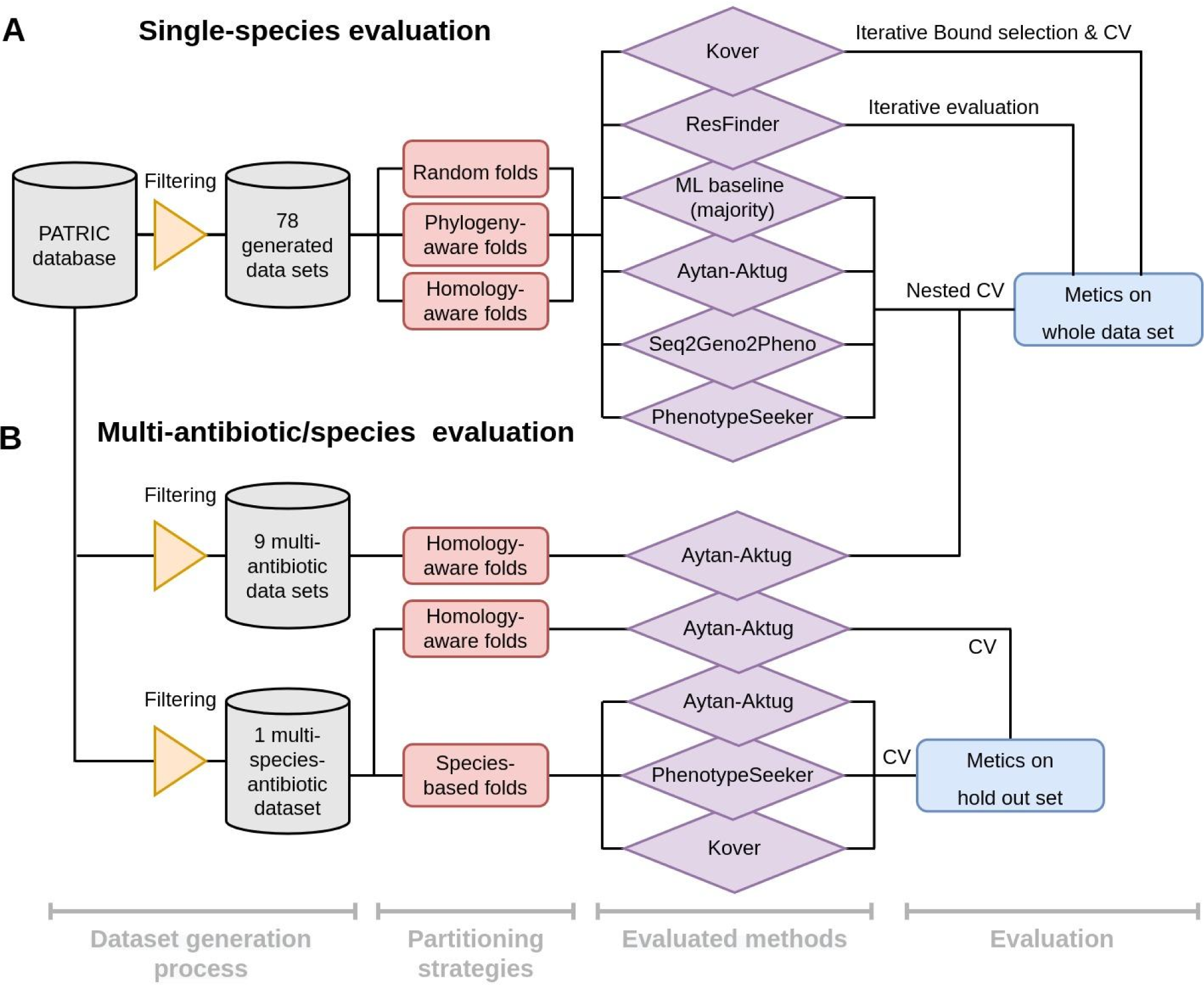
Overview of the evaluation workflow for predicting antimicrobial resistance (AMR), spanning single-species and multi-species models, and various data processing strategies. (A) The process of evaluating the single-species model, using 78 datasets created from the PATRIC database. Software for determining the AMR phenotype is evaluated through one of three approaches (nested cross-validation (CV), iterative bound selection [38] with CV, or iterative evaluation), each in conjunction with three sample partitioning methods (random folds, phylogeny-aware folds, and homology-aware folds). The outcomes are expressed as the mean and standard deviation for various model-oriented metrics (F1-macro, F1-negative, F1-positive, accuracy) and clinically relevant metrics (F1-negative, precision-negative, and recall-negative), benchmarked against phenotypes inferred *in silico* and culture-based antimicrobial susceptibility testing standards. (B) Evaluation of multi-antibiotic or multi-species models illustrating the evaluation workflow for the Aytan-Aktug single-species multi-antibiotic model; two types of Aytan-Aktug models with multiple species and multiple antibiotics; and cross-species models using Aytan-Aktug, Kover, and PhenotypeSeeker techniques. The Aytan-Aktug single-species multi-antibiotic model was evaluated across nine multi-antibiotic datasets using nested CV with homology-aware partitioning. Two types of Aytan-Aktug models with multiple species and multiple antibiotics were evaluated on a dataset with multiple species and multiple antibiotics using conventional CV with a holdout test set on samples partitioned by sequence similarity, or using conventional CV with a holdout test set obtained by species-based stratification. Aytan-Aktug (multi-antibiotic), Kover (single-antibiotic), and PhenotypeSeeker (single-antibiotic) cross-species models are evaluated by leave-one-species-out on the dataset with multiple species and multiple antibiotics. Metrics are reported for each combination in the holdout set in the case of conventional CV.

ML-based AMR phenotyping has inherent challenges, notably the ability of models to be generalized to new input data. To assess the AMR phenotyping software in various scenarios, we used three distinct sample splitting methods (random partitioning, phylogeny-based, and homology-based approaches) to reflect different degrees of evolutionary divergence or dissimilarity between taxa in the training and test data, and to balance realistic evaluation with statistical reliability (**Supplemental Fig. S3, Supplemental Methods**). To ensure reproducibility, and maintain consistency and uniformity across evaluations, each splitting method was applied uniformly to every dataset, resulting in the same three dataset partitions for assessing the evaluated methods. In addition to the three sample partitioning methods for the single-species model (**Fig. 1A**), we applied a challenging species-based sample partitioning method for evaluating cross-species models, using leave-one-species-out (LOSO) evaluation (**Fig. 1B**). The hyperparameter optimizations and performance evaluations of the models were implemented using nested CV designed to mitigate the bias in the error estimated by CV [76], as well as two iterative evaluations, in order to accommodate different techniques.

We measured each method’s performance using a set of common metrics: precision, quantifying the ratio of correct predictions to all predictions for a given class; recall, measuring the proportion of correctly assigned instances of a particular class; and the F1-score, providing a balanced measure across these metrics for each class. Additionally, we assessed the overall assignment accuracy across the entire dataset and the macro F1-score, offering a performance estimate across both resistant and susceptible classes, independent of the class sizes (**Fig. 1A,B**). Considering the comprehensiveness of the assessment (**Supplemental Methods**) for an overall comparison, we used the mean of the macro F1-scores (referred to as the F1-macro mean), averaged across the 10 folds. To assess the clinical relevance of the evaluated methods, the precision and the F1-score for the susceptible (negative) class were used, recognizing their critical role in identifying actionable antibiotic susceptibilities.

### Assessing software for a comprehensive benchmarking of AMR phenotyping

To select state-of-the-art ML-based software for our study, we followed a systematic approach. Initially, we limited our search to software released since 2017 and with available source codes, ensuring relevance and accessibility for reproducibility, resulting in 17 methods (**Supplemental Table S1**). We then further considered the universal applicability of the software, excluding those that were species-specific (most were only on one species, with one study on three species), requiring a reference strain (usually solely SNP-based software [47–50,52,60,62,63,65]) or an AMR gene catalog [48,59,60,62,63,65] for generating features, resulting in seven methods [38,40,43,44,54,55,74].

We also aimed to cover a wide spectrum of ML classifiers. The selected methods included both common ML classifiers such as support vector machine, logistic regression, and random forest (used by Seq2Geno2Pheno [43] and PhenotypeSeeker [40]), and neural networks (used by Aytan-Aktug [55]). Additionally, we included more novel approaches such as the bound selection rule-based learning models of Kover [38]. The variety in the selected classifiers provided a broad and representative overview of the different algorithmic approaches available in the field, showcasing both established techniques and innovative methodologies.

Crucially, we also selected methods so that they also provided a comprehensive comparison of ML methods based on various features that can be derived from the genomic dataset that we used for benchmarking. These features can generally be categorized into k-mers, pan-genomes (including presence or absence of genes, sequence similarity clusters, SNPs, indels), AMR catalog-encoded features (presence or absence of AMR-associated SNPs and genes), SNPs, and non-genomic features (such as gene expression and year of isolation; **Supplemental Table S1**). Our selection thus included the ML-based methods Aytan-Aktug (AMR catalog based) [55], Seq2Geno2Pheno (pan-genomes and k-mer counting) [43], PhenotypeSeeker [40] and Kover [38] (both using the presence or absence of k-mers). Additionally, we incorporated the AMR catalog mapping method ResFinder 4.0 [21], which is recognized for its use of comprehensive and regularly updated databases, including both resistance genes and SNPs, as well as a methodological ML baseline that predicted the most frequently occurring phenotype in the training data for each experiment [38]. Overall, this selection strategy reflects the various ways in which genomic information can be utilized for predicting AMR. To evaluate the software within our rigorous workflow, we made some adaptations or refinements of ResFinder, Aytan-Aktug, Seq2Geno2Pheno, and PhenotypeSeeker (**Supplemental Methods**, https://github.com/hzi-bifo/AMR_benchmarking_khu).

### Creation of comprehensive benchmark datasets for AMR phenotyping

We curated a collection of 78 datasets from the PATRIC AMR database [77,78], each including microbial genomes for a distinct combination of species and antibiotics (Fig. 2A**, Supplemental Fig. S1**). These genomes were selected after a filtering process based on specific criteria, ensuring their quality and relevance. Specifically, we considered the antimicrobial susceptibility testing phenotype annotation, sequence type (excluding plasmid-only sequences), contig number, genome length, and PATRIC scores for genome quality, fine consistency, coarse consistency, and completeness. This filtering process was crucial for excluding any inconsistencies, inaccuracies, or irrelevant information that could skew the assessment or reduce the reliability of the evaluation.

**Figure 2:**
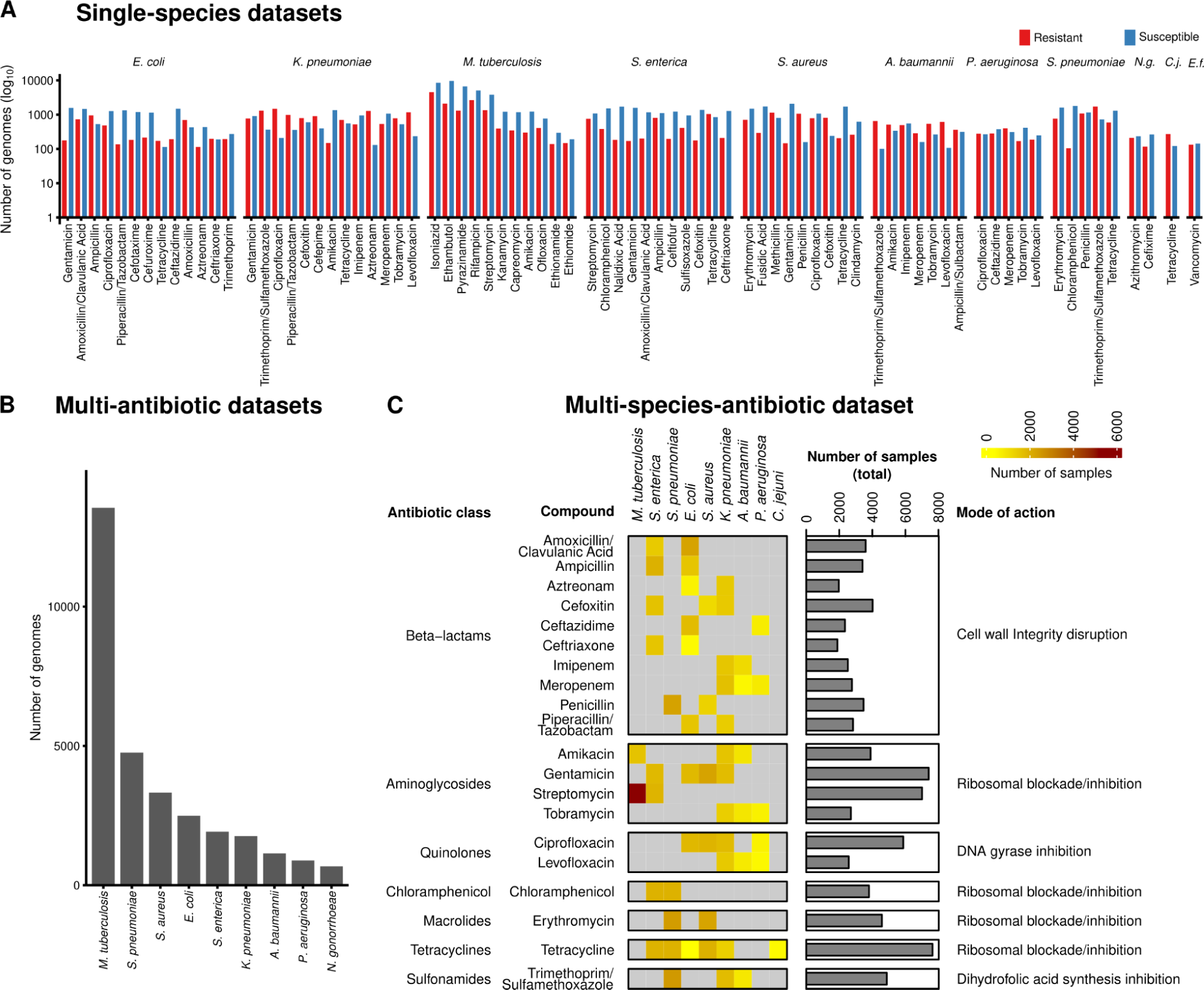
Overview of datasets generated for evaluating AMR phenotyping. (A) Seventy-eight datasets for the single-species evaluation (**Fig. 1A**), spanning 11 species, annotated with resistant (red) and susceptible (blue) genome isolates. Abbreviation: *N.g.*: *N. gonorrhoeae*; *C.j.*: *C. jejuni; E.f.*: *E. faecalis*. (B) Number of genomes included in the multi-antibiotic datasets for evaluating multi-antibiotic models (**Fig. 1B**). (C) Overview of the dataset with multiple species and antibiotics, spanning nine species and 20 antibiotics for evaluating multi-species models (**Fig. 1B**).

The 78 datasets, ranging from 200 to 13,500 genomes each and encompassing 31,195 genomes in total, provide coverage of at least 100 genomes for each evaluated phenotype class (resistant or susceptible to a certain antibiotic). The datasets include genomes from 11 bacterial species, including key pathogens and species with high clinical relevance. These include eight high-priority pathogens linked to the most AMR-related deaths [6], namely *Escherichia coli*, *Staphylococcus aureus*, *Klebsiella pneumoniae*, *Streptococcus pneumoniae*, *Acinetobacter baumannii*, *Pseudomonas aeruginosa*, *Mycobacterium tuberculosis*, and *Enterococcus faecium*, as well as *Salmonella enterica*, which is among the 11 most concerning pathogens. We further incorporated the genomes of two additional species of clinical and biological relevance, namely *Campylobacter jejuni*, the most common bacterial cause of gastroenteritis with the rise of AMR [79,80], and *Neisseria gonorrhoeae*, which poses a significant clinical challenge with the emergence of pan-resistant isolates [81]. For each species, we compiled genomes along with their phenotypic annotations, indicating susceptibility or resistance to a given antimicrobial. The resulting 44 antimicrobials span 14 major drug classes such as beta-lactams, fluoroquinolones, aminoglycosides, tetracyclines, folate pathway inhibitors, lipopeptides, oxazolidinones, tuberculosis-specific drugs, and combination therapies (**Supplemental Table S2**). The coverage of these combinations aimed to reflect the complexity of AMR across diverse species associated with a variety of infectious diseases and potential resistance mechanisms [81,82].

We also created more complex datasets to extend the scope of the evaluation from single-species single-label models to multi-label and multi-species AMR phenotyping models (**Supplemental Methods**). First, we generated nine multi-antibiotic datasets, each containing genomes from one of nine species that were annotated with resistance phenotypes from all the antibiotics compiled in that species’ datasets with single species and antibiotics (**Fig. 2B**). For example, the *E. coli* multi-AMR phenotype dataset encompassed 2493 genomes annotated with AMR phenotypes for up to 13 antimicrobials. Second, we created a dataset of multiple species and antimicrobials covering nine key species (such as *S. aureus*, *K. pneumoniae*, *P. aeruginosa*, and *M. tuberculosis*) annotated with resistance against 20 antimicrobials, encompassing 20,989 genomes (**Fig. 2C, Mendeley Data**). This covers antimicrobials that occur in at least two of the 78 species–antimicrobial combinations. In total, the multi-species dataset covers 54 species–antimicrobial combinations.

### Comparative assessment of AMR phenotyping software: performance, robustness, and applicability

We critically evaluated the selected AMR phenotyping methods by assessing the software’s performance across a variety of conditions, exploring the quality of performance and robustness to provide insights into the practical applicability of these methods under different conditions. The software’s predictive quality and the uncertainty of these estimates were measured by the F1-macro mean and F1-macro standard deviation, respectively, across the 10 evaluation folds. The software’s robustness refers to the stability of performance when the trained software was evaluated on test sets with genomes with various phylogenies or sequence similarity relationships to the training set. The smaller the change in performance across different evaluation scenarios, the greater the software’s robustness and the more the trained ML-based software can be trusted by users to provide stable and accurate predictions on samples with unknown relationships to the training data.

We evaluated the five methods (four ML-based and the rule-based method ResFinder) on 78 datasets with random and homology-aware dataset splitting, as well as phylogeny-aware splitting on 67 datasets (11 datasets splitting failed because of runtime issues). For each method, the configurations of the hyperparameter selection range are specified in the **Supplemental Methods**. Seq2Geno2Pheno failed to generate pan-genome-based features for *M. tuberculosis*, and PhenotypeSeeker could not complete the CV evaluation for four *M. tuberculosis*–antibiotic combination datasets because of computational time exceeding 2 months with a maximum of 20 available cores. For these experiments, metric values of 0 were assigned. Additionally, 13 of the 78 species–antibiotic combinations could not be assessed by ResFinder, owing to AMR catalogs being absent, with the metric values also consequently set to 0. In total, with these 78 species–antibiotic combinations (referred to as cases), we conducted 390 (78 datasets x 4 ML methods + 78 ResFinder) experiments each for random folds and homology-aware folds, and 335 (67 datasets x 4 ML methods + 67 ResFinder) experiments with phylogeny-aware folds (see **Supplemental Table S3** for the metric scores). To assess the methods’ performance, we first compared these methods with the methodological ML baseline for each experiment [38]. With the exception of six experiments and 30 experiments that failed because of excessive training times, the ML methods outperformed (better or equal to) the ML baseline in 96% of the experiments. Notably, the performance of ResFinder was comparably worse, primarily because of the absence of AMR catalogs for specific datasets in 37 of 43 underperforming experiments, surpassing the ML baseline only in 78% of the experiments (**Supplemental Table S4A**).

To further compare the performance of the various methods, we ranked them on the basis of the frequency that each performed the best (see **Fig. 3** for a heatmap of the F1-macro metrics, **Table 1**). A method performed the best by having the highest F1-macro mean or the lowest standard deviation in case of ties of the highest mean. Methods were assigned an equal fraction in a tie of the mean and standard deviation for a particular species–antibiotic combination. Although the ML methods’ predictive capacity varied, they maintained a generally declining trend across evaluation settings: these methods performed best in random split evaluations, performed less well in phylogeny-based evaluations when the strains in the test set were more divergent from the training data, and the least well in homology-based evaluations, with Kover being the notable exception. Across all settings, Kover most frequently emerged as the top performer, followed by PhenotypeSeeker, Seq2Geno2Pheno, and the Aytan-Aktug models (**Table 1**). In the context of random split evaluations, Kover significantly outperformed the other methods (p = 0.01 and 3.27×10^-7^, compared with PhenotypeSeeker and ResFinder, respectively, using one-sided paired *t*-tests), achieving the highest F1-macro values for 30% of the species–antibiotic combinations. ResFinder and PhenotypeSeeker followed, with 25% and 20%, respectively (**Table 1**). On the other hand, ResFinder demonstrated a particular strength in predicting AMR phenotypes for evolutionarily distant strains from the training samples. During the phylogeny- and homology-based evaluations, ResFinder performed the best on 44% and 50% of the datasets, respectively, followed by Kover, with 28% and 34%, respectively (**Table 1**). Despite these findings, the lack of AMR catalogs and the consequent inability to offer predictions for 17% species–drug combinations meant that ResFinder did not prove to be significantly better than Kover overall (**Fig. 3, Fig. 4A**). In addition, we evaluated the reproducibility of ML methods, showing that all ML-based methods exhibited some variability in their results caused by stochastic factors, except for PhenotypeSeeker (**Supplemental Results**).

**Figure 3:**
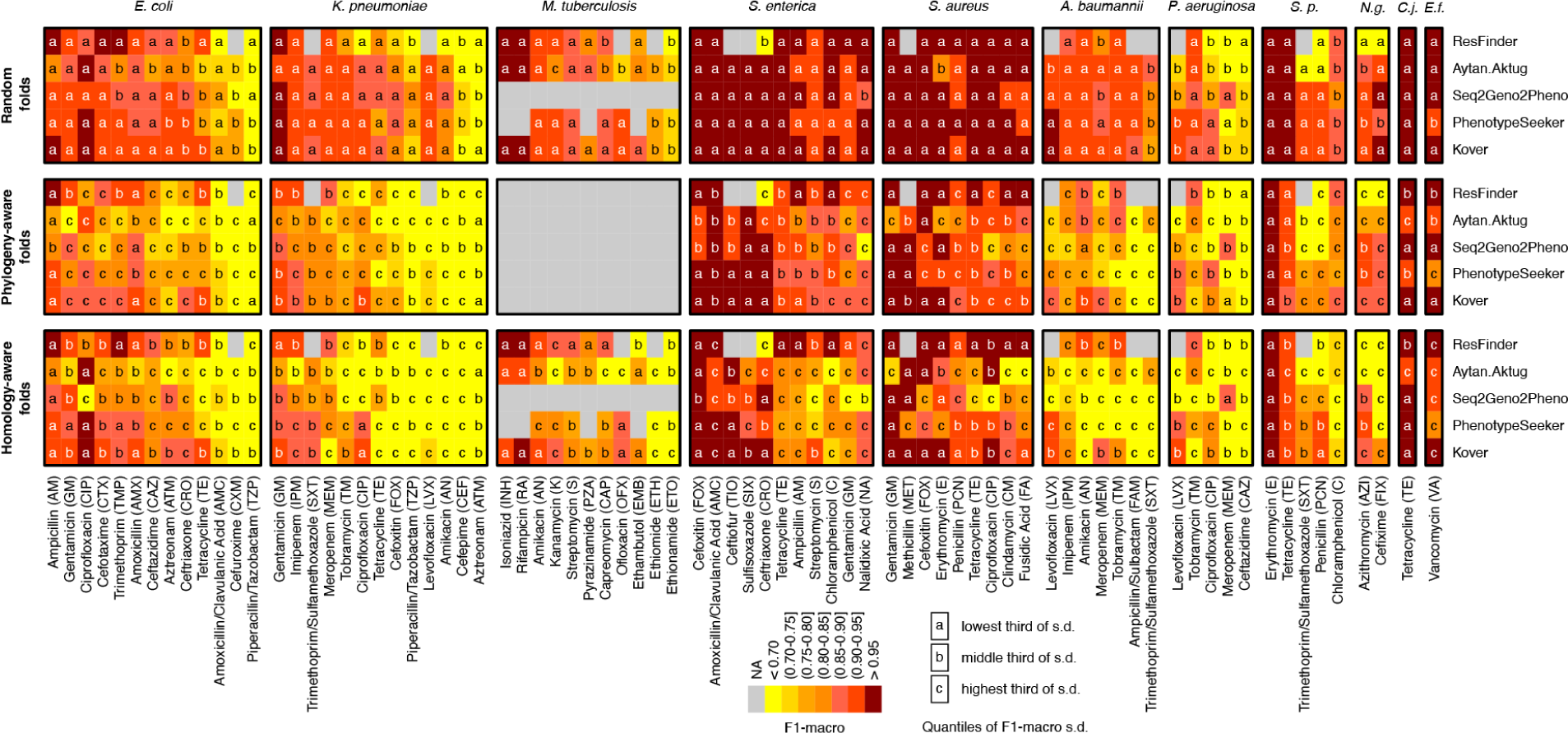
Heatmaps comparing AMR phenotyping performance (F1-macro) with random folds, phylogeny-aware folds, and homology-aware folds for the methods evaluated on 78 species–antibiotic combinations. The gray cells indicate either the absence of an AMR catalog for ResFinder, or the failure of ML-based evaluations caused by running time issues. Colors represent the F1-macro mean, whereas the markers ‘a,’ ‘b,’ and ‘c’ denote the categories of the F1-macro standard deviation, which are ‘lowest’, ‘middle’, and ‘highest’, respectively, based on their tertiles. Abbreviation: *S.p*: *S. pneumoniae*; *N.g.*: *N. gonorrhoeae*; *C.j.*: *C. jejuni; E.f.*: *E. faecalis*.

**Figure 4:**
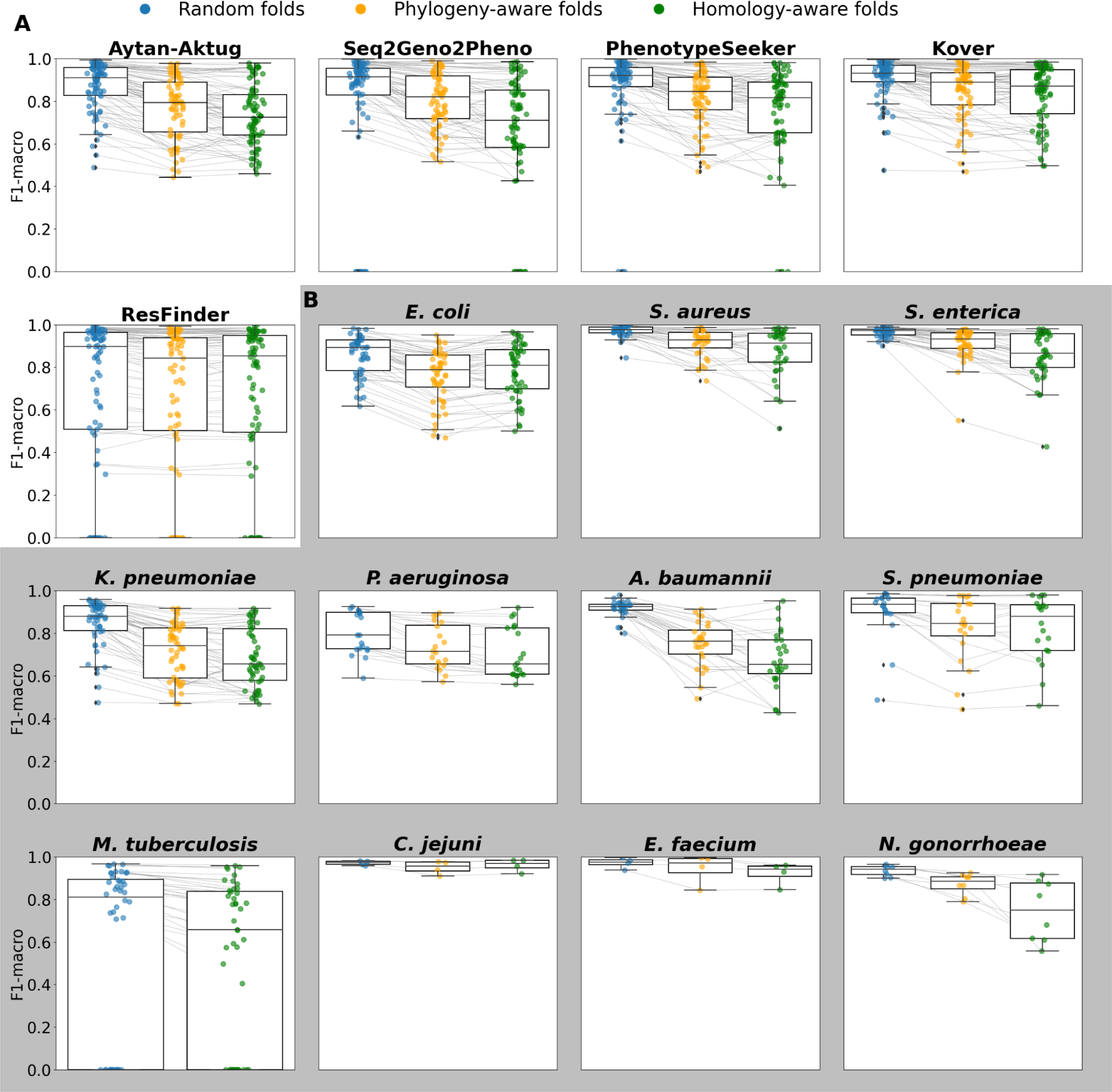
Changes in performance (F1-macro mean) under random folds, phylogeny-aware folds, and homology-aware folds. (A) Changes in performance for each method. Each dot represents one species–antibiotics combination for the corresponding method. (B) Changes in performance for each species for the ML-based methods Kover, PhenotypeSeeker, Seq2Geno2Pheno, and Aytan-Aktug. Each point represents one ML-based method–antibiotic combination for the corresponding species.

**Table 1:** Percentage of species–antibiotic combinations where each method performed the best, stratified by the method of splitting the dataset. Each dataset covers a specific species–antimicrobial combination. A method is considered to perform the best for a species–antimicrobial combination by having the highest F1-macro mean or the lowest standard deviation in case of ties of the highest mean (see **Supplemental Table S6** for the best-performing method for each species–antimicrobial combination). Methods were assigned a fraction of the count for a tie of the mean and standard deviation. The fractions in brackets stand for the number of corresponding winning combinations out of the total. Bold indicates the top-ranking method.

| AMR prediction method | Fraction of experiments where the AMR prediction method performed the best (by splitting method) |  |  |
| --- | --- | --- | --- |
|  | Random folds | Phylogeny-aware folds | Homology-aware folds |
| ResFinder | 25% (19.53/78) | **44% (29.25/67)** | **50% (38.75/78)** |
| Aytan-Aktug | 8% (6.12/78) | 6% (4.08/67) | 2% (1.58/78) |
| Seq2Geno2Pheno | 17% (13.28/78) | 10% (6.58/67) | 4% (3.25/78) |
| PhenotypeSeeker | 20% (15.37/78) | 13% (8.58/67) | 10% (7.92/78) |
| Kover | **30% (23.70/78)** | 28% (18.50/67) | 34% (26.50/78) |

Collectively, ML methods demonstrated very good performance, achieving an F1-macro mean of ≥0.9 in 64% of the experiments and ≥0.8 in 81% of the experiments during a random evaluation (**Table 2**). They were the best-performing methods in 64% of the species–antibiotic combinations, outdoing the AMR catalog mapping technique of ResFinder, which was best in 17% of the cases. In 19% of the cases, both approaches were tied (**Supplemental Table S4C**). However, the effectiveness of the ML methods dropped noticeably when applied to strains that were more evolutionarily divergent from the model’s training genomes. This decrease was pronounced in phylogeny-based evaluations and even more so in homology-based evaluations. In these settings, ML methods only achieved an F1-macro mean of ≥0.9 in 33% and 25% of the homology-based and phylogeny-based experiments, respectively, and an F1-macro mean of ≥0.8 in 60% and 50%, respectively. When compared with ResFinder, the ML methods achieved higher or equal F1-macro scores in fewer experiments (47% or 40%, respectively, for the homology-based and phylogeny-based experiments), with higher scores being observed in 40% and 35% of the experiments (**Table 2, Supplemental Table S4B**).

**Table 2:**
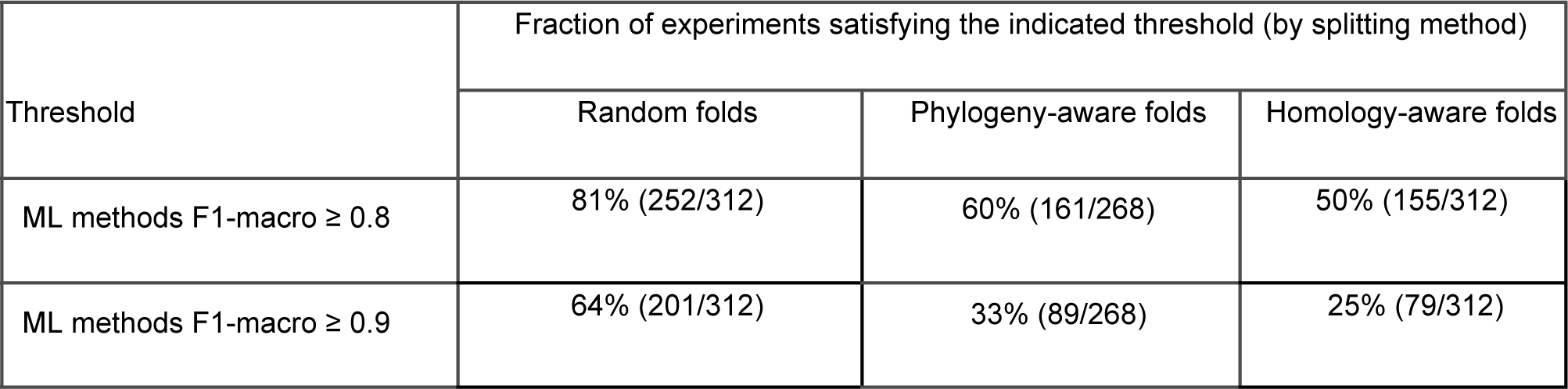
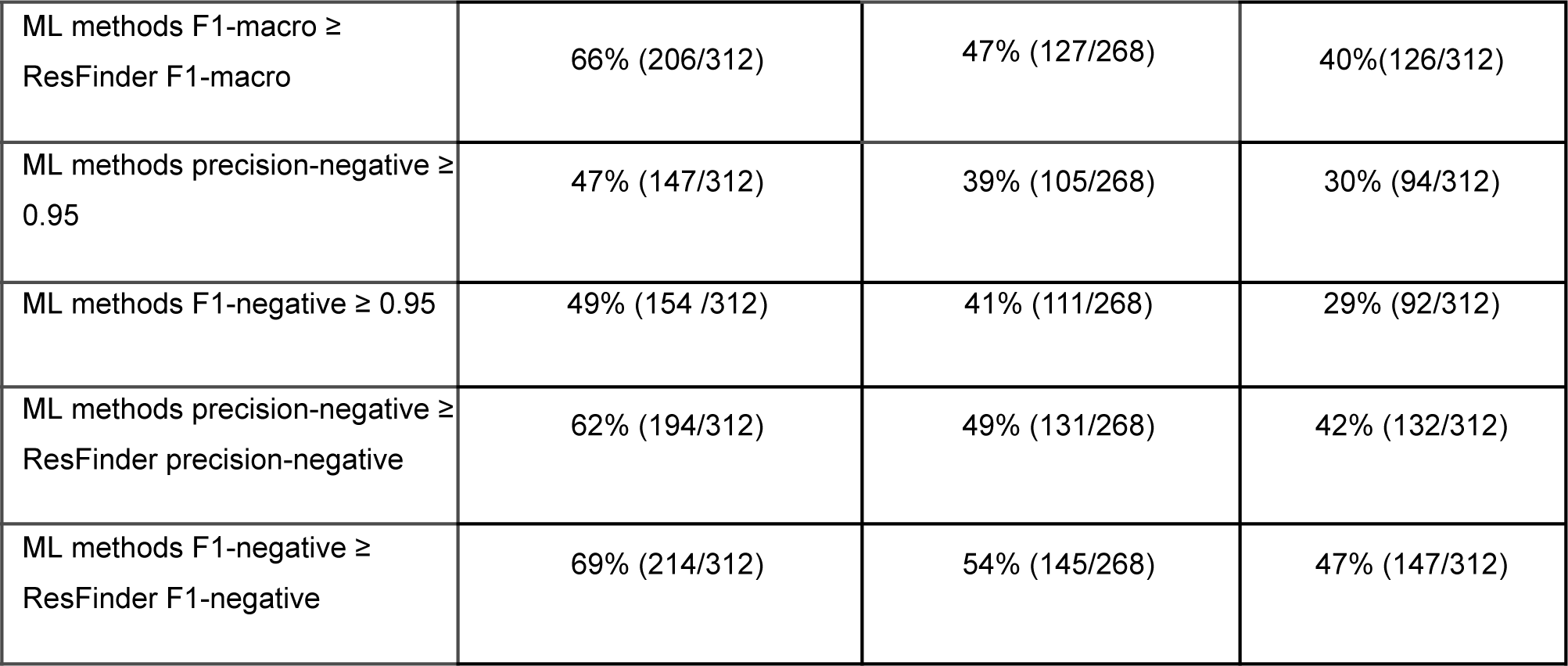
Percentage of experiments in which the machine learning (ML) methods performed above the indicated threshold, stratified by the method of splitting the dataset. The fractions in brackets stand for the number of corresponding experiments out of the total. In total, there were 312 (78 datasets x 4 ML methods) experiments with random folds, 312 experiments with homology-aware folds, and 268 (67 datasets x 4 ML methods) experiments with phylogeny-aware folds.

When considering the application to clinical diagnostics and treatment, predicting a pathogen’s susceptibility to a specific drug is of high importance [83]. Thus, high precision in detecting the susceptibility is essential [8]; moreover, high recall in detecting susceptibility is required for identifying viable treatment options. To this end, we evaluated the methods’ performances, with an emphasis on the F1-score and precision for the negative (i.e., susceptible) class (**Supplemental Table S3**). With the ML methods, a precision of at least 0.95 for the negative class was reached for 47% (147/312) of the experiments evaluating random splits. However, this decreased to 39% (105/268) and 30% (94/312) for the phylogeny-based and homology-based evaluations, respectively. Similarly, an F1-score for the negative class of ≥0.95 was achieved in 49% (154/312) of experiments with random splits, compared with 41% (111/268) and 29% (92/312) for the phylogeny-aware and homology-aware evaluations, respectively (**Table 2**). Thus, depending on the evolutionary divergence of the strains relative to the training data, the susceptibility of these pathogens to specific antibiotics can be predicted using ML methods with high confidence in almost one-third to half of the cases. Similar to evaluations measured by the F1-macro mean, with the metrics for the negative-class, ML methods performed better in evaluations with random splits, whereas AMR catalog mapping with ResFinder performed comparably well in the phylogeny-based and homology-based evaluations (**Table 2, Supplemental Results**).

### Species- and antibiotic-specific trends in the performance of AMR predictions

For two pathogen species, *C. jejuni* and *E. faecium*, the predictive quality remained consistently high across the three types of evaluation. This consistency highlights the capacity of all methods to robustly predict AMR phenotypes, even for strains that are more evolutionarily divergent (**Fig. 2, Fig. 4B** for the F1-macro mean; **Supplemental Fig. S10B** for the F1-macro standard deviation). In contrast, for the other nine species evaluated, the models’ performance experienced a decline in the more challenging scenarios involving test data from phylogenetically divergent strains relative to the training data. While for *S. aureus*, *S. enterica*, *A. baumannii*, *N. gonorrhoeae*, and *S. pneumoniae* ML methods showed a strong performance with random splits, there was a marked deterioration in the median of the F1-macro mean and increased variation in both the F1-macro mean and standard deviations in the more challenging settings. For species such as *E. coli*, *M. tuberculosis*, *K. pneumonia*, and particularly *P. aeruginosa*, the ML models did not perform quite as well (the median of F1-macro means was between 0.89 and 0.79) with random splits, and the performances declined further in the other settings. For all species, variance increased for the more challenging settings, indicating the increased uncertainty of the trained models. For all species, with the exception of *M. tuberculosis*, for which the phylogenetic tree could not be generated because of runtime issues, there was an uneven distribution of the misclassification frequencies across the corresponding phylogenetic tree (**Supplemental Results, Supplemental Fig. S13–22**). This indicated that AMR phenotyping across a range of drugs might be complicated by the background lineage.

Our results revealed substantial variations in the quality of predictions for different antibiotic classes (**Supplemental Fig. S8**) and across different antibiotics (**Supplemental Fig. S9**), excluding experiments with computing failures caused by runtime issues. For the antibiotic classes, the macrolides, sulfonamides, rifamycins, glycopeptides, and lipoglycopeptides tended to be more accurately predicted than the other antibiotic classes across the three evaluation scenarios. For the aminoglycosides, quinolones, chloramphenicol, tetracyclines, lincosamides, fusidane, TB drugs, and dihydrofolate reductase inhibitors, the AMR prediction methods generally achieved high accuracy in random evaluations, but experienced a decline in the two more challenging scenarios. Intriguingly, the performance for the beta-lactams class exhibited exceptionally high variation in the F1-macro means for both random splits and the two more challenging evaluation scenarios. Further, our analysis demonstrated variations among different species treated with the same antibiotic (**Supplemental Fig. S5—7**). This variation was particularly evident in the case of five beta-lactams (aztreonam, amoxicillin/clavulanic acid, cefoxitin, ceftazidime, and piperacillin/tazobactam) and tetracycline with random folds. These variations, except that for piperacillin/tazobactam, still appeared with phylogeny-aware and homology-aware folds. Moreover, the variation between individual species–antibiotic combinations was pronounced, with cefoxitin–*S. aureus* and aztreonam–*K. pneumoniae* predicted with the highest (F1-macro ≥ 0.99) and lowest accuracies (F1-macro mean = 0.59).

### Assessment of multi-antibiotic and multi-species AMR prediction models

A substantial number of sequenced genomes with AMR metadata, which are vital for training ML-based AMR phenotype prediction models, are not available for all species–antibiotic combinations. Similarly, high-quality AMR catalogs, which are crucial for developing ML-based AMR phenotyping software relying on AMR catalogs, are not available for all species–antibiotic combinations. Furthermore, multidrug-resistance, seen in strains like methicillin-resistant *S. aureus* and multidrug-resistant *M. tuberculosis*, often arises from interacting mechanisms, which are primarily intrinsic via multiple drug-resistance efflux systems [82] and acquired via mutations or mobile multidrug resistant genetic material [81,82,84–86]. AMR genes coding for multidrug efflux pumps are highly prevalent in all studied organisms [86]. It is of particular interest, therefore, to create models that are generalizing to multiple antibiotics and multiple species, thus potentially improving AMR phenotype predictions for less-studied taxa and taxa with limited data. Therefore, in addition to models with single species and antibiotics (**Fig. 1A**), we also evaluated multi-antibiotic models, which fall into the category of multi-label classification, as well as multi-species models trained on genomes from multiple species (**Fig. 1B**). For the multi-species models, we examined their performance under two conditions, first when applied to a species already included in the training data, and second when applied to a species the model had never encountered before.

To assess the multi-antibiotic model, we retrained and tested the Aytan-Aktug multi-label model on nine species with data available for multiple antibiotics, using homology-based data stratification and nested CV (**Supplemental Methods**). Similar to Aytan-Aktug et al. [55], we observed no substantial difference in the predictive performance of multi-antibiotic and single-antibiotic models, evaluated on nine species instead of only one, as in the original study (*p* = 0.48 with a paired *t*-test, **Supplemental Fig. S23**).

To access the multi-species model where multiple species were involved in both the training and testing procedures, Aytan-Aktug techniques were used and evaluated on a dataset with multiple species and antibiotics, covering 54 combinations of one of nine species and one of 20 antibiotics with homology-based data stratification (**Supplemental Methods**, **Supplemental Fig. S25**). We did not find a significant difference between the homology-aware CV evaluation of Aytan-Aktug single-species models and a control multi-species model relying on species-specific reference databases for detecting mutations (paired *t*-test; *p*-value 0.47). However, the second type of Aytan-Aktug cross-species model that relied on a combined reference database for detecting mutations performed less well than the Aytan-Aktug single-species models (paired *t*-test; *p =* 1.15×10^-4^).

To assess the multi-species model on a species that was not included in the training set, we trained and tested cross-species models via the Aytan-Aktug technique (multi-antibiotic model), Kover (single-antibiotic models), and PhenotypeSeeker (single-antibiotic models), using LOSO evaluation, on the abovementioned dataset with multiple species and antibiotics (**Supplemental Methods**). For all the LOSO cross-species models, we found each performed significantly less well than the corresponding method’s single-species models (**Fig. 5, Supplemental Fig. S24**), with paired *t*-test *p*-values of 1.38× 10^-13^, 7.17×10^-19^, 3.42×10^-12^, and 4.01×10^-18^ for Aytan-Aktug, Kover set covering machines, Kover classification and regression trees, and PhenotypeSeeker, respectively. This agrees with a previous study [55], which tested this approach using only the Aytan-Aktug technique for only one species.

**Figure 5:**
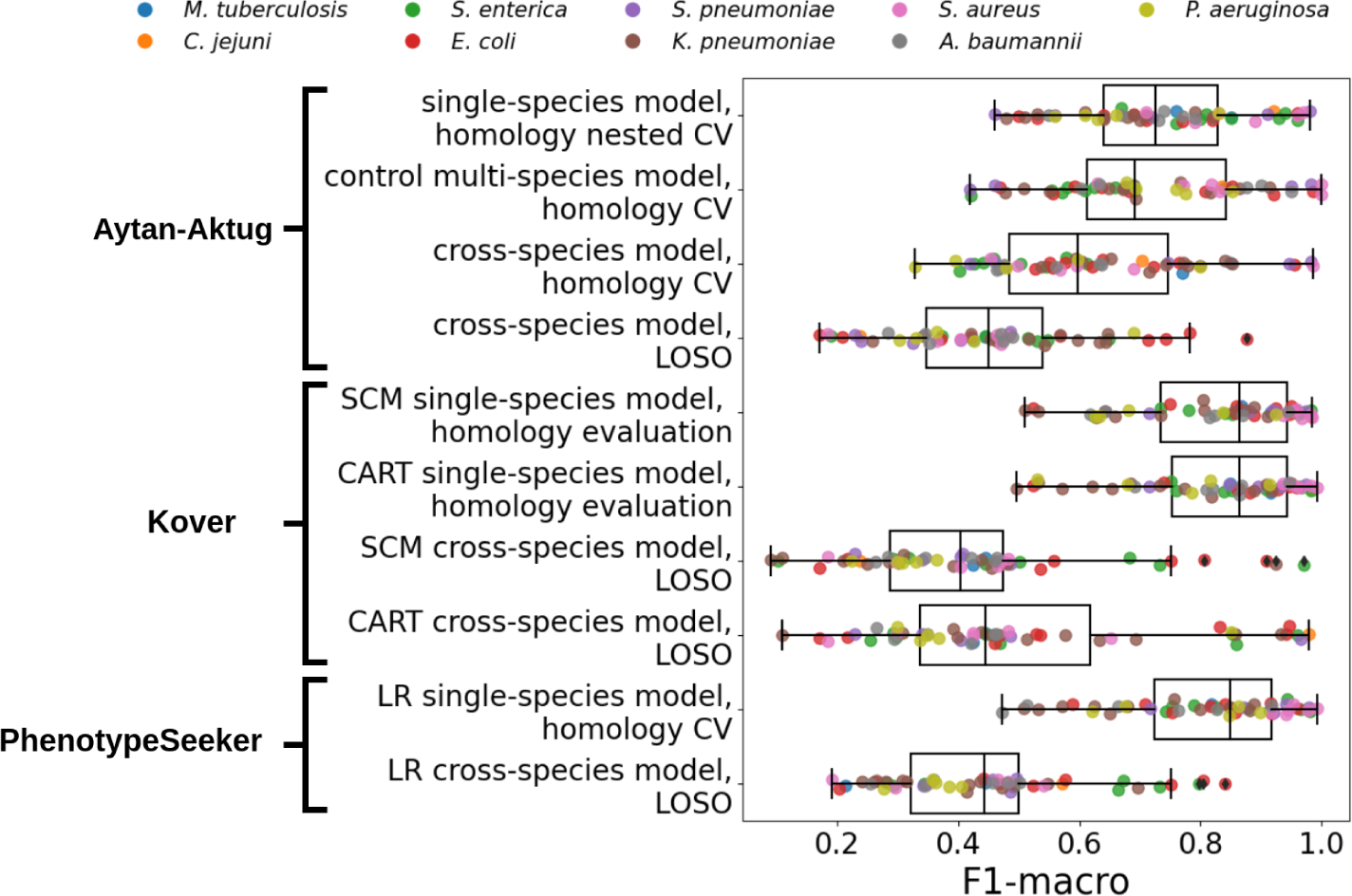
Comparison of the performance (F1-macro) of multi-species models and single-species models. Performance (F1-macro) of LOSO evaluations of cross-species models using Aytan-Aktug, Kover set covering machines (SCM), Kover classification and regression trees (CART), and PhenotypeSeeker logistic regression (LR), as well as a homology-aware evaluation of the control multi-species model (Aytan-Aktug) compared with the performance (F1-macro mean) of the corresponding single-species models (Aytan-Aktug, Kover, and PhenotypeSeeker). The evaluations were conducted on the dataset with multiple species and antibiotics, covering nine species (indicated by the color).

Compared with all the single models evaluated with homology-aware folds (**Fig. 1A**), at least one multi-factor model (**Fig. 1B**) performed best or tied first for 13 species–antibiotic combinations according to the F1-macro metric (**Supplemental Table S8A**; for the results of multi-factor models for all possible species–antibiotic combinations, see **Table S8B–J**). Notably, the Aytan-Aktug control multi-species model achieved significantly better performance on *A. baumannii*–trimethoprim/sulfamethoxazole, outperforming the second-best method (the Aytan-Aktug model with single species and antibiotics), with a higher F1-macro by 0.18.

### A guide to selecting software for specific species–antibiotic combinations

To help users in selecting the most suitable software for their analysis, we compiled a table of recommendations derived from our evaluations across the different scenarios (**Fig. 1A**), where we selected the software with the highest F1-macro mean, resorting to the lowest standard deviation for a tie, across the different folds for each species–antibiotic combination (**Supplemental Table S6**). This table reflected the upper limit of the accuracy of AMR predictions, which was achieved by selectively using the best-performing software for each species–antibiotic combination (the outer “envelopes” of the F1-macro mean circles in **Supplemental Fig. S4**). In the random evaluations, this approach allowed the prediction of AMR phenotypes with an F1-macro of ≥0.9 for 79% of the species–antibiotic combinations. Predictions for evolutionarily divergent strains declined to 60% in phylogeny-aware evaluations and to 55% in homology-aware evaluations. In 49% of the species–antibiotic combinations, the software recommendations contained at least one consistent method across different evaluation settings, indicative of the robust behavior of the corresponding models in cases of varying evolutionary divergences between the training and test genomes’ data. In the results of evaluating the “multi-” models (**Fig. 1B**), the F1-macro of the Aytan-Aktug control multi-species model was substantially higher by 0.18 compared with the second-best method for the combination of *A. baumannii* and trimethoprim/sulfamethoxazole. This led to the model’s inclusion in the selection guidelines (**Supplemental Table S6**) for that antibiotic–species pairing. It should be noted that different data splits and evaluation approaches were used because of the “multi-” configuration of the models (as indicated in **Fig. 1A,B**).

To facilitate the use of the recommended software, we provided an integrated automated pipeline for predicting AMR phenotypes based on the top three models (Kover, PhenotypeSeeker, and ResFinder) (**Table 1**) after training the ML models on each of the 78 datasets (**Fig. 2A**, https://github.com/hzi-bifo/AMR_prediction_pipeline). The models were trained using the set of hyperparameters selected via CV under the three above mentioned scenarios (**Supplemental Methods**). For a particular species–antibiotic combination, in the case when the recommendations differ between scenarios, users need to determine the degree of evolutionary divergence between the training genomes and the genomes they are analyzing. If the genome is evolutionarily close to these training genomes, the best choice would be to use the method that performed best in random evaluations; otherwise, the method performing best in the other settings should be chosen.

## Conclusions

Addressing the lack of a systematic comparison of ML-based AMR prediction methods, our study provides a broad and nuanced evaluation of five AMR prediction methods, including both ML-based techniques and the rule-based AMR catalog mapping approach. Our analysis revealed the capabilities of leading computational methods for AMR phenotyping across species and antibiotics in different scenarios, guiding selection of the software for specific species–antibiotic pairs. To date, no studies have benchmarked ML-based AMR software across such a broad spectrum.

When comparing the ML-based and rule-based catalog mapping methods, a notable discrepancy in performance was observed between the two. The ML methods’ superiority under random folds underscores the strength of ML techniques in identifying unknown or novel AMR determinants, a crucial capability, given the lack of comprehensive AMR catalogs for numerous species–antibiotic combinations. The rule-based approach’s superiority under more challenging scenarios highlights its advantage of being generalized to evolutionarily divergent genomes when AMR catalogs are available, which was the case for 83% of all species–antibiotic combinations evaluated. Discrepancies in performance between the two approaches have also been reported by others [48,61–63,65].

This tradeoff is demonstrated by Kover, which underperformed in the more complex scenarios compared with the rule-based method but consistently outperformed the other ML-based methods in all scenarios. Kover’s superiority may be attributable to procedures specially designed to address the issues of generalizing learning by mitigating dimensionality problems in ML through rule-based ML models based on compression theory. The Aytan-Aktug method might be limited by the AMR catalog used to extract the features, as its developers have noted no significant improvement over the results obtained by ResFinder [21], and Seq2Geno2Pheno’s performance might be weakened by the lack of information of core gene mutations in its features, since it provides only the reference strain for detecting SNPs in *P. aeruginosa* genomes.

For individual species—antibiotic combinations, the performance of ML methods generally declined for more challenging evaluation scenarios, whereas ResFinder maintained consistent performance. This trend can potentially be explained by three hypotheses pertaining to the role of the background lineage in ML models [49,61,68]. The first is shortcut learning. As homologous isolates from a specific source can predominantly exhibit one AMR phenotype caused by factors such as the environment [82], models learning AMR patterns may pick up spurious clade-specific signals from the dataset. This hypothesis proposes the incorporation of information on gene expression for AMR phenotyping. The second hypothesis posits the independent stepwise evolution of strains leading to different AMR variants, exemplified by the emergence of multidrug-resistant *E. coli* ST131 [87,88] and carbapenem-resistant *K. pneumoniae* ST258 [89] clones. This indicates the importance of having comprehensive training data. Although evolution can be both contingent and deterministic [90], an ML-based investigation [91] aligns with the deterministic theory, and some methods [17,44] have achieved moderately accurate AMR phenotyping using phylogenetic information, indicating the potential efficacy of exclusively using ancestral lineages for training ML models in the absence of comprehensive data. Lastly, the third hypothesis suggests that the combined effects of AMR variants and background lineages could impact the efficacy of ML-derived predictors, as indicated by the frequency of misclassified genomes. Our misclassification analysis offers a novel insight by uncovering clade-specific variations in the accuracy of AMR predictions, shedding some light over the complex interactions between an organism’s phylogeny and the AMR phenotype. Our approach complements existing genome-wide association [92] and ML [68] studies.

Among the 14 benchmarked antibiotic classes, predictions for macrolides and sulfonamides yielded the highest accuracies, whereas the performance for beta-lactams exhibited high variation, even with random folds. This suggests that it is more difficult for ML algorithms to capture the multifaceted resistance to beta-lactams, including altered penicillin-binding proteins, beta-lactamase production, and changes in porin channels [2]. We evaluated the performance of both multi-antibiotic and multi-species models, finding either no significant improvement or, in some cases, a decrease in their effectiveness compared with single-species and single-antibiotic models.

Unlike most studies in this domain, our workflow incorporates more sophisticated and less biased evaluation techniques and distinct sample partitioning methods. For assessing ML classifiers, studies commonly rely on repeated CV [45,61,63] or nested CV [41,52], where, for the latter, hyperparameters are optimized on the training fold using an inner CV loop. We used nested CV for the ML methods, as it offers more rigorous and unbiased performance estimates. For the rule-based method ResFinder, where hyperparameter optimization is not needed, we evaluated the performance on the same folds as for the other methods, referring to this as iterative evaluation. Similarly, we applied CV for evaluating Kover. Kover uses an intrinsic hyperparameter optimization, which, in combination with CV, we refer to as “iterative bound selection”; here, at each iteration, the classifier maximizing the “F1-macro” on the training set was selected from the two classifiers produced by Kover for each dataset. For generating CV folds, previous ML-based AMR studies mostly partitioned the samples randomly, with a few using phylogeny-aware and random folds [43], homology-aware folds [55], and country-based folds [50]. Uniquely, our research incorporates three distinct sample partitioning methods, aiming to evaluate the generalization power and robustness of ML-based software when confronted with taxa that differ in their evolutionary history from the training data.

Our clinically focused analysis indicated that the tested methods provide very accurate predictions of antimicrobial susceptibility for many genomes with representative training data available, but require further optimization for others, especially for evolutionarily divergent strains. Further improvements could arise from including (1) more genome data with AMR phenotypes from the wet lab for more comprehensive and phenotype-balanced training data; (2) further optimizing the ML techniques to capture the underlying patterns associated with AMR phenotypes, for example, by using transfer learning, up-sampling and/or sub-sampling before training, and an ensemble of existing AMR bioinformatics methods.

## Materials and Methods

### Datasets

We compiled 78 datasets from the PATRIC genome database [77,78] (accessed in Dec 2020) through the following steps to ensure quality and relevance. (a) We downloaded genomes with laboratory-determined AMR phenotypes, resulting in 99 species, including 67,836 genomes. We kept species with more than 500 genomes, resulting in 13 species, including 64,738 genomes. (b) We filtered genomes using PATRIC’s quality measuring attributes [93,94], avoiding plasmid-only sequences, ensuring good genome quality, limiting contig count, and checking the fine consistency, coarse consistency, completeness, and contamination criteria (see **Supplemental Methods** for thresholds). (c) We computed the average genome length for each species. We then kept only the genomes whose length deviated from the average by no more than 10% of the average. (d) We retained genomes with either resistant or susceptible phenotypes, excluding genomes with intermediate phenotypes or without phenotypes. (e) We retained species–antibiotic combinations with a minimum of 100 genomes for both resistant and susceptible phenotypes. (f) We included 402 *P. aeruginosa* genomes, and excluded the *S. pneumoniae*–beta-lactam and *M. tuberculosis*–rifampin combinations (**Supplemental Methods,** https://github.com/hzi-bifo/AMR_benchmarking/blob/main/data/README.md).

### Sample partitioning

For phylogeny-aware folds, core gene alignments were created via Seq2Geno using Prokka [95] and a modified Roary version [43,96]; subsequently, joint neighborhood phylogenetic trees were derived from these alignments using the R packages phangorn and phytools. Folds were then generated, based on the clade structure of the phylogenetic tree for each species, with the sample partitions annotated by Geno2Pheno. Random folds were generated using Geno2Pheno, excluding *M. tuberculosis* folds, which were created using the scikit-learn package’s “model_selection.KFold” module. Homology-aware folds were produced using Aytan-Aktug techniques [55], refined by us (**Supplemental Methods**). Sample partitions indicated by PATRIC’s genome IDs (random, phylogeny-aware, and homology-aware folds) and visualized on phylogenetic trees (random and phylogeny-aware folds) are available at Mendeley Data **(doi: 10.17632/6vc2msmsxj.1)**.

### Evaluation

For PhenotypeSeeker, Seq2Geno2Pheno, Aytan-Aktug, and ML baseline, 10-fold nested CV was used. In each iteration, a fold was held out as the test set, while the remaining nine folds were used for training, with the hyperparameters and classifier chosen via a 9-fold conventional CV (**Supplemental Methods**). This procedure was iterated over the 10 folds (**Supplemental Fig. S2A**). For Kover, CV with 10-fold iterative bound selection was used. Here, in each iteration, a fold was held out as the test set, while the remaining nine folds were used for training either Kover classification and regression trees or Kover set covering machines; the hyperparameters were chosen via bound selection [38]; the classifier was selected via a nine-fold conventional CV (**Supplemental Methods**). This procedure was iterated over the 10 folds (**Supplemental Fig. S2B**). For the rule-based software, a 10-fold iterative evaluation was used. In this process, each fold was assessed sequentially by ResFinder (**Supplemental Fig. S2C**).

### Evaluation metrics

The F1-macro (**Eq. 7**) was computed using the Python scikit-learn package f1_score module. The precision-positive (**Eq. 1**), recall-positive (**Eq. 2**), F1-positive (**Eq. 3**), precision-negative (**Eq. 4**), recall-negative (**Eq. 5**), F1-negative (**Eq. 6**), and accuracy (**Eq. 8**) metrics were computed using the Python scikit-learn package classification_report module. The metrics were defined as follows:

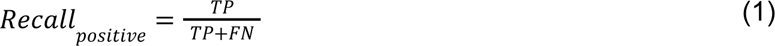

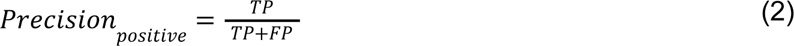

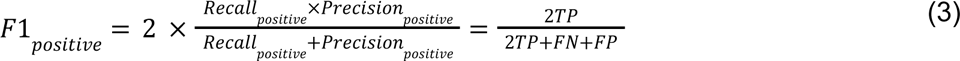

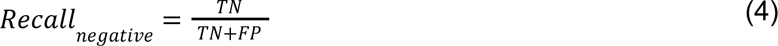

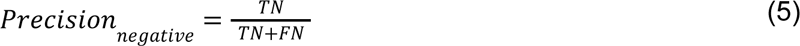

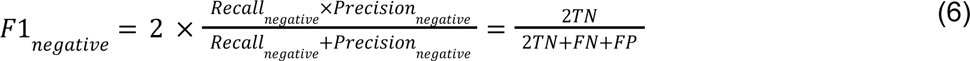

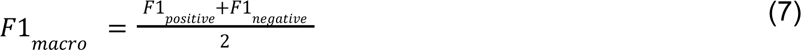

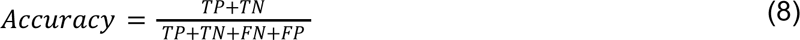

In these formulae, *TN* is the number of true negatives, *FP* is the number of false positives, *FN* is the number of false negatives, and *TP* is the number of true positives.

### Statistical analysis

In iterative evaluation approaches, the mean and standard deviation of the evaluation metrics were calculated across the 10 folds for each species–antibiotic combination and each method (**Fig. 1**). Sample-weighted means and standard deviations were used for homology-aware folds, as the distribution of samples was uneven across the folds. Arithmetic means and normal standard deviations were used for random and phylogeny-aware folds. In the conventional CV for multi-species models, we computed the metrics for the holdout test set (**Fig. 1B**). To avoid ill-defined scores resulting from the uneven distribution of negative samples across folds (Supplemental Methods), the clinically oriented F1-negative, precision-negative, and recall-negative metrics were computed after pooling the prediction results from the 10 folds. All the metric scores were rounded to two decimal points for comparison. The F1-macro mean, and the clinical-oriented F1-negative and precision-negative metrics were used for analyzing the performance. To test if the performance of the two methods was significantly different, a paired *t*-test was applied to the F1-macro means of the two methods using the Python SciPy library stats.ttest_rel module. To test if a method significantly outperformed the other method, a one-sided paired *t*-test was applied using the abovementioned Python module, with the “alternative” parameter set to “greater”.

## Supplemental material

### Supplemental Texts

- Supplemental Results
- Supplemental Methods

### Supplemental Figures

- Figure S1: Overview of the datasets.
- Figure S2: Three cross-validation approaches in the benchmarking framework.
- Figure S3: The sample partitioning methods used for cross-validation.
- Figure S4: Radar plot of the performance of the AMR prediction methods.
- Figure S5–7: Performance (F1-macro) of the methods with antibiotics shared by multiple species under the scenarios of random folds, phylogeny-aware folds, and homology-aware folds.
- Figure S8: Changes in performance (F1-macro mean) for each antibiotic class with random folds, phylogeny-aware folds, and homology-aware folds.
- Figure S9: Changes in performance (F1-macro mean) for each antibiotic with random folds, phylogeny-aware folds, and homology-aware folds.
- Figure S10: Changes in the stability of performance (standard deviation of the macro F1-score) for (A) each method and (B) each species, under the scenarios of random folds, phylogeny-aware folds, and homology-aware folds.
- Figure S11–12: Changes in performance and stability (mean and standard deviation of the negative F1-score) for each species–antibiotic combination with random folds, phylogeny-aware folds, and homology-aware folds.
- Figure S13–22: Distribution of misclassified genomes of *E. coli*, *S. enterica, S. aureus*, *K. pneumoniae*, *S. pneumoniae*, *A. baumannii*, *P. aeruginosa, C. jejuni, N. gonorrhoeae*, and *E. faecium* on the corresponding phylogenetic trees.
- Figure S23: Comparison of the performance (F1-macro mean and standard deviation) of the Aytan-Aktug single-species models with single or multiple antibiotics.
- Figure S24: Performance (F1-macro) of the multi-species models using Aytan-Aktug, Kover, and PhenotypeSeeker, in comparison with the corresponding single-species model (F1-macro mean and standard deviation).
- Figure S25: Sample partitioning method for the control Aytan-Aktug multi-species model and the cross-species model.
- Figure S26: The performance (F1-macro) of the Aytan-Aktug control multi-species model, the cross-species model, and the single-species models compared by the original Aytan-Aktug methods.

### Supplemental Tables

- **Table S1:** ML-based studies on determining AMR phenotypes on the basis of genomic data.
- **Table S2:** The antibiotics’ acronyms, classes, and mechanisms of action.
- **Table S3:** Performance (F1-macro, F1-negative, F1-positive, accuracy, precision, recall, clinical_F1-negative, clinical_precision-negative, and clinical_recall-negative) of five benchmarked methods alongside the baseline method with respect to random folds, phylogeny-aware folds, and homology-aware folds.
- **Table S4: A.** Fraction of experiments that performed above the ML baseline threshold, stratified by the methods of splitting the dataset. **B.** Fraction of ML-based experiments that performed above the indicated threshold, stratified by the method of splitting the dataset. **C.** Fractions of species–antibiotic combinations for which the ML-based methods and rule-based methods performed best.
- **Table S5:** Performance (F1-macro, F1-positive, F1-negative, accuracy, precision, and recall) of the KMA-based ResFinder and the BLAST-based ResFinder on each species—antibiotic combination’s whole dataset (instead of iteratively on folds).
- **Table S6:** Three software recommendation lists, corresponding to the random, phylogeny-aware, and homology-aware evaluations. For each species–antibiotic combination, we selected the methods (or several of them) with the highest F1-macro mean and, in the case of a tie, the lowest F1-macro standard deviation. The lists can guide researchers in selecting software for each of the 78 species–antibiotic combinations.
- **Table S7:** Performance (F1-macro, F1-positive, F1-negative, accuracy, precision, and recall) of the Aytan-Aktug model for single species and antibiotics using the default hyperparameters, in comparison with our modified version.
- **Table S8: A.** The 13 species–antibiotic combinations for which at least one multi-factor model performed best or tied first with the single-factor models (F1-macro metric). **B–J.** Performance (F1-macro, F1-positive, F1-negative, accuracy, precision, and recall) of the Aytan-Aktug single-species multi-antibiotic model, the control multi-species model, and the cross-species model, as well as the LOSO cross-species models for Aytan-Aktug, Kover and PhenotypeSeeker.
- **Table S9:** Benchmarking results of 14 datasets spanning *C. jejuni* and *E. coli* in two repeated independent evaluations via our workflow.
- **Table S10:** Mean, standard deviation, and median of the F1-macro mean for each benchmarked method.
- **Table S11:** Dataset folds potentially suffering from ill-defined F1-negative and F1-positive metrics in the phylogeny-aware and homology-aware evaluations.

### Data and code availability

The benchmarking datasets were based on the open-source PATRIC (https://patricbrc.org/) and are available at **Mendeley Data (doi: 10.17632/6vc2msmsxj.1)**. The source code used for benchmarking the AMR software and software adaptations is available on GitHub (https://github.com/hzi-bifo/AMR_benchmarking_khu). Tutorials for evaluating ML-based AMR software are provided (https://github.com/hzi-bifo/AMR_benchmarking/wiki/AMR-phenotyping-benchmarking-tutorial).

### Competing interests

E.A. and T.K. are two of the co-authors of the software package Seq2Geno2Pheno [43].

## Acknowledgements

P.C.M. received funding from the Cluster of Excellence RESIST (EXC 2155) funded by Deutsche Forschungsgemeinschaft (DFG) (project number 390874280). This work was funded in part by the German Center for Infection Research (DZIF) Translational Infrastructure Bioresources, Biodata, and Digital Health (TI BBD) (project number TI 12.002) and the DFG (project number 460129525; NFDI4Microbiota).

## Key Points

- We introduce a new benchmarking standard using advanced, unbiased evaluation techniques to rigorously test the performance and generalizability of AMR prediction models.
- We performed benchmarking by training and testing four ML methods and a rule-based method on 78 species–antibiotic datasets.
- Benchmarking revealed that although ML methods excelled with closely related bacterial strains, the rule-based ResFinder was superior for divergent genomes, with Kover emerging as the leading ML tool in terms of performance.
- The study provides explicit software recommendations for predicting AMR phenotypes, aiding in the accurate detection of resistance across diverse bacterial species.
- The findings emphasize the need for refined prediction models, especially for clinically critical antibiotics such as beta-lactams, to enhance their utility in real-world applications.

## Author Biographies

**Kaixin Hu** is a PhD student at the Department Computational Biology of Infection Research of the Helmholtz Centre for Infection Research. She focuses on machine learning and antimicrobial resistance.

**Fernando Meyer** is a postdoctoral researcher at the Department Computational Biology of Infection Research of the Helmholtz Centre for Infection Research. He focuses on metagenomics.

**Zhi-Luo Deng** is a postdoctoral researcher at the Department Computational Biology of Infection Research of the Helmholtz Centre for Infection Research. He focuses on microbial genomics and microbiome.

**Ehsaneddin Asgari** is a postdoctoral researcher at the Department Computational Biology of Infection Research of the Helmholtz Centre for Infection Research. He focuses on natural language processing and bioinformatics.

**Tzu-Hao Kuo** was a PhD student at the Department Computational Biology of Infection Research of the Helmholtz Centre for Infection Research. He is currently a research associate at Evotec International GmbH.

**Philipp C. Münch** is a research scientist at the Department Computational Biology of Infection Research of the Helmholtz Centre for Infection Research. He focuses on deep learning and computational biology.

**Alice C. McHardy** is the head of Department Computational Biology of Infection Research of the Helmholtz Centre for Infection Research. The department develops and assesses computational methods for the analysis of the microbiome, viral and bacterial pathogens from large-scale biological and epidemiological datasets.

## Notes

https://github.com/hzi-bifo/AMR_benchmarking

https://github.com/hzi-bifo/AMR_prediction_pipeline

### Summary of Updates

Main manuscript revised; Supplemental files updated.

