## Supplemental Texts for "Assessing computational predictions of antimicrobial resistance phenotypes from microbial genomes"

### Supplemental Results

#### Comparative assessment of AMR phenotyping software

In random split evaluation, all benchmarked methods demonstrated strong performance, with median F1-macro scores ranging from 0.91 (Aytan-Aktug) to 0.93 (Kover). Kover also excelled in predicting AMR for evolutionarily divergent strains, with a median F1-macro of 0.89 and 0.87 in phylogeny-based and homology-based evaluation, respectively. ResFinder had a median F1-macro of 0.84 and 0.85 in these settings (**Fig. 4A**, **Supplemental Table S10**).

When considering the application to clinical diagnostics and treatment, predicting a pathogen’s susceptibility to a specific drug is of high importance [[1]](https://paperpile.com/c/6VFvZ6/ZdztR). Similar to evaluations measured by the F1-macro mean, with the metrics for the negative class, ML methods performed better in evaluations with random splits, whereas in the phylogeny-based and homology-based evaluation settings, AMR catalog mapping with ResFinder performed comparably well. Specifically, ML methods achieved a precision in antibiotic susceptibility prediction at least as high as that of ResFinder in 62% (194/312) of random evaluation experiments, and in 49% (131/268) and 42% (132/312) of phylogeny-aware and homology-aware evaluation, respectively. The F1-score for ML-based methods in antibiotic susceptibility prediction was at least as high for 69% (214/312), 54% (145/268), and 47% (147/312) of the experiments with random, phylogeny-based, and homology-based evaluation, respectively.

#### Misclassification in AMR phenotyping

To further investigate the influence of genetic ancestry or evolutionary history on model accuracy, for each species benchmarked in phylogeny-based scenarios, we quantified the relationship between the background lineage of each genome and its frequency of being correctly predicted with random folds, but misclassified with phylogeny-aware or homology-aware folds across all the antibiotic combinations and four ML-based AMR prediction methods (**Supplemental Methods**). We observed an uneven distribution of the misclassification frequencies across the phylogenetic tree for all ten species, *E. coli*, *S. enterica, S. aureus*, *K. pneumoniae*, *S. pneumoniae*, *A. baumannii*, *P. aeruginosa, C. jejuni, N. gonorrhoeae* and *E. faecium* (**Supplemental Methods**, **Supplemental Fig. S13-S22**). Specifically, a small number of genomes spreading across various clades exhibited relatively high misclassification frequencies, whereas the majority, even those sharing the same clades as genomes frequently misclassified, presented low misclassification frequencies. For all the eight species (except for *C. jejuni* and *E. faecium* where an antibiotic’s phenotype was annotated for each genome), despite the varied number of antibiotics for which each genome's phenotypes were annotated, we observed no clear association of the misclassification frequency pulse and the number of antimicrobial resistances(**Supplemental Fig. S13-S22**). This indicated that AMR phenotyping across a range of drugs might be complicated by the background lineage.

#### Evaluating the software reproducibility

Computational reproducibility is a key property in software evaluation. As we evaluated software iteratively ten times (**Fig. 1A**), reporting a mean and standard deviation of metrics, our benchmarking study largely reduced the impact of stochasticity in software on evaluation. Of the five AMR methods, we observed deterministic behavior by ResFinder on all the datasets, as expected. To explore the reproducibility of ML methods, after our native benchmarking was finished, we set up our benchmarking at a new address on our computing server and re-evaluated the four ML methods (via the workflow outlined in **Fig. 1A**) on 14 datasets for two randomly selected species, *C. jejuni* and *E. coli*, using the phylogeny-based evaluation. This provided us with another mean F1-macro value rounded to 2 decimals for each method and each dataset (**Supplemental Table S9**).

PhenotypeSeeker produced deterministic evaluation results; for Kover, two out of 14 datasets had varied mean F1-macro values compared with our first evaluation, with a difference between 0.02 and 0.04; for Aytan-Aktug, six out of 14 datasets had varied mean F1-macro values, with a difference between 0.01 and 0.05, which was attributable to stochasticity in neural networks weight initialization and dropout mechanism (**Materials and Methods**); for Seq2Geno2Pheno, eleven out of 14 datasets had varied mean F1-macro scores, with a difference between 0.01 and 0.05, which was attributable to stochasticity in the Roary [[2]](https://paperpile.com/c/6VFvZ6/kHTiu) pan-genomes analysis. Therefore, computational analyses with the evaluated ML models are not fully deterministic, with the exception of PhenotypeSeeker.

#### Assessment of multi-antibiotic and multi-species AMR prediction models

We examined the performance of both multi-antibiotic and multi-species models. We found no significant difference between multi-antibiotic and single-antibiotic models, and also no significant difference between the control multi-species model and single-species model. The multi-antibiotic model (also known as multi-label/output classification) has been investigated in several AMR phenotype prediction studies on *M. tuberculosis* [[102,113,117,119]](https://paperpile.com/c/3bYSnr/7u8m8+fhu7q+rCNVb+55vFt). Aytan-Aktug et al. [[109]](https://paperpile.com/c/3bYSnr/KRdUH) have investigated both the multi-antibiotic model and multi-species models. Whereas, all these methods used either random forest or neural networks. We extended Aytan-Aktug et al. [[109]](https://paperpile.com/c/3bYSnr/KRdUH) research by first refining their neural networks training procedure and then evaluating both models on many more species–antibiotic combinations, and evaluating the cross-species model (evaluation of multi-species model by LOSO) using more techniques. Notably, in general, cross-species models exhibited inferior performance compared to single-species models. The LOSO evaluation of cross-species models rigorously tested the software in challenging metagenomic scenarios, when the pathogen sequences can not be reliably assigned to a specific taxon or are assigned to a less-studied taxon lacking high-quality catalogs and trained ML models. This LOSO evaluation approach could push the boundaries of AMR prediction models, driving advancements in methodological robustness and adaptability for more translational and clinical applications. Our LOSO evaluation results indicated that the inherent variability in resistance patterns across different organisms hindered the direct application of state-of-the-art ML-based AMR prediction methods, which are designed for single isolates, on metagenomic data.

### Supplemental Methods

#### Datasets

We used the PATRIC genome database [[8,9]](https://paperpile.com/c/6VFvZ6/pusdH+XhJKJ) (accessed in Dec 2020), which enables easy access to genomes and their metadata through the command line. We provided the codes for retrieving and preprocessing data through phenotype availability control and genome quality control. (a) We downloaded all the genomes and corresponding metadata that were with available laboratory-determined AMR phenotypes. This resulted in 99 species, including 67,836 genomes. We retained species with more than 500 genomes, resulting in 13 species, including 64,738 genomes. (b) We applied a set of criteria for genome quality control using PATRIC’s quality measuring attributes: (1) sequence data is not plasmid-only; (2) genome quality (provided by PATRIC) is Good; (3) contig count is limited to the greater of either 100 or 0.75 quantiles of the contig count across all genomes of the same specie; (4) fine consistency (provided by PATRIC) higher than 97%; (5) coarse consistency (provided by PATRIC) higher than 98%; (6) completeness (provided by PATRIC) higher than 98% and contamination (provided by PATRIC) lower than 2%, or one of them is null value with the other one meets the criteria. (c) For each species, we computed the mean genome length of the selected genomes from step (b), we then retained the genomes with lengths within the range of one-twentieth of the calculated mean from the calculated mean. (d) We retained the genomes with either R (resistant) or S (susceptible) phenotypes, and excluded genomes with intermediate phenotypes or without phenotypes. (e) We retained species–antibiotic combinations with at least 100 genomes for both R and S phenotype classes. (f) Apart from the procedures mentioned above, we made additional appends and filters: we included 402 *P. aeruginosa* genomes sourced from Seq2Geno2Pheno [[10]](https://paperpile.com/c/6VFvZ6/Hx67c), which were filtered out in the above-mentioned procedures due to the lack of completeness or contamination indicators in PATRIC; we excluded the *S. pneumoniae-* beta-lactam combination as beta-lactam represents a class of antibiotics; we excluded the *M. tuberculosis*–rifampin combination as we included in *M. tuberculosis*–*r*ifampicin combination.

Our procedures resulted in 78 datasets, each representing a combination of species and antibiotics among 11 species (*E. coli, S. aureus, K. pneumoniae, S. pneumoniae, A. baumannii, P. aeruginosa, S. enterica, E. faecium, C. jejuni, N. gonorrhoeae,* and *M. tuberculosis*), and 44 antibiotics, totaling 31,195 genomes (**Fig. 2A**, Mendeley Data).

For evaluating the Aytan-Aktug single-species multi-antibiotic model (**Fig. 1B**), we further merged the 78 datasets that shared the same species, resulting in 9 datasets each corresponding to 1 of 9 species (except for *E. faecium* and *C. jejuni* of the above-mentioned 11 species), all-together containing 76 species–antibiotic combinations (**Fig. 2B**). And to evaluate multi-species models (**Fig. 1B**), we merged the 78 datasets to a multi-species–antibiotic dataset covering all antibiotics that have been associated with at least two species, containing 54 species–antibiotic combinations of 9 species (except for *N. gonorrhoeae* and *C. jejuni* of the above-mentioned 11 species) and 20 antibiotics (**Fig. 2C**). The PATRIC identities of samples in these 10 datasets were provided in Mendeley Data.

Of note, only two studies have validated their methods on more than five pathogen species: Drouin A et al. benchmark their novel algorithms on twelve species against various ML classifiers, which are based on k-mers feature representation [[11]](https://paperpile.com/c/6VFvZ6/dVXxb); Kim J et al. evaluate their software on nine species but only compared it to another software for one species, without re-training the external software on the same dataset [[12]](https://paperpile.com/c/6VFvZ6/RmDni).

#### Sample partitioning

In our benchmarking workflow A (**Fig. 1A**), all methods were evaluated with 3 sets of folds. Phylogeny-aware folds were proposed by Seq2Geno2Pheno [[10]](https://paperpile.com/c/6VFvZ6/Hx67c), in which they split the samples based on a phylogenetic tree generated through mapping to a *P. aeruginosa* reference strain. As we were benchmarking with 11 species instead of only *P. aeruginosa*, we modified the pipeline of Seq2Geno to generate phylogenetic trees based on core gene alignment. Specifically, we generated core gene alignments through the pipeline of Seq2Geno, in which Prokka [[13]](https://paperpile.com/c/6VFvZ6/6u2nx) and a modified version of Roary [[2,10]](https://paperpile.com/c/6VFvZ6/kHTiu+Hx67c) were used, and then we inferred joint neighborhood phylogenetic trees from the core gene alignment using the R package of phangorn and phytools. Finally, we used Geno2Pheno to generate the folds split based on the clade structure of the phylogenetic tree for each species, accompanied by visualization of the tree annotated with the sample partition (**Supplemental Fig. S3 B, Mendeley Data**). Seq2Geno2Pheno generates folds of equal size, which facilitate evaluation in terms of statistical reliability, but the downside is that some phylogenetically close strains may be allocated into both the training set and the test set, depending on the tree structure. Besides, of the 11 species in benchmarking, Seq2Geno failed to generate a phylogenetic tree for *M. tuberculosis* within the two-month timeframe, resulting in phylogeny-aware folds for 67 datasets spanning 10 species.

Random folds (except for *M. tuberculosis* folds) were generated using Geno2Pheno [[10]](https://paperpile.com/c/6VFvZ6/Hx67c). For each species–antibiotic combination, we annotated the corresponding phylogenetic tree with the random split of samples (**Supplemental Fig. S3 D, Mendeley Data**). *M. tuberculosis* folds were generated via the scikit-learn package model_selection.KFold module.

Homology-aware folds were proposed by Aytan-Aktug [[7]](https://paperpile.com/c/6VFvZ6/pEK4B), in which for each dataset they first use KMA [[14]](https://paperpile.com/c/6VFvZ6/OscrE) to calculate similarities between samples and cluster them into clusters, and then further group the clusters into 10 folds for evaluation (**Supplemental Fig. S3 A**). For clustering samples of *M. tuberculosis* (amikacin, ethambutol, ethiomide, ethionamide, kanamycin, ofloxacin), the arguments of (-k 16; -ht 0.99; -hq 0.99) were used; for N. gonorrhoeae(2 antibiotics) and *M. tuberculosis* (capreomycin, isoniazid, pyrazinamide, rifampicin, streptomycin), the arguments of (-k 16; -ht 0.98; -hq 0.98) were used; for the other datasets, the arguments of (-k 16; -ht 0.9; -hq 0.9) were used. The -ht and -hq parameters corresponded to the template and query coverage thresholds, respectively, and the -k parameter corresponded to the k-mer space. We optimized the Aytan-Aktug native procedure of allocating clusters into folds by re-allocating clusters iteratively if the ratio of sample size of a fold over the sample size of the dataset was less than a predefined threshold, mainly because their methods would result in a fold with no samples for some *M. tuberculosis*-antibiotic combinations, and our modification also made the sample distribution across the 10 folds slightly more balanced for the other combinations. The PATRIC IDs of samples in each fold in the above-mentioned 3 sets of folds are available in JSON format (**Supplemental File 1**).

Our three sample partitioning methods provide a comprehensive assessment, thoroughly examining the generalization capacity across a wide range of scenarios. First, random partitioning was utilized, where samples were divided into ten nearly equal-sized folds [[10]](https://paperpile.com/c/6VFvZ6/Hx67c). Although easy to implement, this approach falls short in capturing many real-world complexities. It lacks consideration for genome phylogenetic relationships or genetic similarities, potentially leading to a situation where a substantial fraction of genomes in the test set are highly similar to those in the training set. Consequently, the evaluation of the model's phylogenetic generalization capability to strains divergent from training strains is limited. Thus, second, a phylogeny-based method was applied, by computing a phylogenetic tree for the input genomes, after which samples were partitioned into ten folds based on the clade structure of the tree [[10]](https://paperpile.com/c/6VFvZ6/Hx67c). To balance realistic evaluation and statistical reliability, this approach generated nearly equal-sized folds by allocating a few phylogenetically close strains into separate folds, leading to the possibility of having a few highly similar genomes in the training and test sets. Third, a homology-based approach was adopted, where samples were split into 10 folds using sequence similarity-based clusters determined with the KMA software, as described in Aytan-Aktug et al. [[7]](https://paperpile.com/c/6VFvZ6/pEK4B). This approach, characterized by the largest degree of divergence between training and test genomes, represents the most challenging evaluation scenario, followed by phylogenetic stratification and then random evaluation.

In the workflow B (**Fig. 1B**), the Aytan-Aktug multi-antibiotic model and multi-species–antibiotic model were evaluated with homology-aware folds. To establish CV folds of 9 multi-antibiotic datasets for the single-species multi-antibiotic model, the KMA arguments of (-k 16; -ht 0.98; -hq 0.98) were used on *N. gonorrhoeae* and (-k 16; -ht 0.99; -hq 0.99) on *M. tuberculosis* samples to cluster them; the KMA arguments of (-k 16; -ht 0.9; -hq 0.9) were used on each of other species’ datasets to cluster corresponding samples. Then, for each of the 9 datasets, we grouped the clusters into 10 folds the same way as for a dataset of individual species–antibiotic combinations described above. To establish folds of a multi-species–antibiotic dataset for the control-multi-species model and the cross-species multi-species model, the KMA arguments of (-k 16; -ht 0.99; -hq 0.99) were used on *M. tuberculosis* samples to cluster them; the KMA arguments of (-k 16; -ht 0.9; -hq 0.9) were used on samples of the other species to cluster them for 8 times, each time on samples of the same species. Then for each species, we grouped the corresponding KMA clusters into 6 folds, resulting in 54 folds (9 species $\times$6 folds) belonging to 9 fold sets (9 species). Then we picked a fold from the fold set belonging to each of the 9 species to form a new fold, consisting of samples of 9 species. We iterated the picking procedure 6 times, in a predefined order for picking the folds in each species’ fold set, resulting in a set of 6 folds, each consisting of samples of 9 species (**Supplemental Fig. S25**). To establish folds for the cross-species leave-one-species-out model, the KMA arguments of (-k 16; -ht 0.99; -hq 0.99) were used on *M. tuberculosis* samples to cluster them; the KMA arguments of (-k 16; -ht 0.9; -hq 0.9) were used on samples of the other species to cluster them for 8 times, each time on samples of the same species. Then the same procedures as for the control multi-species model were used to generate the final 5 folds, each composed of samples of 9 species. The PATRIC IDs of samples in each fold for the above-mentioned 9 multi-antibiotic datasets and a multi-species–antibiotic dataset were provided in JSON format (**Supplemental File 1**).

#### Evaluation methods for multi-species models

Considering the extremely intensive computing involved in both nested CV and multi-species neural networks model, and the neural networks evaluation practices, the conventional CV with a holdout test was applied to the three Aytan-Aktug multi-species models (**Fig. 1B**). Specifically, the control multi-species model and the cross-species multi-species model were evaluated with a holdout test set, which was a predefined fold in homology-aware folds. The optimal hyperparameters were selected via 5-fold conventional CV under the rest of homology-aware folds.

We further used the LOSO evaluation. The Aytan-Aktug cross-species model, which is also a multi-label model, was evaluated 9 times, each time with a different holdout test set consisting of all the genomes from one of 9 species in the multi-species–antibiotic dataset. Specifically, at each iteration, all the samples from one species were grouped into the test set, and all the rest samples from the other 8 species were grouped into the training set, then a homology-aware 5-fold conventional CV was performed on the training set to select the optimal hyperparameter set, and finally the model with the selected hyperparameters was trained on the whole training set and tested on the test set. The PhenotypeSeeker cross-species model was evaluated 54 times, each time with a different holdout test set consisting of all the genomes from one species–antibiotic combination in the multi-species–antibiotic dataset. At each iteration, all the samples from that species–antibiotic combination were grouped into the test set, and all the samples annotated with the antibiotic’s phenotype but belonging to another species were grouped into the training set, then a homology-aware 10-fold conventional CV was performed on the training set to select the optimal hyperparameter set, and finally, the model with the selected hyperparameters was trained on the whole training set and tested on the test set. Kover cross-species models were evaluated 54 times, each time with the same holdout test set and training set as the PhenotypeSeeker cross-species model, but no CV was needed as bound selection was applied alongside training.

#### F1-macro as the main metric for performance analysis

F1-macro was chosen as the main metric for software performance comparison for two reasons. First, whereas false negative predictions will increase mortality [[15]](https://paperpile.com/c/6VFvZ6/Z8Kr8), false positive predictions may lead to unnecessary prescription of antibiotics, with more side effects or prolonged treatment [[16]](https://paperpile.com/c/6VFvZ6/m2A59). This resulted in two candidates: F1 scores and accuracy. However, accuracy tends to bias the estimation of performance on an imbalanced dataset, making it difficult to interpret. For example, ML algorithms tend to perform poorly on the smaller class if no specialized techniques are applied, in which case, the accuracy will overestimate the performance. Therefore, we used F1-macro, the arithmetic mean of the per-class F1 scores, which are the averages of the harmonic mean of precision and recall for samples with susceptible phenotypes and samples with resistant phenotypes, respectively. It offers an estimate of performance across both resistant and susceptible classes, independent of the class sizes. Besides, F1-negative and F1-positive scores were also provided for reference (**Supplemental Table S3**), although there were "ill-defined" cases due to the occurrence of folds with only one phenotype class in homology-aware and phylogeny-aware folds, thus cannot be used as fair metrics. When samples in the test set all had resistant phenotypes, and the evaluation, for example, resulted in a confusion matrix of (tn=0, fp=0, fn=0, tp=20), F1-negative was 0 by definition. However, this score did not accurately reflect the model's performance in predicting susceptible genomes. The same issue applied to the F1-positive score when all samples in the test set had susceptible phenotypes. On the other hand, we couldn't exclude test sets with the same phenotype when calculating per-class F1 scores, as this would also lead to the exclusion of valuable information. For instance, a test set with only resistant samples and a confusion matrix of (tn=0, fp=0, fn=1, tp=19) contained susceptible-related information and was not considered “ill-defined”. In both above-mentioned "ill-defined" cases, F1-macro was corrected to 1 instead of 0.5 using the Python scikit-learn package f1_score module, so we deemed F1-macro as the optimal evaluation metric for our benchmarking study. In homology-aware evaluation, 3.97% (31/780) folds would potentially suffer from ill-defined F1-positive, and 0.38% (3/780) folds would potentially suffer from ill-defined F1-negative; in phylogeny-aware evaluation, these percentages were 1.05% (8/760) and 1.71% (13/760), respectively (see **Table S11**). Clinical-oriented scores F1-negative, precision-negative, and recall-negative were computed after pooling prediction results from 10 folds (via 10 iterations), to avoid ill-defined scores resulting from the uneven distribution of negative samples across folds, especially for phylogeny-aware and homology-aware partition. To ensure consistency, we calculated the clinical-oriented scores the same way for random folds. The classification_report module from the Python scikit-learn package was used to calculate these clinical-oriented scores based on correct predictions and wrong predictions.

#### Software versions, revisions, parameters, and hyperparameters

To benchmark ML-based AMR phenotyping methods, we evaluated ResFinder 4.0 [[17]](https://paperpile.com/c/6VFvZ6/9o4zY) (software accessed on 2021-05-06, open-source <https://bitbucket.org/genomicepidemiology/resfinder/src/master/>. Database accessed on 2021-05-06, open-source <https://bitbucket>.org/genomicepidemiology/pointfinder_db/src/master/, https://bitbucket.org/genomicepidemiology/resfinder_db/src/master/[)](https://bitbucket.org/genomicepidemiology/resfinder/src/master/), AytanAktug [[7]](https://paperpile.com/c/6VFvZ6/pEK4B) (version: accessed on 2021-04-26, open-source <https://bitbucket.org/deaytan/data_preparation> and [https://bitbucket.org/deaytan/data_preparation)](https://bitbucket.org/deaytan/data_preparation), Seq2Geno2Pheno [[10]](https://paperpile.com/c/6VFvZ6/Hx67c) (accessed on 2021-07-11, open-source https://github.com/hzi-bifo/seq2geno[,](https://github.com/hzi-bifo/AMR_benchmarking_khu/tree/main/seq2geno-precomputed_assemblies) https://github.com/hzi-bifo/GenoPheno[)](https://galaxy.bifo.helmholtz-hzi.de/galaxy/root?tool_id=genopheno), PhenotypeSeeker [[18]](https://paperpile.com/c/6VFvZ6/olbEC) (version: 0.7.3 open source [https://github.com/bioinfo-ut/PhenotypeSeeker)](https://github.com/bioinfo-ut/PhenotypeSeeker), and Kover [[11]](https://paperpile.com/c/6VFvZ6/dVXxb) (version: 2.0, open source: [https://github.com/aldro61/kover)](https://github.com/aldro61/kover).

To evaluate software via our rigorous workflow, we made some adaptations or modifications based on the above-mentioned versions of ResFinder, Aytan-Aktug, Seq2Geno, and PhenotypeSeeker, with the adapted versions available on GitHub ([https://github.com/hzi-bifo/AMR_benchmarking_khu)](https://github.com/hzi-bifo/AMR_benchmarking_khu).

##### Adaptations to rule-based software ResFinder

ResFinder 4.0 [[17]](https://paperpile.com/c/6VFvZ6/9o4zY) predicts AMR based on mobilizable genes (previous versions of ResFinder [[19]](https://paperpile.com/c/6VFvZ6/aSLyX)) and chromosomal mutations (PointFinder [[20]](https://paperpile.com/c/6VFvZ6/3NCav)). The species-antimicrobial with mutation knowledge, and antimicrobial compounds with AMR gene knowledge can be found in the supplements of their article [[17]](https://paperpile.com/c/6VFvZ6/9o4zY). Of the 11 species included in our benchmarking, 6 species (*E. coli, S. aureus, S. enterica, E. faecium, C. jejuni, M. tuberculosis*) were equipped with chromosomal mutations database, 2 species (*Klebsiella, N. gonorrhoeae)* were equipped with in-building chromosomal mutations databases. We used the alignment method BLAST for *N. gonorrhoeae*, and KMA (1.3.15 version on 2021-05-06) for the other 10 species. The ResFinder 4.0 version only provides the KMA-based alignment option for raw sequences, not genomic sequences. We adapted it to make it available for assembled sequences as well (except for *N. gonorrhoeae*, as there were some errors in our adapting version), which made the prediction much faster. Another reason for the modification was that the Aytan-Aktug cross-species model is established based on KMA-based ResFinder [[7]](https://paperpile.com/c/6VFvZ6/pEK4B), so we followed their instructions to make the adaption of ResFinder 4.0. A paired t-test was performed to compare the F1-macro of KMA and BLAST version ResFinder evaluated on each species–antibiotic combination’s whole dataset (instead of on folds iteratively as in **Fig. 1A**, results see **Supplemental Table S5**), resulting in a paired *t*-test *p*-value of 0.64. So we can not reject the null hypothesis that there is no difference in performances between the two versions at the 0.05 significance threshold. For other parameters, we used the default settings (min_cov 0.60 and threshold 0.80).

##### Refinements and hyperparameter selection of ML-based software Aytan-Aktug techniques

For the Aytan-Aktug method, five models, including single-species-antibiotic model, single-species multi-antibiotic model, control multi-species model, cross-species model, and cross-species LOSO model, were evaluated. Features were generated using a combination of scored and binary representation methods, based on mutations and mobilizable genes detected by the above-mentioned ResFinder 4.0 [[17]](https://paperpile.com/c/6VFvZ6/9o4zY) adaptation version for 10 species, except that for *N. gonorrhoeae* the ResFinder 4.0 original version was used. The ResFinder adaption version was run with KMA alignment (1.3.15 KMA version was used except for *P. aeruginosa* samples for three multi-species models, on which the 1.4 version in September 2022 was used). For the control multi-species model and cross-species models, the two database merging methods described by the Aytan-Aktug [[7]](https://paperpile.com/c/6VFvZ6/pEK4B) were used, respectively. In Method 1, the sequence of each species was aligned only to the species-specific reference sequence. In Method 2, the sequence of each species was aligned to the sequences in a concatenated reference database. We modified the feature generation codes of multi-species models to make it easier for reproducibility and to enable a user-specified number of species and antibiotics according to combinations available instead of fixed at 5 species and 6 antibiotics for a specific dataset. Neural network classifiers were used for all Aytan-Aktug models, with a hyperparameter optimization procedure accompanied by early stopping added by us to select hyperparameters, considering the importance of hyperparameter selection for neural network training. We monitored the validation binary cross entropy loss (using torch.nn.BCELoss) every 100 epochs, and stopped the training process when the monitored loss increased twice in a row, which was implemented using a package from <https://github.com/Bjarten/early-stopping-pytorch>. We activated the early stopping mechanism after 500 epochs to speed up the training procedure, as we observed in the training process that it generally took more than 500 epochs before the loss ceased decreasing. We determined the hyperparameter tuning search range by considering both the default set given by the native article and the characters of our data, for each of the five models. After the optimization procedure, the hyperparameter set resulting in the highest F1-macro and the mean of the corresponding epoch number in the CV was used for training the training set. Besides, we optimized their neural network structure by adding the dropout hyperparameter to avoid overfitting.

For the single-species-antibiotic model, the default neural network structure of a one-layer multilayer perceptron (MLP) with 200 hidden neurons was used. We set the learning rate range as (0.001 and 0.0005) and dropout as (0 and 0.2). We set the maximum epoch number to 10,000, as we observed all single-species-antibiotic models in this study terminate before reaching the number. In each outer loop, after the optimization procedure in 8 inner loops, the hyperparameter set resulting in the highest F1-macro and the mean of the corresponding epoch number in the 8 inner loops was used for training on the corresponding outer loop training set. The average epoch number of the corresponding set of hyperparameters from the 8 inner loops was used as the epoch number parameter for training in the corresponding outer loop. A paired *t*-test on the performance (F1-macro mean) with homology-aware folds of our modified hyperparameter optimization version and the default version, resulted in the paired *t*-test *p*-value of 5.33E-06, indicating that we can reject the null hypothesis that there is no significant difference between the two versions. By F1-macro mean metric, our modified version outperformed the default version with a higher F1-macro mean (and a lower standard deviation if the mean was equal) in 64% (50/78) cases, and the same as the default version in 18% (14/78) cases (**Supplemental Table S7**).

For the single-species multi-antibiotic model, the default neural network structure of a one-layer MLP with 200 hidden neurons was used. We set the learning rate range as 0.001, and the dropout selection range as (0 and 0.2). We set the maximum epoch number to 50000, as we observed all single-species multi-antibiotic models in this study terminate before reaching the number.

For the control multi-species model, cross-species model, and cross-species LOSO model we set the learning rate range as 0.0005, 0.0001, the dropout range as 0 and 0.2, the MLP layer number range as 1 and 2, and hidden neuron number range as 200 and 400. The maximum epoch number was set to 30.000, although in some CV processes, the training was not stopped by the early stopping mechanism before reaching the maximum epochs; But usually, this indicated that another hyperparameter set with a lower learning rate could do a better job, thus 30.000 was a suitable value to set to the maximum epoch number.

Moreover, we refined the Aytan-Aktug’s method for comparing the single-species-antibiotic model and multi-species models. In their original method, for each antibiotic, the metric of the multi-species model is computed based on samples from multiple species, so the metric measures the collective performance of genomes from multiple species. Therefore, when such a metric of the multi-species model was compared to the metric calculated for genomes from a specific species in the single-species model (**Supplemental Fig. S26**), the comparison was biased by the mixed-species samples. In contrast, for each antibiotic, we computed the multi-species model metric scores based on genomes solely belonging to one species–antibiotic combination for comparison (**Supplemental Fig. S24**).

##### Adaptations and hyperparameter selection of ML-based software Seq2Geno2Pheno

Seq2Geno2Pheno maps sequences to genotype (Seq2Geno) and then maps genotype to phenotype (Geno2Pheno). For benchmarking, we used the genome-precomputed version of Seq2Geno, which skips the assembly procedure and only uses the gene presence/absence (GPA) part in the main version of Seq2Geno. We used the default parameters set by Seq2Geno. k-mers (-k 6) were generated using KMC (-k6 -m24 -fm -ci0 -cs1677215) and kmc_dump (-ci0 -cs1677215). .

Regarding classifiers, four default classifiers of the Geno2Pheno (linear and non-linear support-vector machine, logistic regression, random forest) were evaluated. For each classifier, the default hyperparameter range was used for hyperparameter selection, the class_weight was set to “balance”, and random_state was set to 0. “GridSearchCV” module in the scikit-learn package was used for optimization, with “scoring” set to “f1_macro”. In single-species model nested CV evaluation (**Fig. 1A**), we selected the classifier with the highest F1-macro mean (resorting to the lowest F1-macro standard deviation in case of a tie) in the inner loop CV for each of the 10 outer loop evaluations.

##### Adaptations of and hyperparameter selection of ML-based software PhenotypeSeeker

We provided a modified version of PhenotypeSeeker for evaluation using nested CV, for two reasons. First, the native evaluation renders the information leakage of the test set into training sets via the k-mers selection procedure, which makes use of the whole dataset. Besides, the native version with the Python package “multiprocess.Manager” stopped at 99% completeness indicated by the progress bar for the "Conducting the k-mers specific chi-square tests'' step on our machine (nevertheless it might be the approximation issues), and led to suspected memory leaks in some cases. Therefore, we refactored PhenotypeSeeker’s workflow to single-threaded to avoid the use of a multiprocess package, and in the k-mers selection step, we only performed the Chi-squared test on the training set samples. We used the default parameters used by the native study in the feature extraction procedure, which included compilation of k-mer lists (-k 13), weighting, Chi-squared test, and filtering (a maximum of 1000 lowest *p*-valued k-mers were used by setting -n_kmer as 1000).

Regarding ML classifiers, the default classifier in the native article is logistic regression; but as a few more were provided in their codes, we evaluated their software on three classifiers (support vector machine, logistic regression with L1 penalty, random forest). For each classifier, the default hyperparameter range provided by the PhenotypeSeeker 1.0.0 command line version was used for hyperparameter optimization, the class_weight was set to "balance", and random_state was set to 0. The “GridSearchCV” module in the scikit-learn package was used for optimization, with “scoring” set to “f1_macro”. In single-species model nested CV evaluation (**Fig. 1A**), we used the same method as mentioned above in Seq2Geno2Pheno, to select the classifier for nested CV outer loop evaluation.

##### Hyperparameter selection of ML-based software Kover

We used the command line and bound selection version of Kover, running it 10 times, each time using a different fold as the test set and the rest as the training set. We used the default k-mers length of 31. Regarding classifiers, classification and regression trees and set covering machines were evaluated. The default hyperparameter range was used for hyperparameter optimization. For set covering machines, max-rules was 10, and the trade-off parameter P was (0.1 0.178 0.316 0.562 1.0 1.778 3.162 5.623 10.0 999999.0). For classification and regression trees, max-depth was 20, min-samples-split was 2, and class-importance was (0.25 0.5 0.75 1.0). In single-species model nested CV evaluation (**Fig. 1A**), at each iteration, an extra CV, maximizing “F1-macro”, was performed on the training set (9 folds) to select a classifier (classification and regression trees and set covering machines) for reporting the final performance of Kover. The ML baseline in the Kover article, which returns the majority’s phenotype in the training set for samples in the corresponding test set, was used as the ML baseline for our benchmarking.

#### Misclassification analysis

The misclassification frequency ratio for a specific species was determined for each genome within that species. This ratio quantified the number of times a genome was correctly predicted with random folds but misclassified with phylogeny-aware or homology-aware folds. To normalize this ratio, it was divided by two factors: the number of species–antibiotic combinations in which the genome was involved and the total number of ML-based AMR prediction methods applied, which was four. Subsequently, we annotated the corresponding phylogenetic tree, which was generated in the phylogeny-aware sample partitioning procedure using Seq2Geno2Pheno, with the misclassification frequencies of genomes. This annotation process was conducted using iTOL V6.6 [[21]](https://paperpile.com/c/6VFvZ6/el2Iz).

#### Training ML models on the whole dataset

For PhenotypeSeeker, we trained a model for each species–antibiotic combination on the corresponding whole dataset using the optimal set of hyperparameters. Specifically, we grouped all sets of hyperparameters that were selected as the optimal in the inner loop of nested CV; then we evaluated each hyperparameter-set on the 10 folds 10 times, each time using one fold for testing and the other 9 folds for training, resulting in the F1-macro mean and standard deviation (both rounded to 2 decimal points) of the 10 iterations; finally for training on the whole dataset, we selected the set of hyperparameters with the highest F1-macro mean.

Regarding Kover, for each species–antibiotic combination, we selected the best from the classification and regression trees and the set covering machines based on iterative bound selection evaluation, and then directly trained the model on the corresponding whole dataset.

In both ML methods’ selection procedures, if multiple hyperparameter sets had the tied highest F1-macro means, we selected the one with the lowest standard deviation; and if multiple hyperparameter sets had tied highest F1-macro means and tied lowest standard deviation, we selected the set covering machines for Kover, and selected the one with the highest F1-macro mean before decimal rounding for PhenotypeSeeker; and if multiple hyperparameter sets had tied highest F1-macro means and tied lowest standard deviation before rounding, we randomly selected one for PhenotypeSeeker. Selections were applied with random, phylogeny-aware, and homology-aware folds, respectively, resulting in three trained models for each species–antibiotic combination.
