## Supplemental Fig for "Assessing computational predictions of antimicrobial resistance phenotypes from microbial genomes"

### Supplemental Figure S1-S26

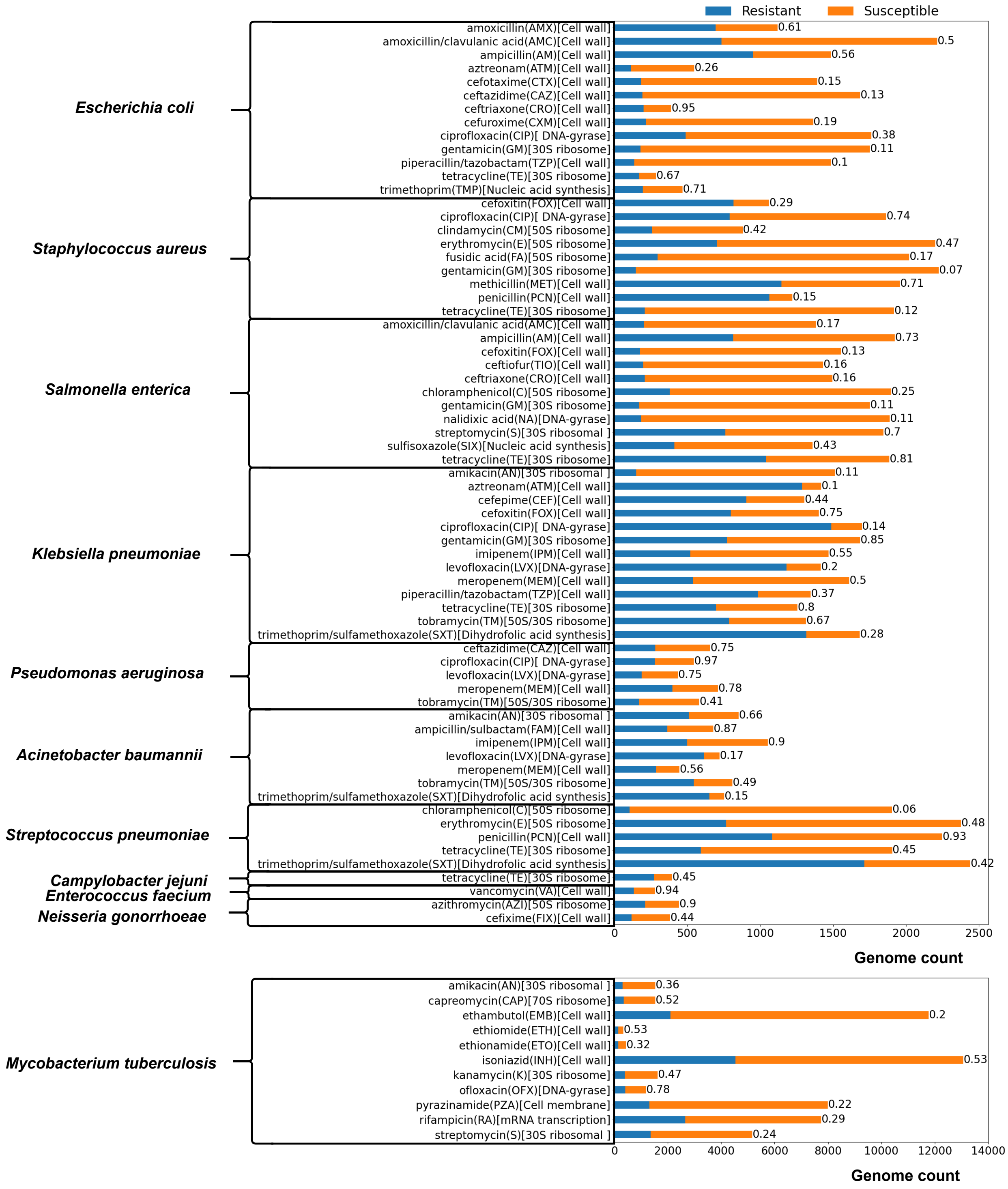


**Figure S1.** Seventy-eight datasets for the single-species evaluation, spanning 11 species, annotated with resistant (orange) and susceptible (blue) genome isolates, the ratio of minority class over majority class, and the antibiotic action of mechanism site.

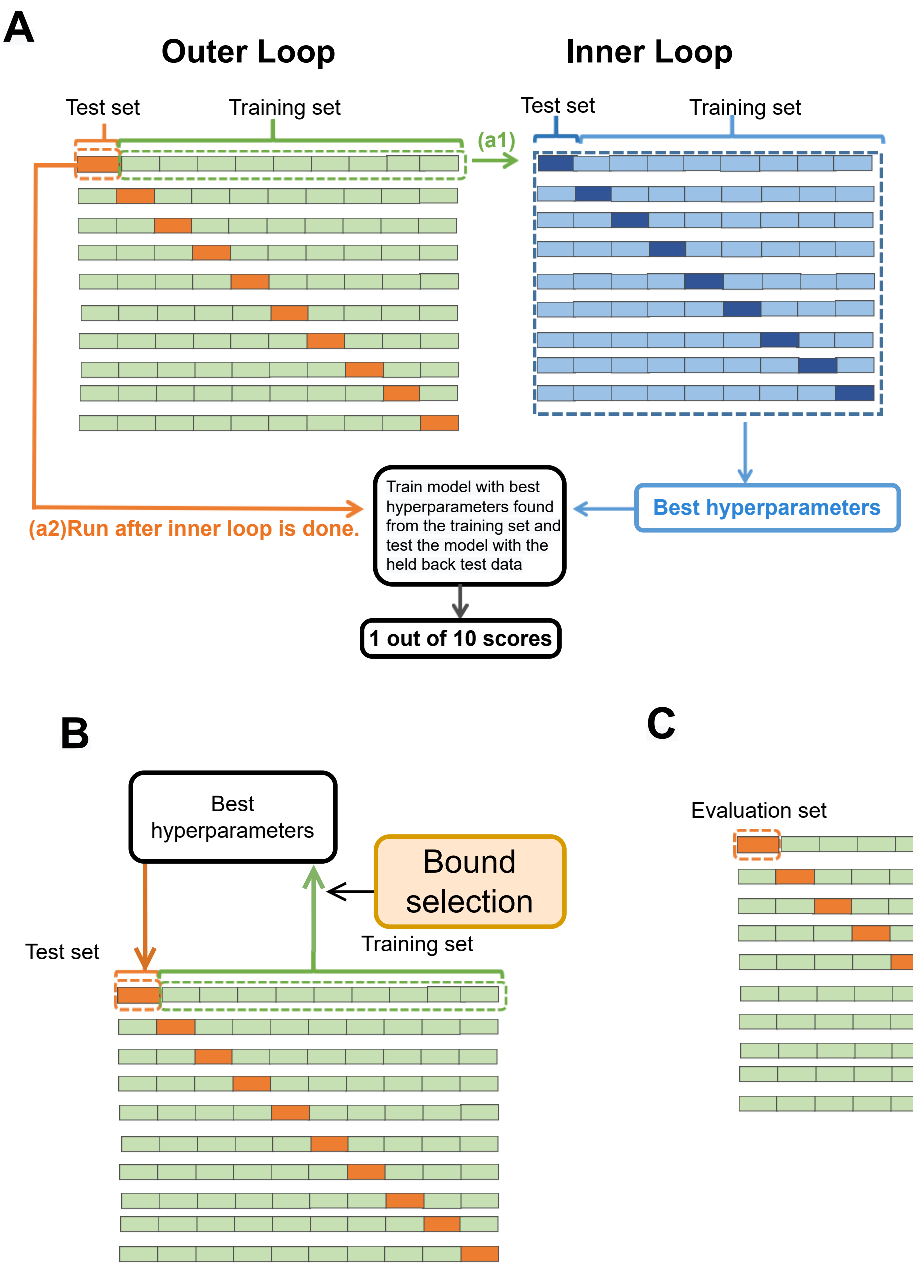


**Figure S2.** Three evaluation approaches in the benchmarking workflow. A. Nested cross-validation (graph adapted from `https://mlfromscratch.com/nested-cross-validation-python-code`). (a1) In each outer loop, a test set (a fold) was held back, and the training set (9 folds) was further split into training and test sets (a fold and 8 folds respectively), which were then fed into the inner loop, in which the best hyperparameter set was chosen via a 9-fold CV. (a2) We trained the model on the training set in the outer loop using the chosen hyperparameter set, which was then tested on the holdout test set. We iterated (a1) and (a2) 10 times, each time holding a different fold as the test set. B. Iterative bound selection[1] evaluation for Kover. At each iteration, the model was trained on the entire training set (9 folds); the best classifier was chosen from the two classifiers produced by Kover via a 9-fold conventional CV, which was then used to evaluate the test set (a fold). This procedure was iterated over each of the 10 folds. C. Iterative evaluation for ResFinder 4.0. Each fold was evaluated sequentially.

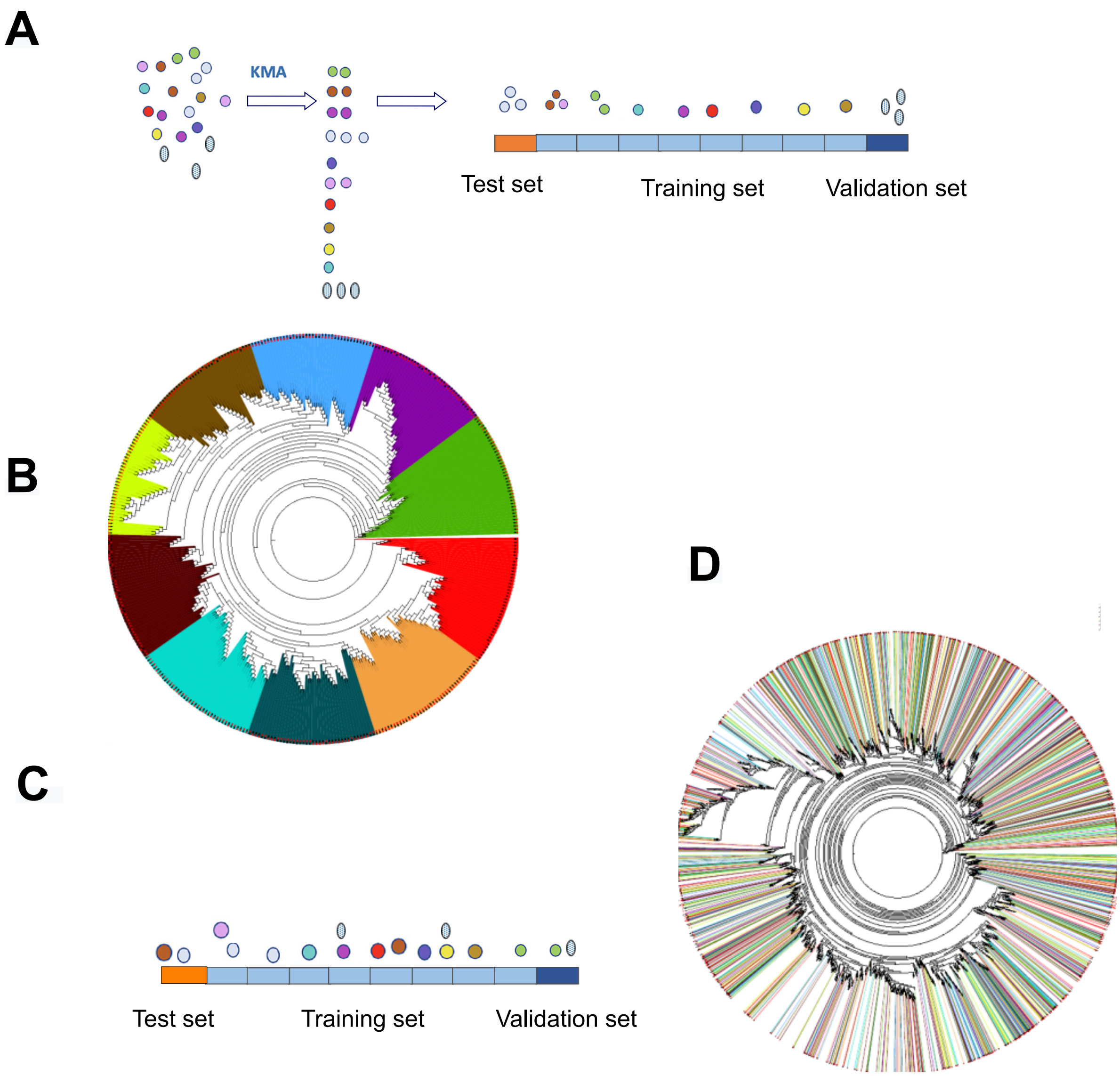


**Figure S3.** Three sample partitioning methods. (A) Homology-aware partitioning[2]. Similar samples (represented by the same color) in each cluster stay together either in the training set, the validation set, or the test set in evaluation. The Visualization was adapted from Aytan-Aktug et al. [2]. (B) Phylogeny-aware partitioning[3]. To achieve nearly equal-sized folds (each represented by 10 colors), some phylogenetically close strains might be distributed into separate folds. (C, D) The random partitioning method generates 10 nearly equal-sized folds, each of which is composed of dissimilar strains and evolutionarily divergent strains.

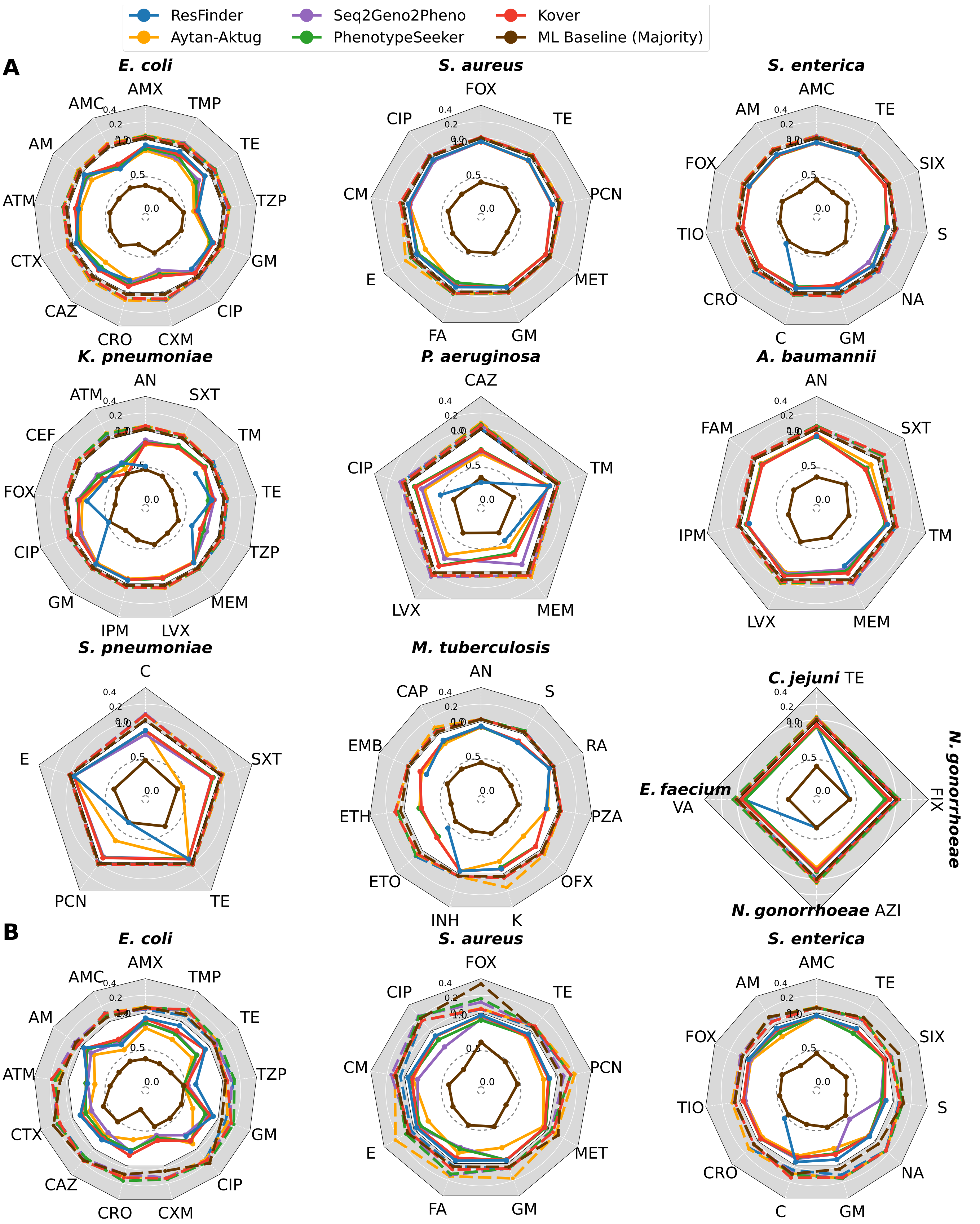


**Figure S4.** Continued on next page

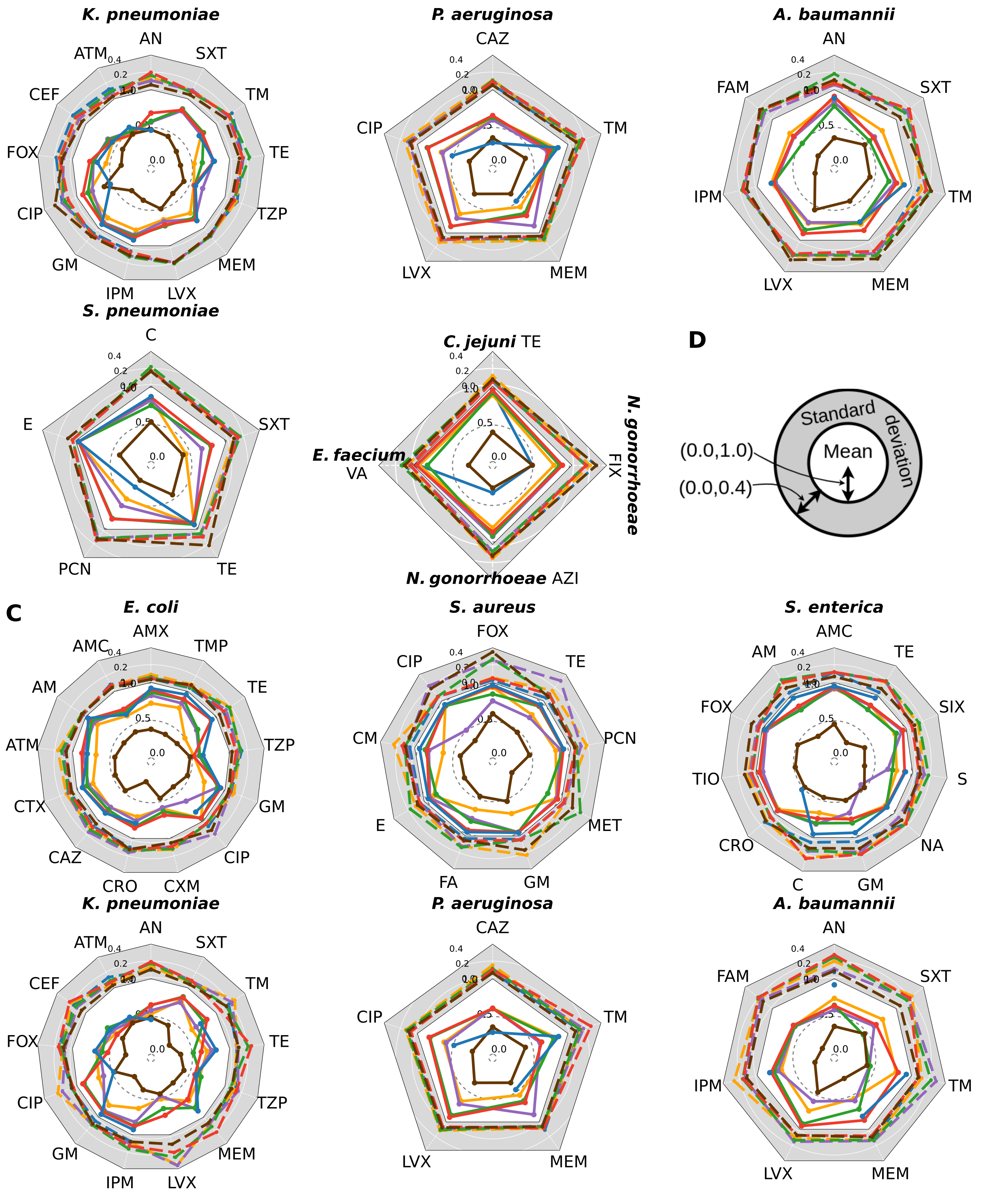


**Figure S4.** Continued on next page

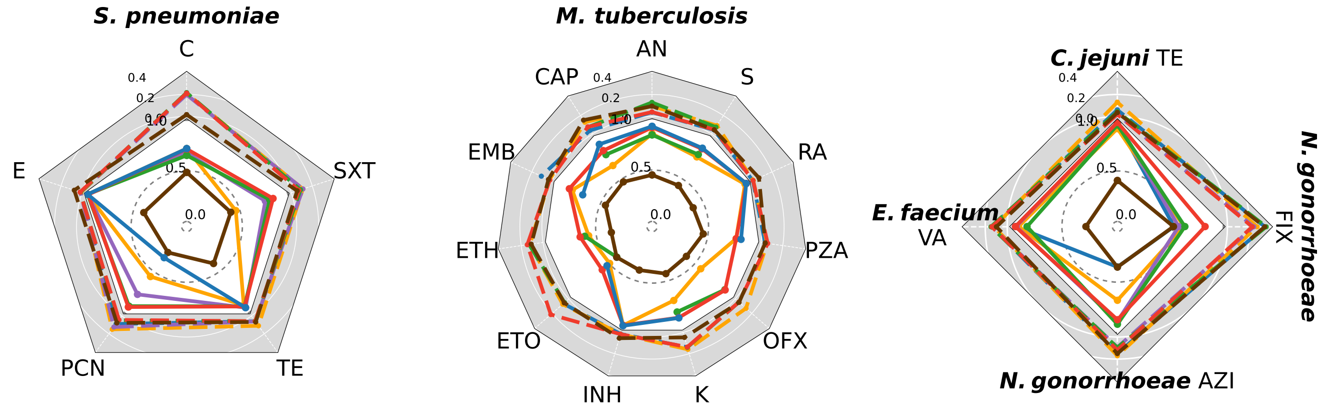


**Figure S4.** Performance of five AMR prediction methods and the ML baseline method (Majority) with (A) random folds, (B) phylogeny-aware folds, and (C) homology-aware folds. (D). The F1-macro mean is represented by the inner circle of a solid line with a white background, and the standard deviation is represented by the outer gray ring of the dashed line. The closer to the outside of the solid line, the better the software; the closer to the center of the dashed line, the more stable the software. The mapping of the acronyms to antibiotics is in Supplemental Table S2.

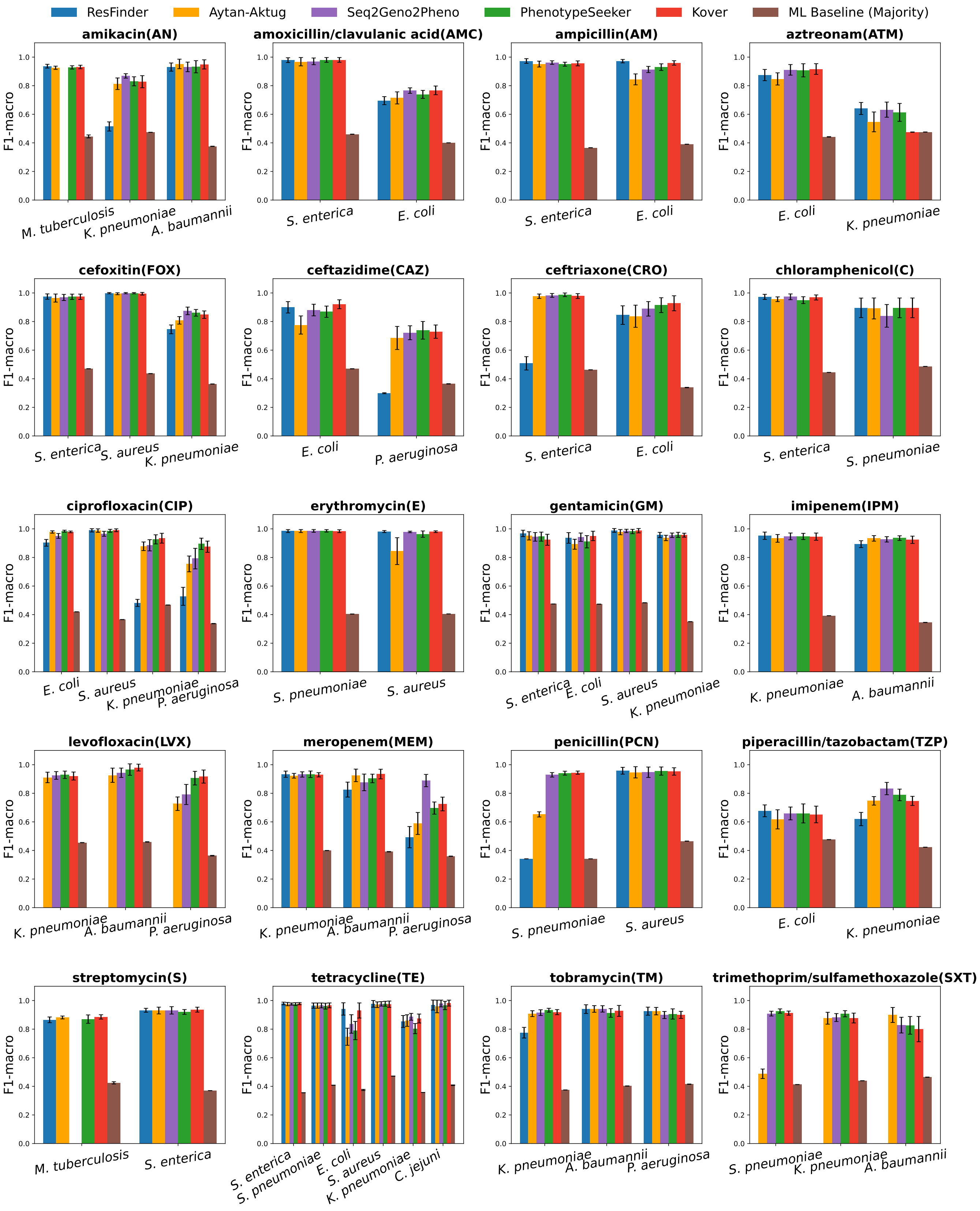


**Figure S5.** Performance (F1-macro) of the methods with antibiotics shared by multiple species under the scenarios of random folds.

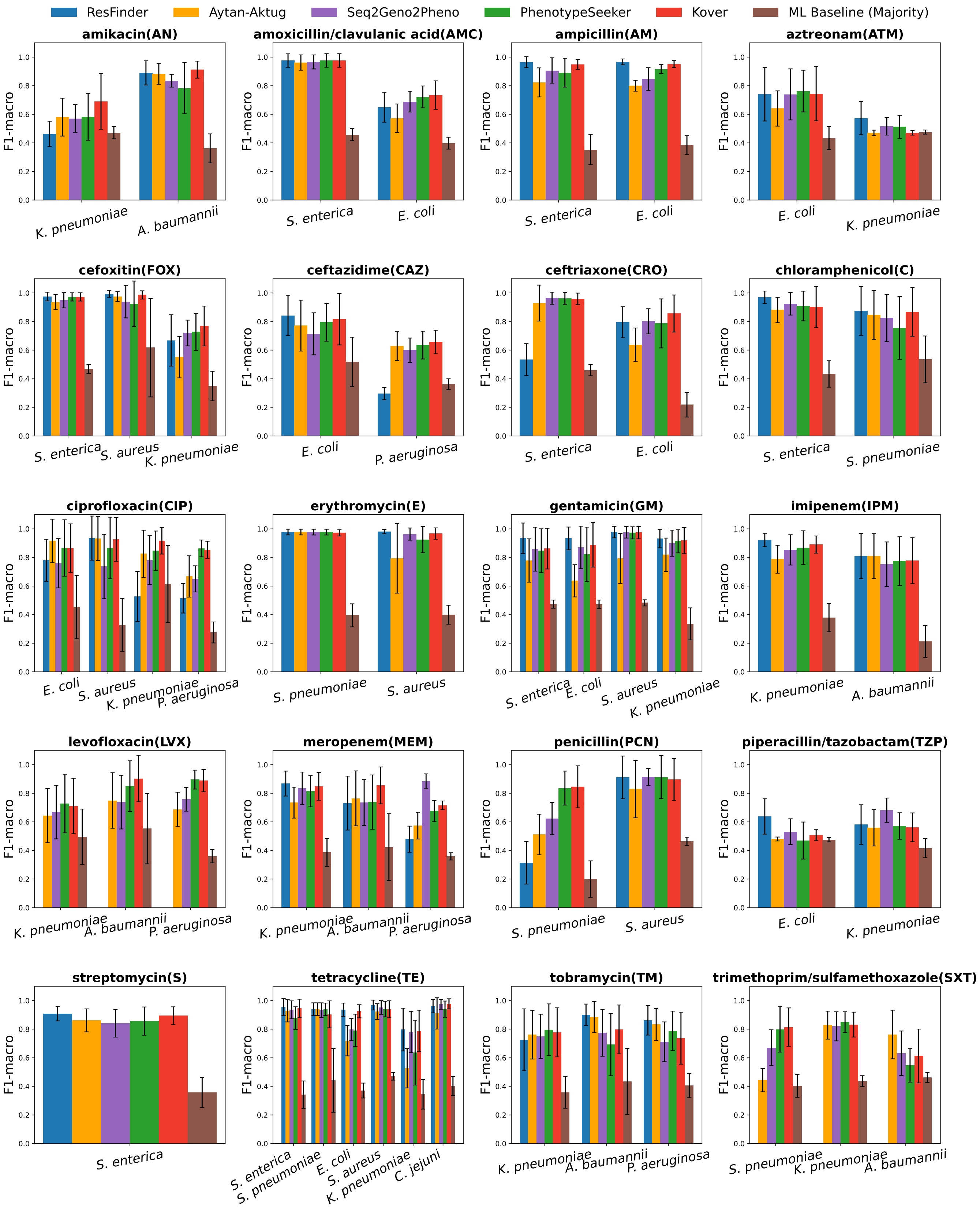


**Figure S6.** Performance (F1-macro) of the methods with antibiotics shared by multiple species under the scenarios of phylogeny-aware folds.

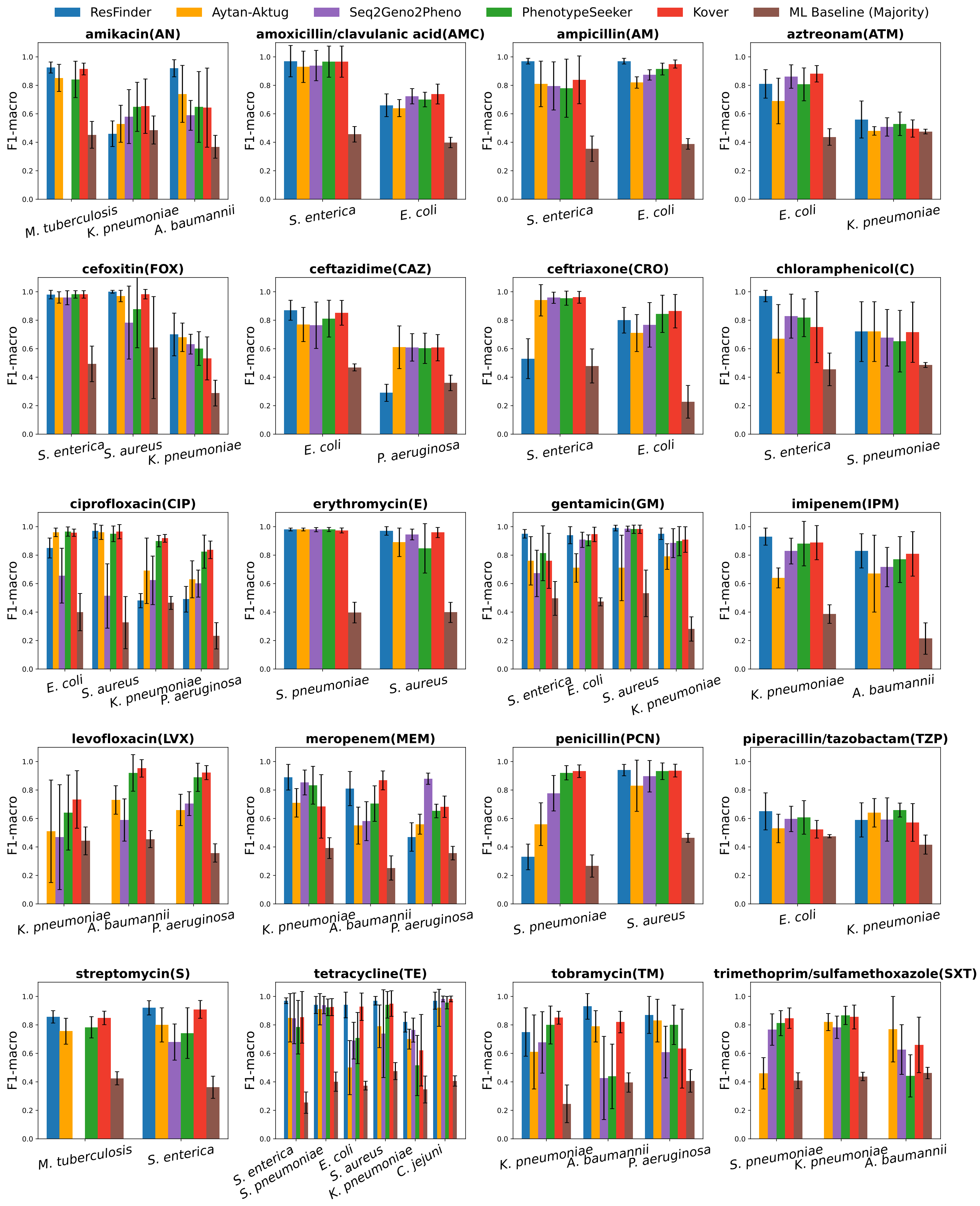


**Figure S7.** Performance (F1-macro) of the methods with antibiotics shared by multiple species under the scenarios of homology-aware folds.

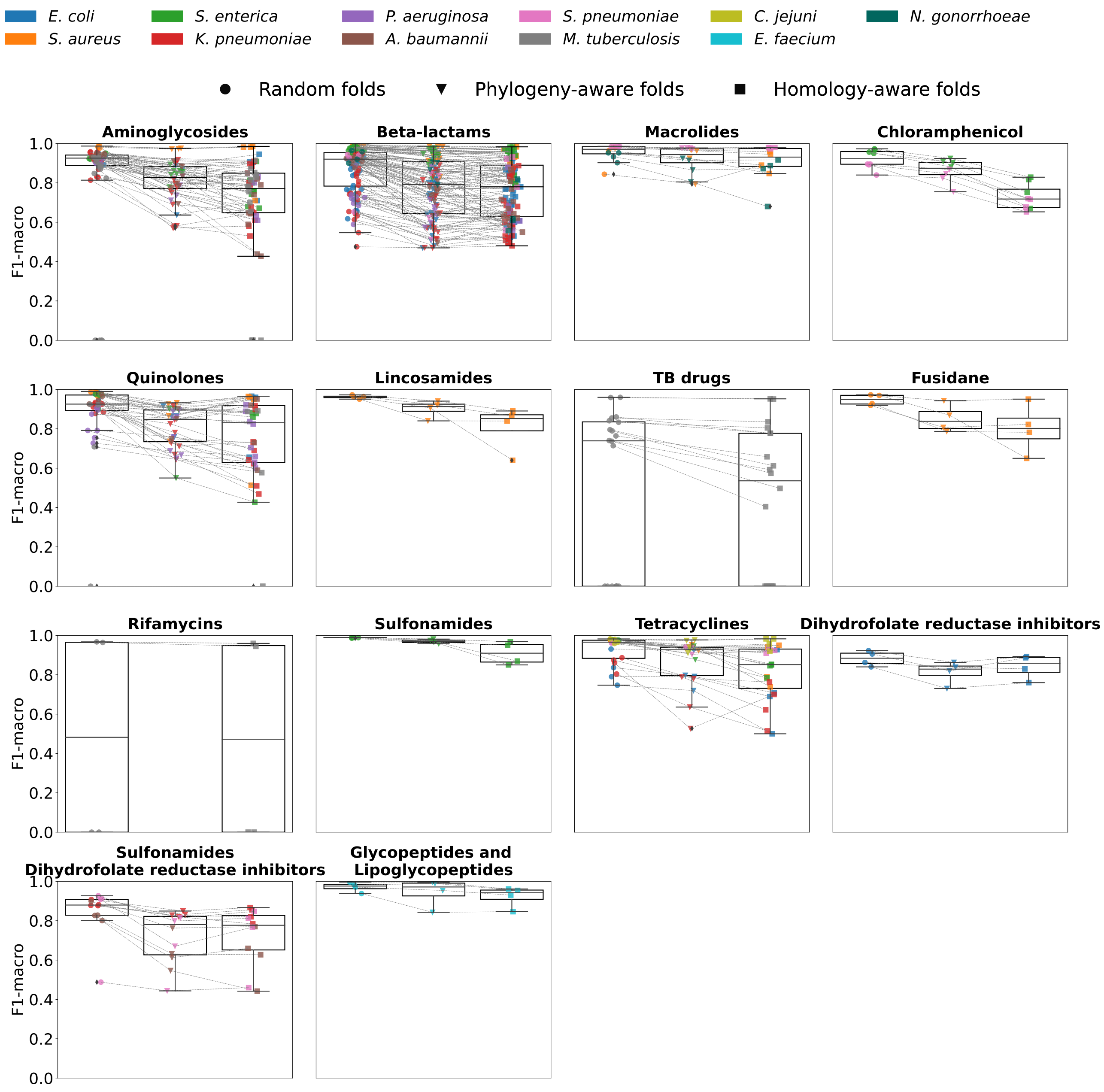


**Figure S8.** Changes in performance (F1-macro mean) for each antibiotic class with random folds, phylogeny-aware folds, and homology-aware folds. The color of the dots represents species.

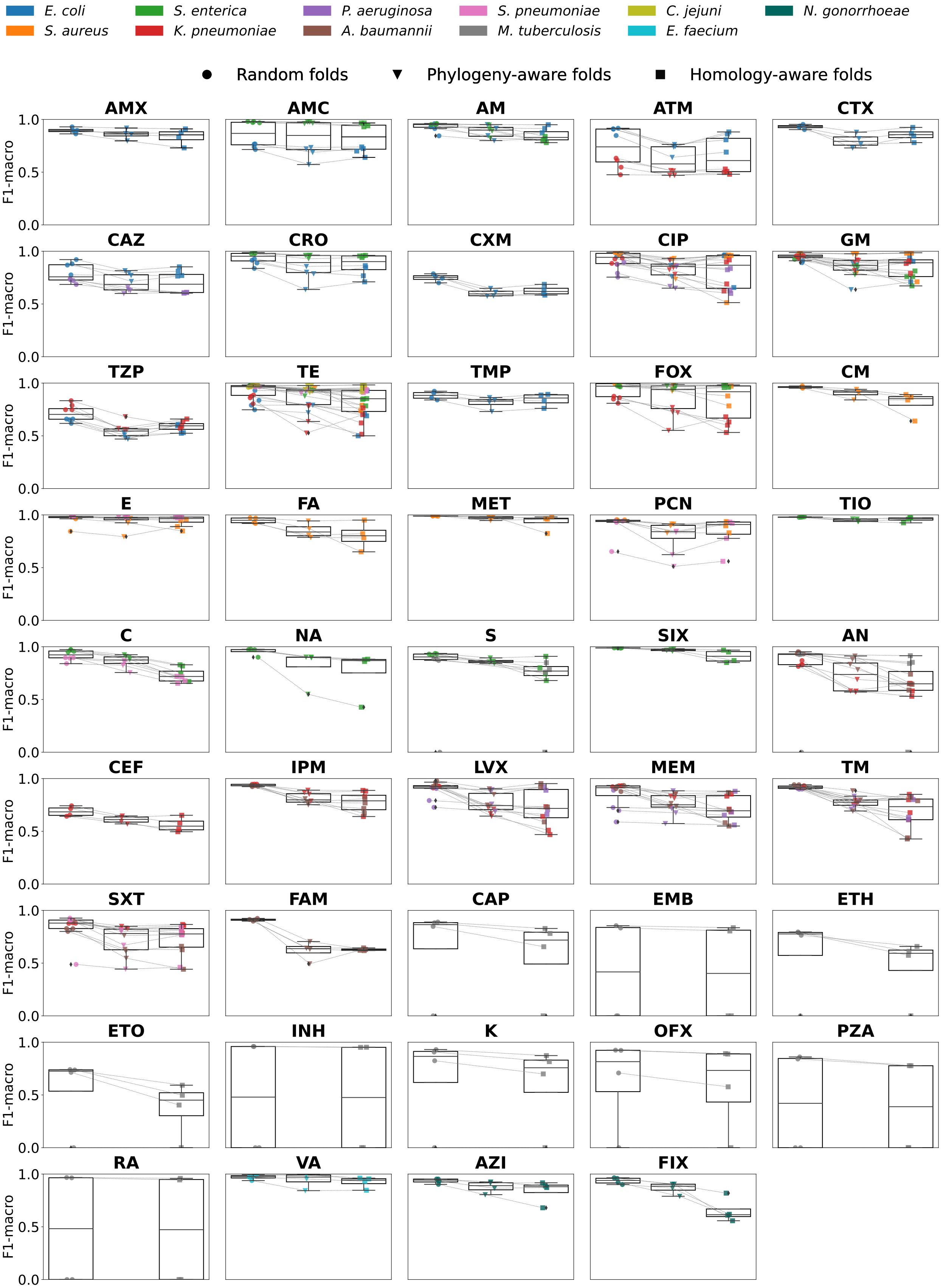


**Figure S9.** Changes in performance (F1-macro mean) for each antibiotic with random folds, phylogeny-aware folds, and homology-aware folds. The color of the dots represents species. The mapping of the acronyms to antibiotics is in Supplemental Table S2.

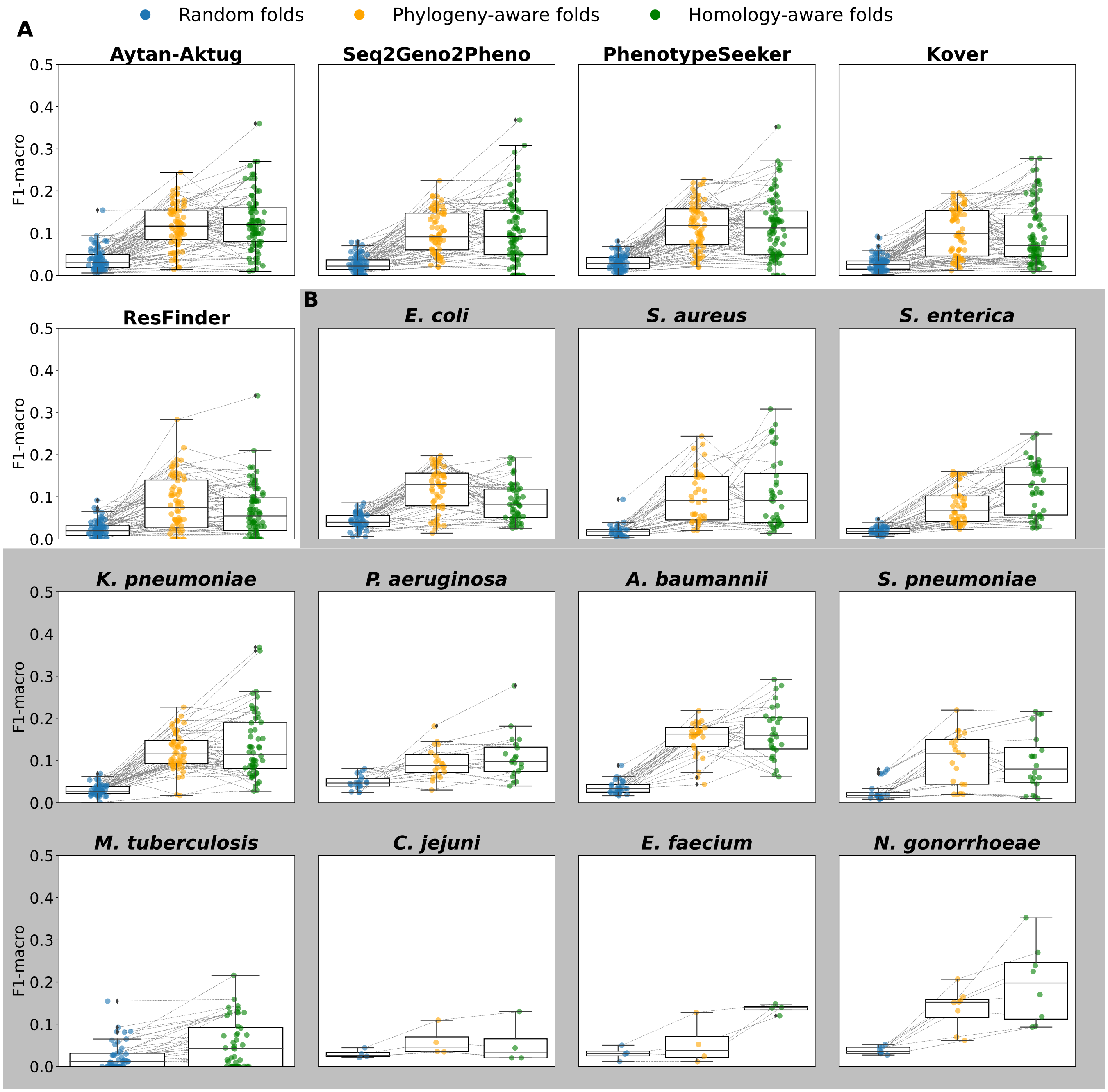


**Figure S10.** Changes in performance stability (standard deviation of F1-macro) with random folds, phylogeny-aware folds, and homology-aware folds. A. Changes in performance stability for each method. Each dot represents one species-antibiotic combination for the corresponding method. B. Changes in performance stability for each species for the ML-based methods Kover, PhenotypeSeeker, Seq2Geno2Pheno, and Aytan-Aktug. Each point represents one ML-based method–antibiotic combination for the corresponding species.

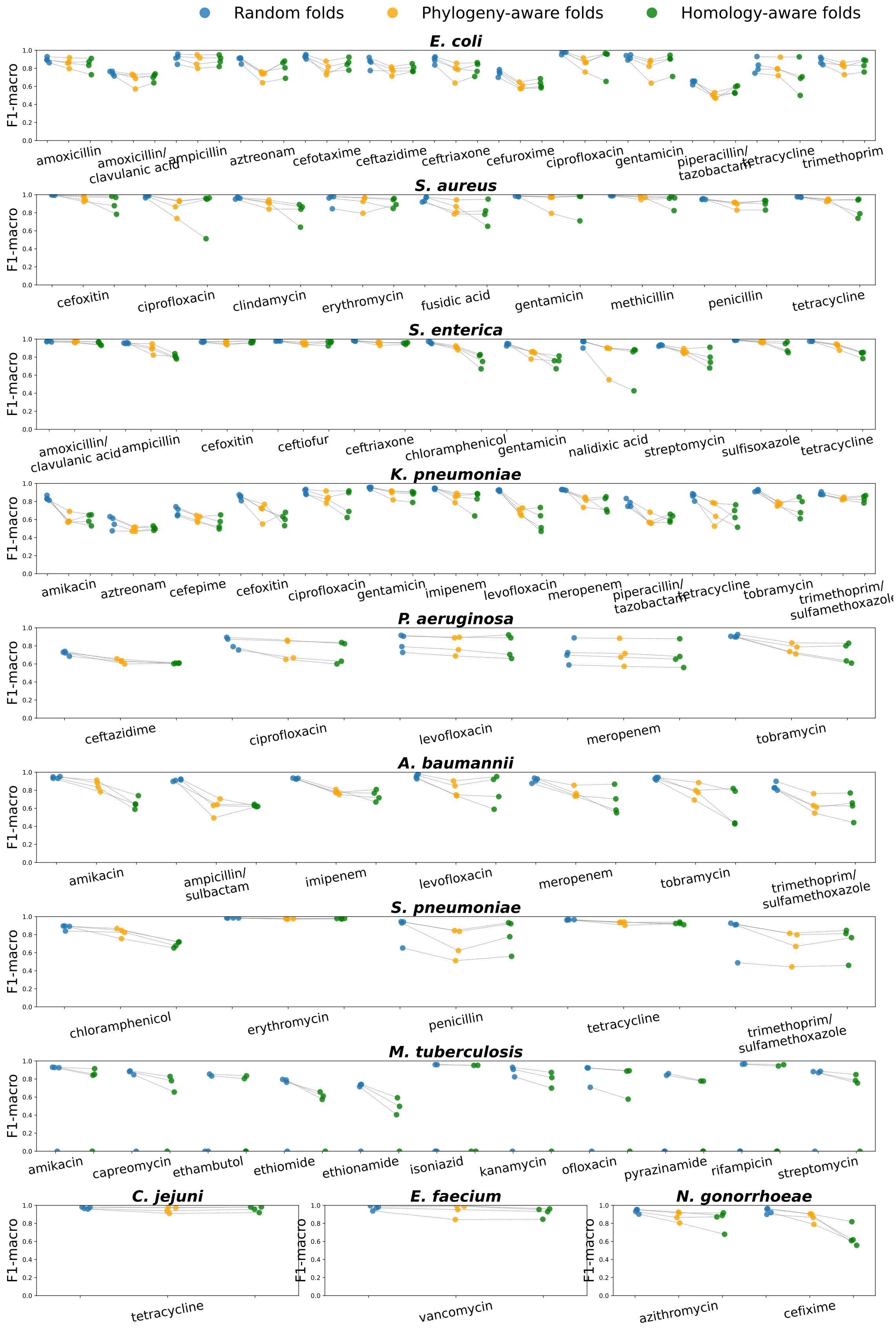
Random folds
Phylogeny-aware folds
Homology-aware folds
E. coli
S. aureus
S. enterica
K. pneumoniae
P. aeruginosa
A. baumannii
S. pneumoniae
M. tuberculosis
C. jejuni
E. faecium
N. gonorrhoeae
F1-macro
amoxicillin
amoxicillin/clavulanic acid
ampicillin
aztreonam
cefotaxime
ceftazidime
ceftriaxone
cefuroxime
ciprofloxacin
gentamicin
piperacillin/tazobactam
tetracycline
trimethoprim
cefoxitin
clindamycin
erythromycin
fusidic acid
methicillin
penicillin
ceftiofur
chloramphenicol
nalidixic acid
streptomycin
sulfisoxazole
amikacin
cefepime
imipenem
levofloxacin
meropenem
tobramycin
trimethoprim/sulfamethoxazole
ampicillin/sulbactam
capreomycin
ethambutol
ethiomide
ethionamide
isoniazid
kanamycin
ofloxacin
pyrazinamide
rifampicin
vancomycin
azithromycin
cefixime

**Figure S11.** Changes in performance (F1-macro mean) for each species-antibiotic combination with random folds, phylogeny-aware folds, and homology-aware folds. Each dot represents one ML-based method for the corresponding species and antibiotic.

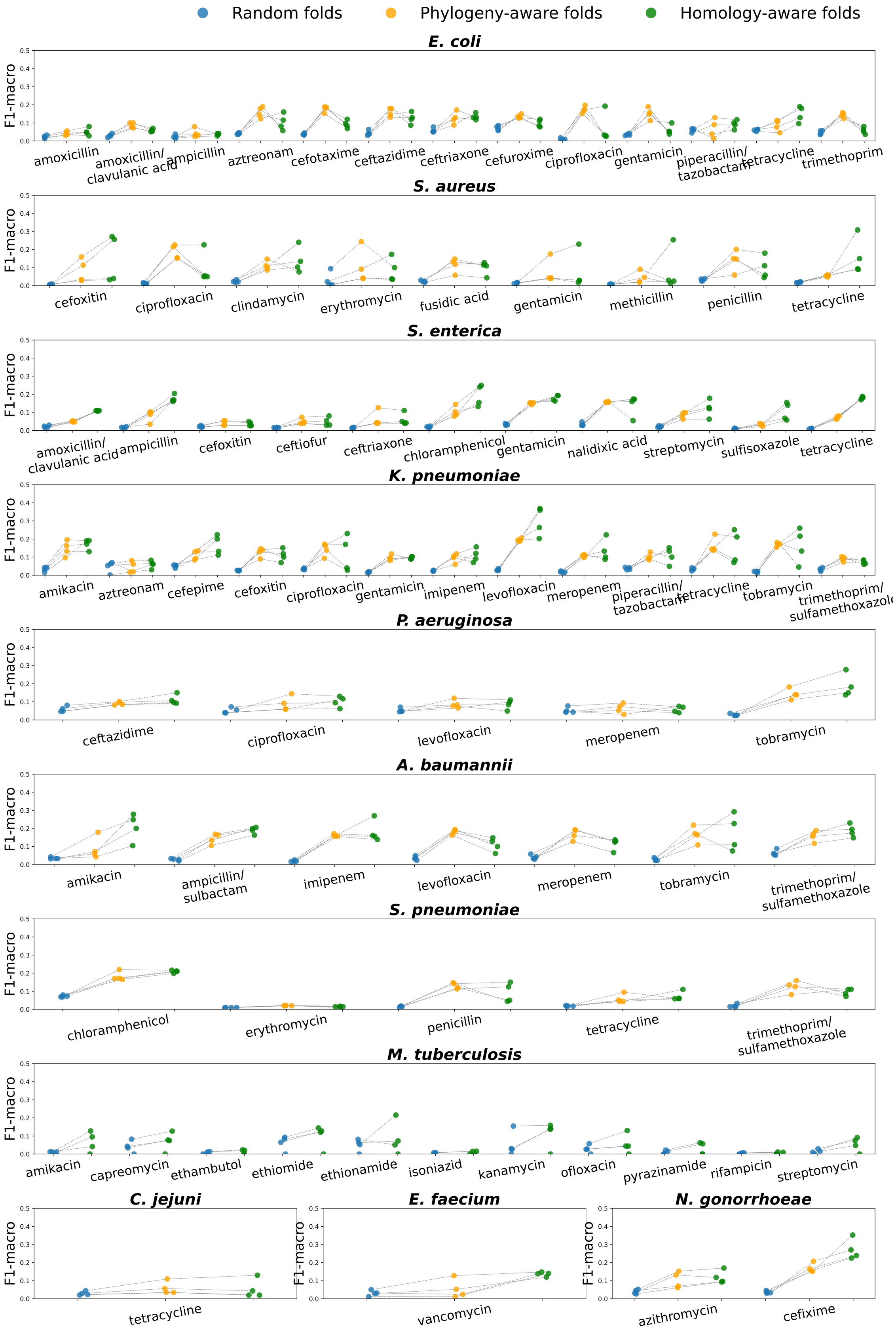
Random folds
Phylogeny-aware folds
Homology-aware folds
E. coli
F1-macro
amoxicillin
amoxicillin/ clavulanic acid
ampicillin
aztreonam
cefotaxime
ceftazidime
ceftriaxone
cefuroxime
ciprofloxacin
gentamicin
piperacillin/ tazobactam
tetracycline
trimethoprim
S. aureus
cefoxitin
ciprofloxacin
clindamycin
erythromycin
fusidic acid
gentamicin
methicillin
penicillin
tetracycline
S. enterica
amoxicillin/ clavulanic acid
ampicillin
cefoxitin
ceftiofur
ceftriaxone
chloramphenicol
gentamicin
nalidixic acid
streptomycin
sulfisoxazole
tetracycline
K. pneumoniae
amikacin
aztreonam
cefepime
cefoxitin
ciprofloxacin
gentamicin
imipenem
levofloxacin
meropenem
piperacillin/ tazobactam
tetracycline
tobramycin
trimethoprim/ sulfamethoxazole
P. aeruginosa
ceftazidime
ciprofloxacin
levofloxacin
meropenem
tobramycin
A. baumannii
amikacin
ampicillin/ sulbactam
imipenem
levofloxacin
meropenem
tobramycin
trimethoprim/ sulfamethoxazole
S. pneumoniae
chloramphenicol
erythromycin
penicillin
tetracycline
trimethoprim/ sulfamethoxazole
M. tuberculosis
amikacin
capreomycin
ethambutol
ethiomide
ethionamide
isoniazid
kanamycin
ofloxacin
pyrazinamide
rifampicin
streptomycin
C. jejuni
tetracycline
E. faecium
vancomycin
N. gonorrhoeae
azithromycin
cefixime

**Figure S12.** Changes in performance stability (standard deviation of F1-macro) for each species-antibiotic combination with random folds, phylogeny-aware folds, and homology-aware folds. Each dot represents one ML-based method for the corresponding species and antibiotic.

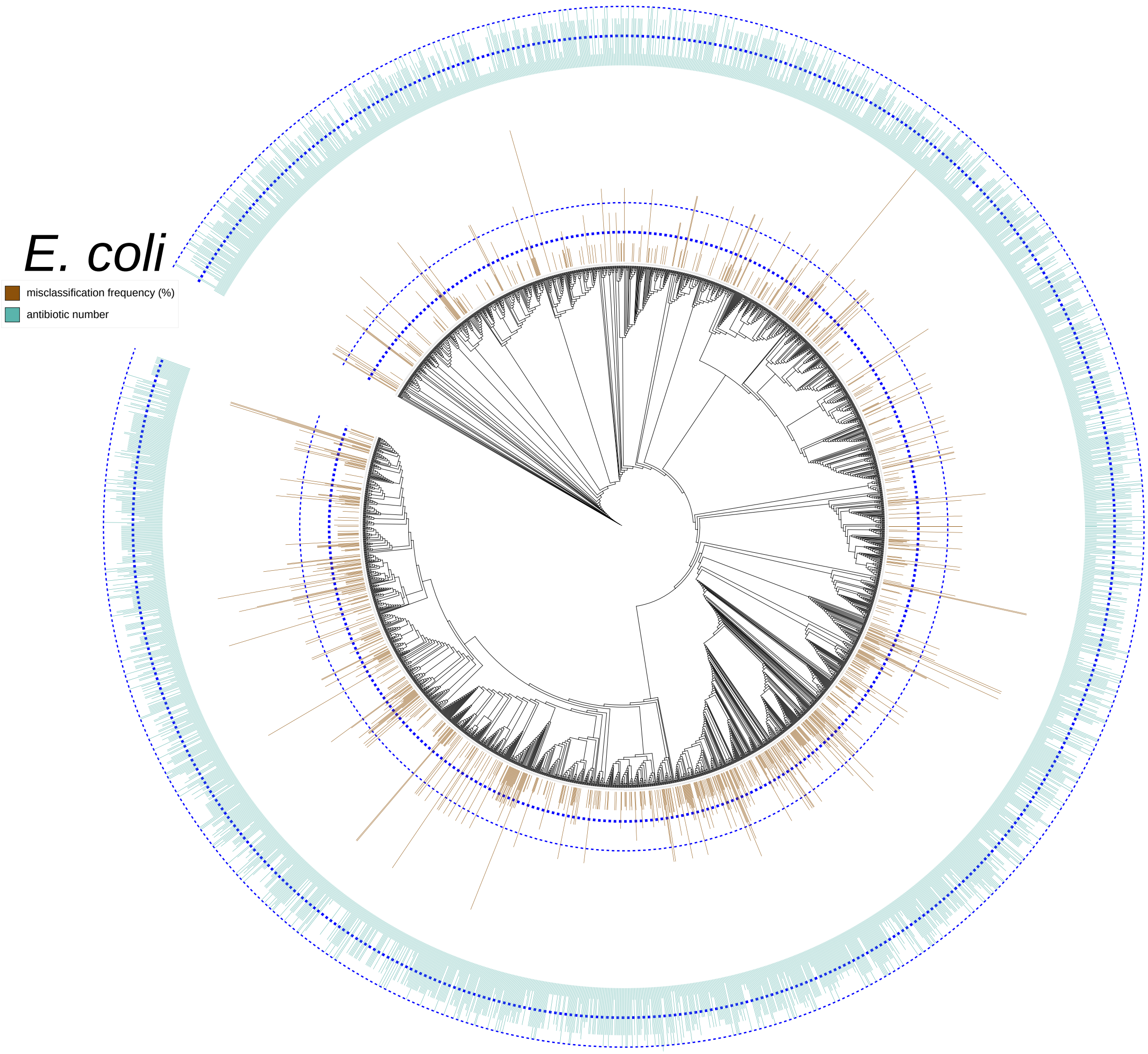


**Figure S13.** Distribution of misclassified genomes of *E. coli* on the phylogeny tree. The height of the brown bar for each leave, (i.e. a genome) stands for the misclassification frequency ratio that the genome was correctly predicted with random folds but misclassified with phylogeny-aware or homology-aware folds, normalized by the number of antibiotic combinations the genome was involved, which is 13, and by the number of ML-based AMR prediction methods, which is four. The height of the green bar for each leave (i.e. a genome) stands for the number of antibiotics for which the genome's phenotype was annotated. The inner dashed blue lines in both bar plots indicate an occurrence count of 5, while the outer dashed blue lines mark 10. This phylogeny tree was visualized and annotated using iTOL V6.6[4].

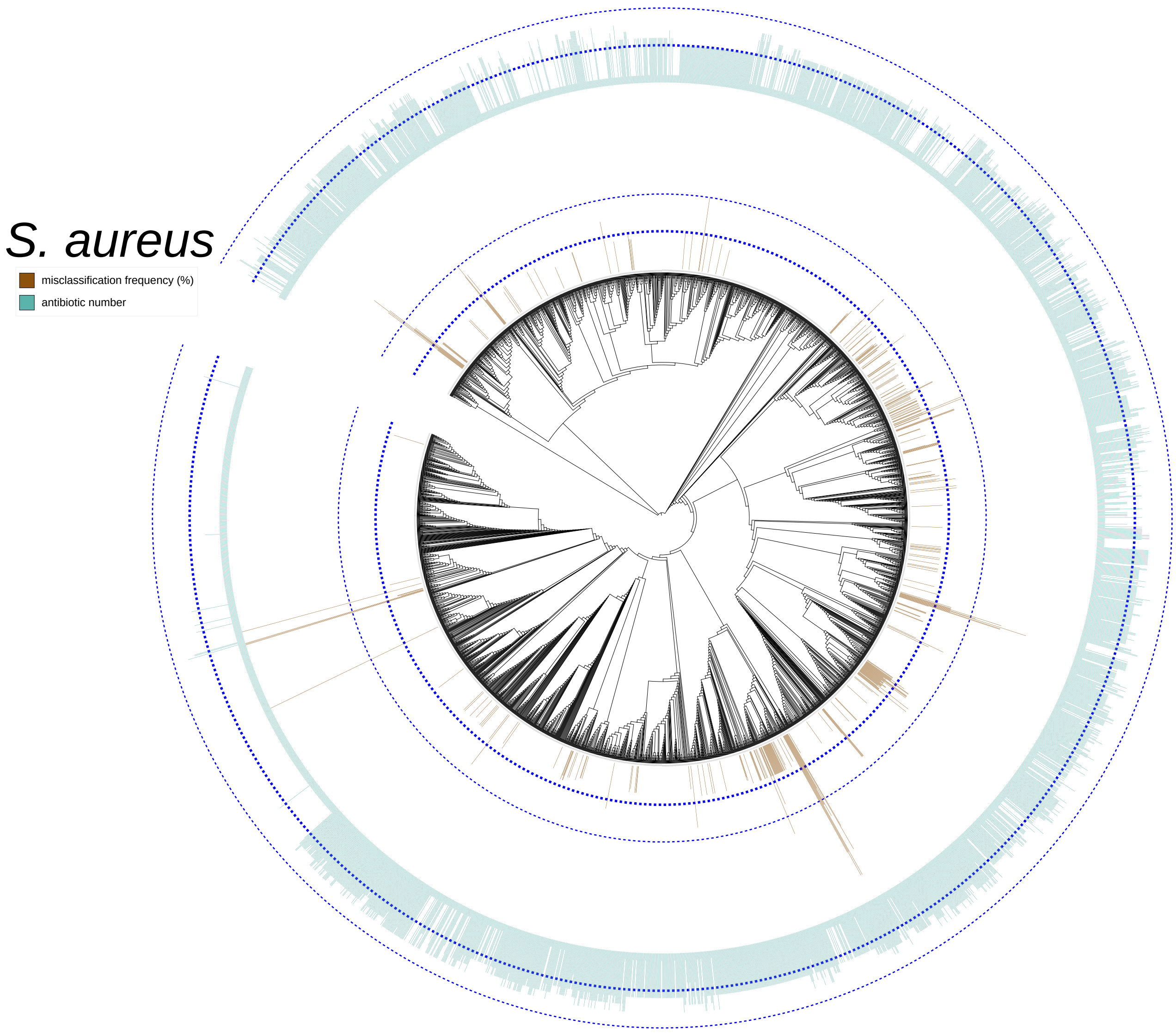


**Figure S14.** Distribution of misclassified genomes of *S. aureus* on the phylogeny tree. The height of the brown bar for each leave (i.e. a genome) stands for the misclassification frequency ratio that the genome was correctly predicted with random folds but misclassified with phylogeny-aware or homology-aware folds, normalized by the number of antibiotic combinations the genome was involved, which is 9, and by the number of ML-based AMR prediction methods, which is four. The height of the green bar for each leave (i.e. a genome) stands for the number of antibiotics for which the genome's phenotype was annotated. The inner dashed blue lines in both bar plots indicate an occurrence count of 5, while the outer dashed blue lines mark 10. This phylogeny tree was visualized and annotated using iTOL V6.6[4].

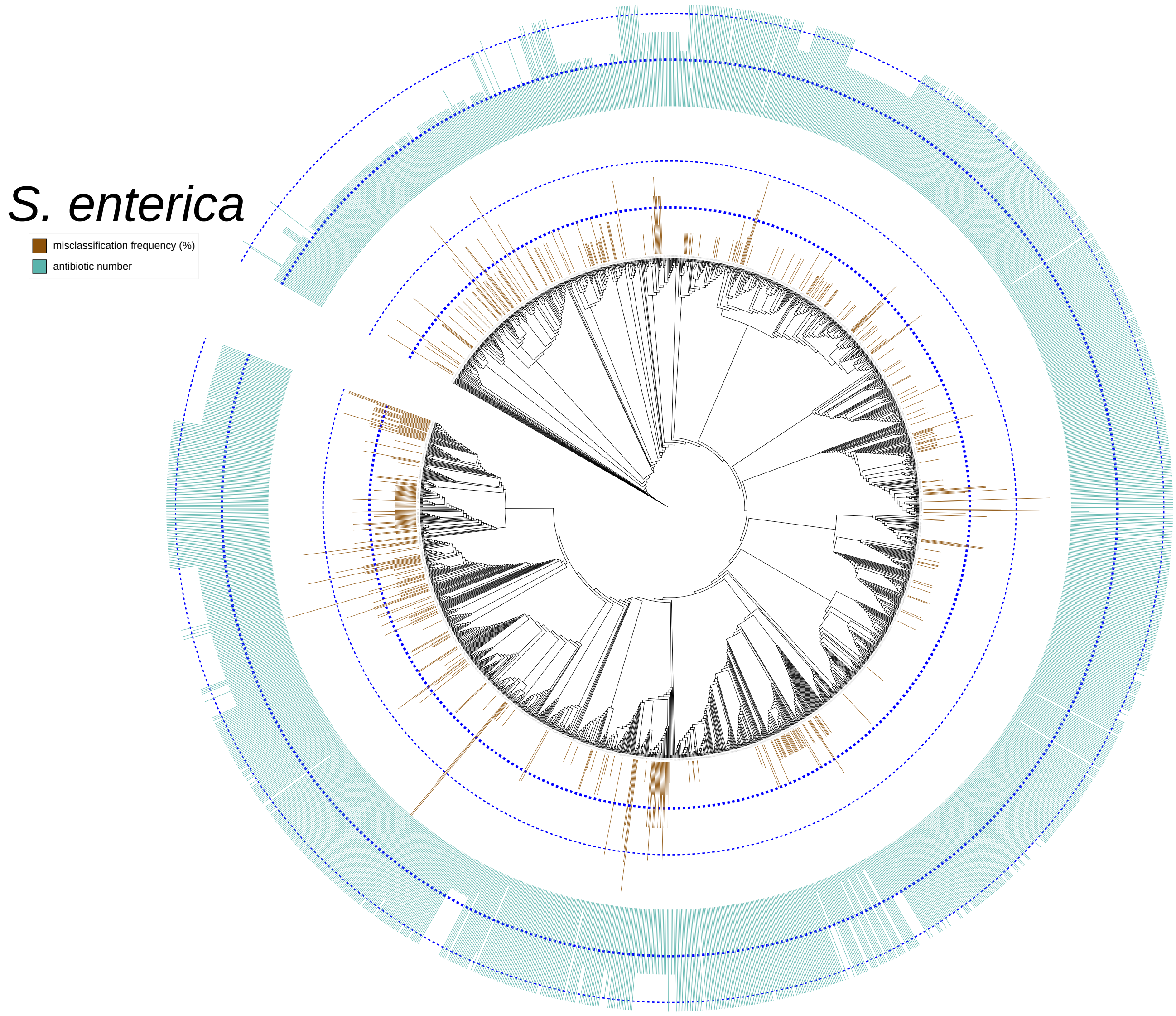


**Figure S15.** Distribution of misclassified genomes of *S. enterica* on the phylogeny tree. The height of the brown bar for each leave (i.e. a genome) stands for the misclassification frequency ratio that the genome was correctly predicted with random folds but misclassified with phylogeny-aware or homology-aware folds, normalized by the number of antibiotic combinations the genome was involved, which is 11, and by the number of ML-based AMR prediction methods, which is four. The height of the green bar for each leave (i.e. a genome) stands for the number of antibiotics for which the genome's phenotype was annotated. The inner dashed blue lines in both bar plots indicate an occurrence count of 5, while the outer dashed blue lines mark 10. This phylogeny tree was visualized and annotated using iTOL V6.6[4].

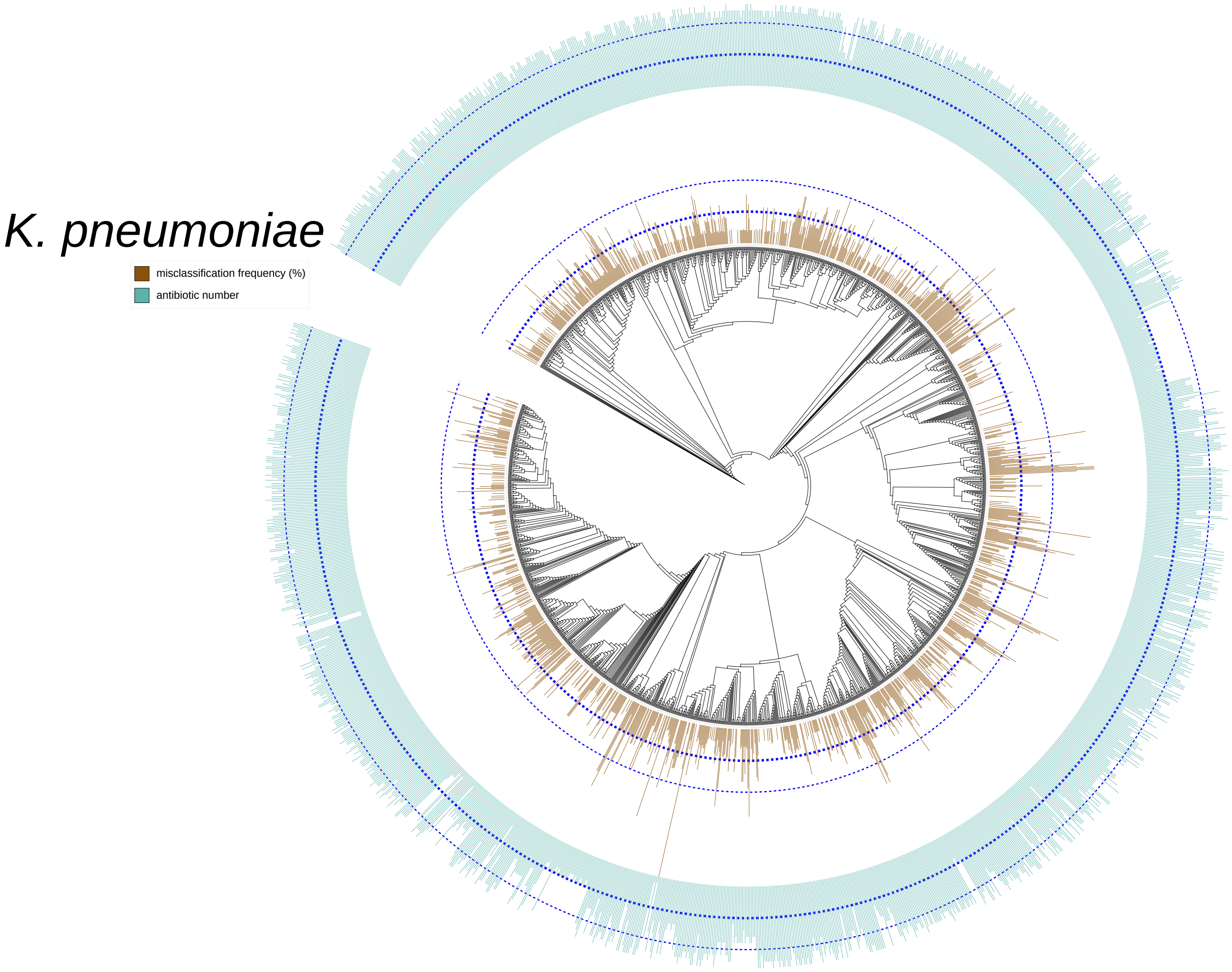


**Figure S16.** Distribution of misclassified genomes of *K. pneumoniae* on the phylogeny tree. The height of the brown bar for each leave (i.e. a genome) stands for the misclassification frequency ratio that the genome was correctly predicted with random folds but misclassified with phylogeny-aware or homology-aware folds, normalized by the number of antibiotic combinations the genome was involved, which is 13, and by the number of ML-based AMR prediction methods, which is four. The height of the green bar for each leave (i.e. a genome) stands for the number of antibiotics for which the genome's phenotype was annotated. The inner dashed blue lines in both bar plots indicate an occurrence count of 5, while the outer dashed blue lines mark 10. This phylogeny tree was visualized and annotated using iTOL V6.6[4].

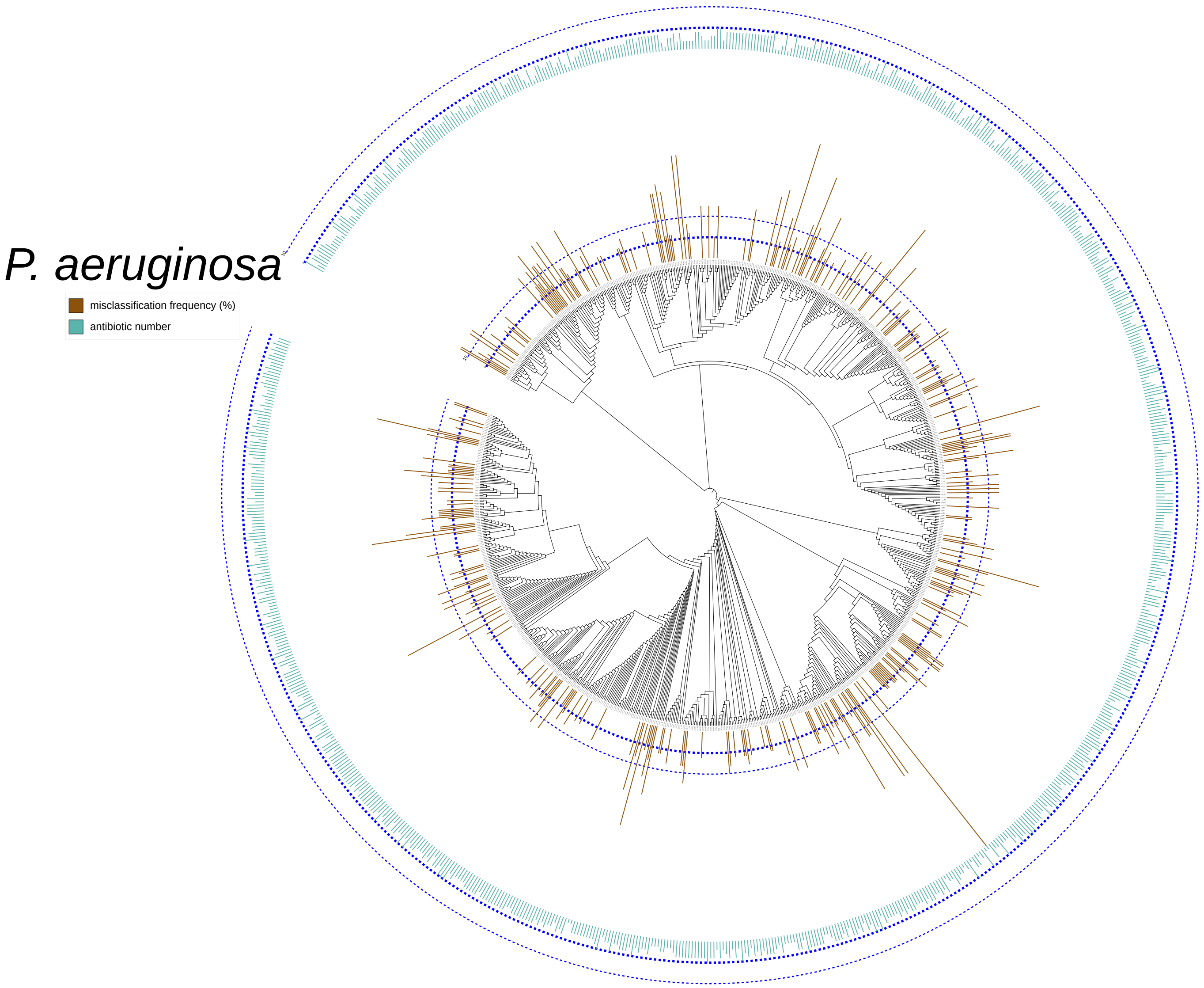


**Figure S17.** Distribution of misclassified genomes of *P. aeruginosa* on the phylogeny tree. The height of the brown bar for each leave (i.e. a genome) stands for the misclassification frequency ratio that the genome was correctly predicted with random folds but misclassified with phylogeny-aware or homology-aware folds, normalized by the number of antibiotic combinations the genome was involved, which is 5, and by the number of ML-based AMR prediction methods, which is four. The height of the green bar for each leave (i.e. a genome) stands for the number of antibiotics for which the genome's phenotype was annotated. The inner dashed blue lines in both bar plots indicate an occurrence count of 5, while the outer dashed blue lines mark 10. This phylogeny tree was visualized and annotated using iTOL V6.6[4].

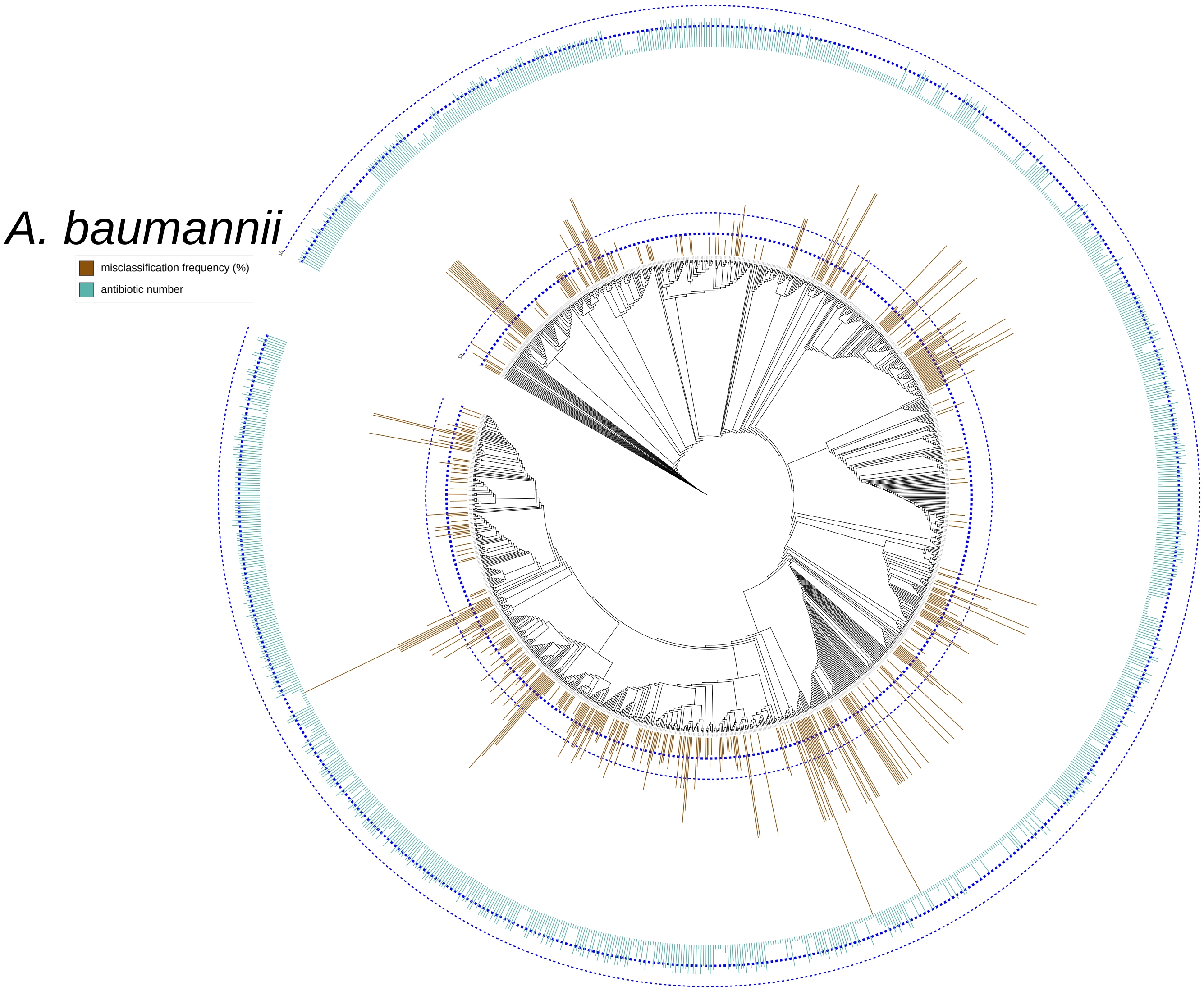


**Figure S18.** Distribution of misclassified genomes of *A. baumannii* on the phylogeny tree. The height of the brown bar for each leave (i.e. a genome) stands for the misclassification frequency ratio that the genome was correctly predicted with random folds but misclassified with phylogeny-aware or homology-aware folds, normalized by the number of antibiotic combinations the genome was involved, which is 7, and by the number of ML-based AMR prediction methods, which is four. The height of the green bar for each leave (i.e. a genome) stands for the number of antibiotics for which the genome's phenotype was annotated. The inner dashed blue lines in both bar plots indicate an occurrence count of 5, while the outer dashed blue lines mark 10. This phylogeny tree was visualized and annotated using iTOL V6.6[4].

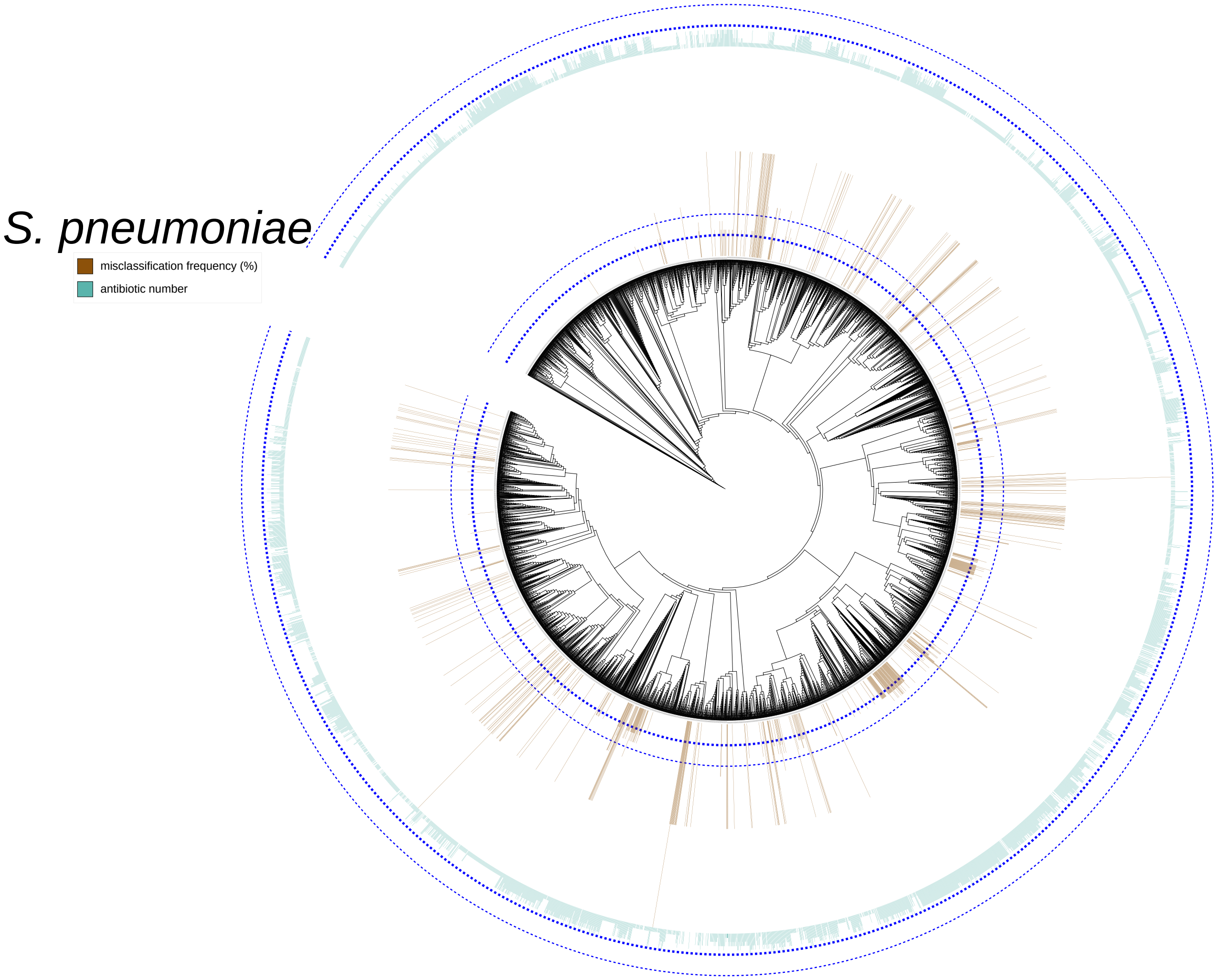


**Figure S19.** Distribution of misclassified genomes of *S. pneumoniae* on the phylogeny tree. The height of the brown bar for each leave (i.e. a genome) stands for the misclassification frequency ratio that the genome was correctly predicted with random folds but misclassified with phylogeny-aware or homology-aware folds, normalized by the number of antibiotic combinations the genome was involved, which is 5, and by the number of ML-based AMR prediction methods, which is four. The height of the green bar for each leave (i.e. a genome) stands for the number of antibiotics for which the genome's phenotype was annotated. The inner dashed blue lines in both bar plots indicate an occurrence count of 5, while the outer dashed blue lines mark 10. This phylogeny tree was visualized and annotated using iTOL V6.6[4].

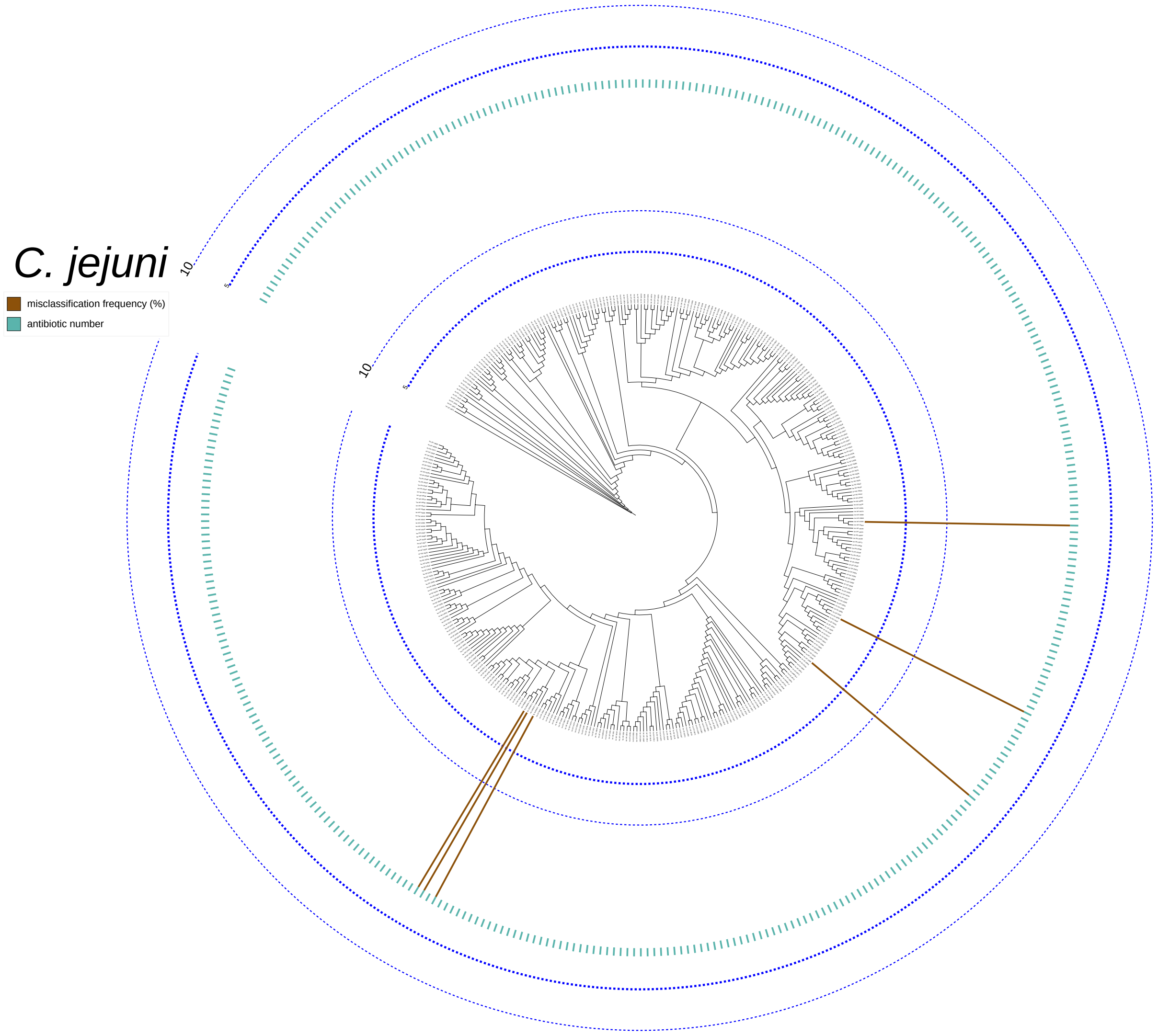


**Figure S20.** Distribution of misclassified genomes of *C. jejuni* on the phylogeny tree. The height of the brown bar for each leave (i.e. a genome) stands for the misclassification frequency ratio that the genome was correctly predicted with random folds but misclassified with phylogeny-aware or homology-aware folds, normalized by the number of antibiotic combinations the genome was involved, which is 1, and by the number of ML-based AMR prediction methods, which is four. The height of the green bar for each leave (i.e. a genome) stands for the number of antibiotics for which the genome's phenotype was annotated. The inner dashed blue lines in both bar plots indicate an occurrence count of 5, while the outer dashed blue lines mark 10. This phylogeny tree was visualized and annotated using iTOL V6.6[4].

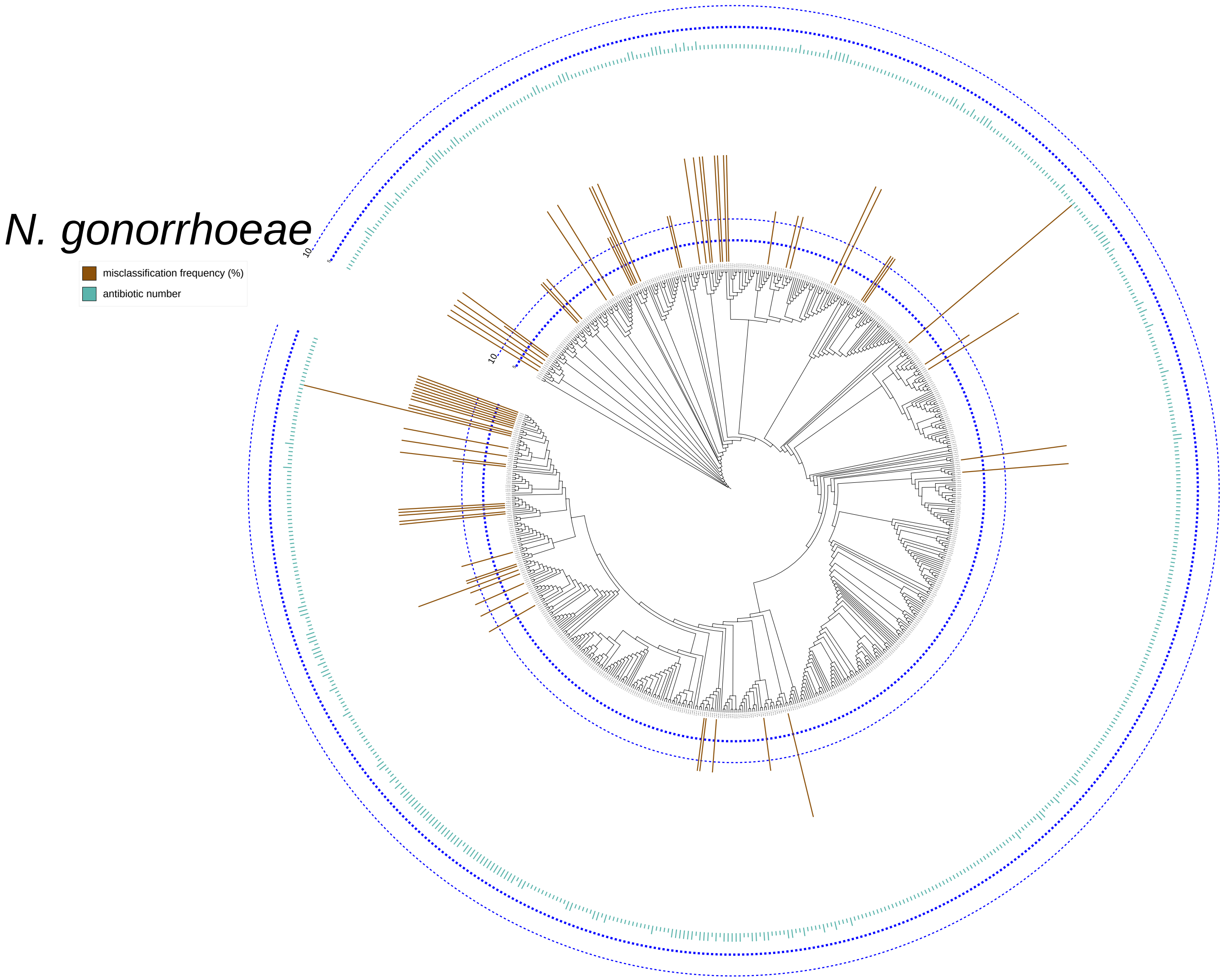


**Figure S21.** Distribution of misclassified genomes of *N. gonorrhoeae* on the phylogeny tree. The height of the brown bar for each leave (i.e. a genome) stands for the misclassification frequency ratio that the genome was correctly predicted with random folds but misclassified with phylogeny-aware or homology-aware folds, normalized by the number of antibiotic combinations the genome was involved, which is 2, and by the number of ML-based AMR prediction methods, which is four. The height of the green bar for each leave (i.e. a genome) stands for the number of antibiotics for which the genome's phenotype was annotated. The inner dashed blue lines in both bar plots indicate an occurrence count of 5, while the outer dashed blue lines mark 10. This phylogeny tree was visualized and annotated using iTOL V6.6[4].

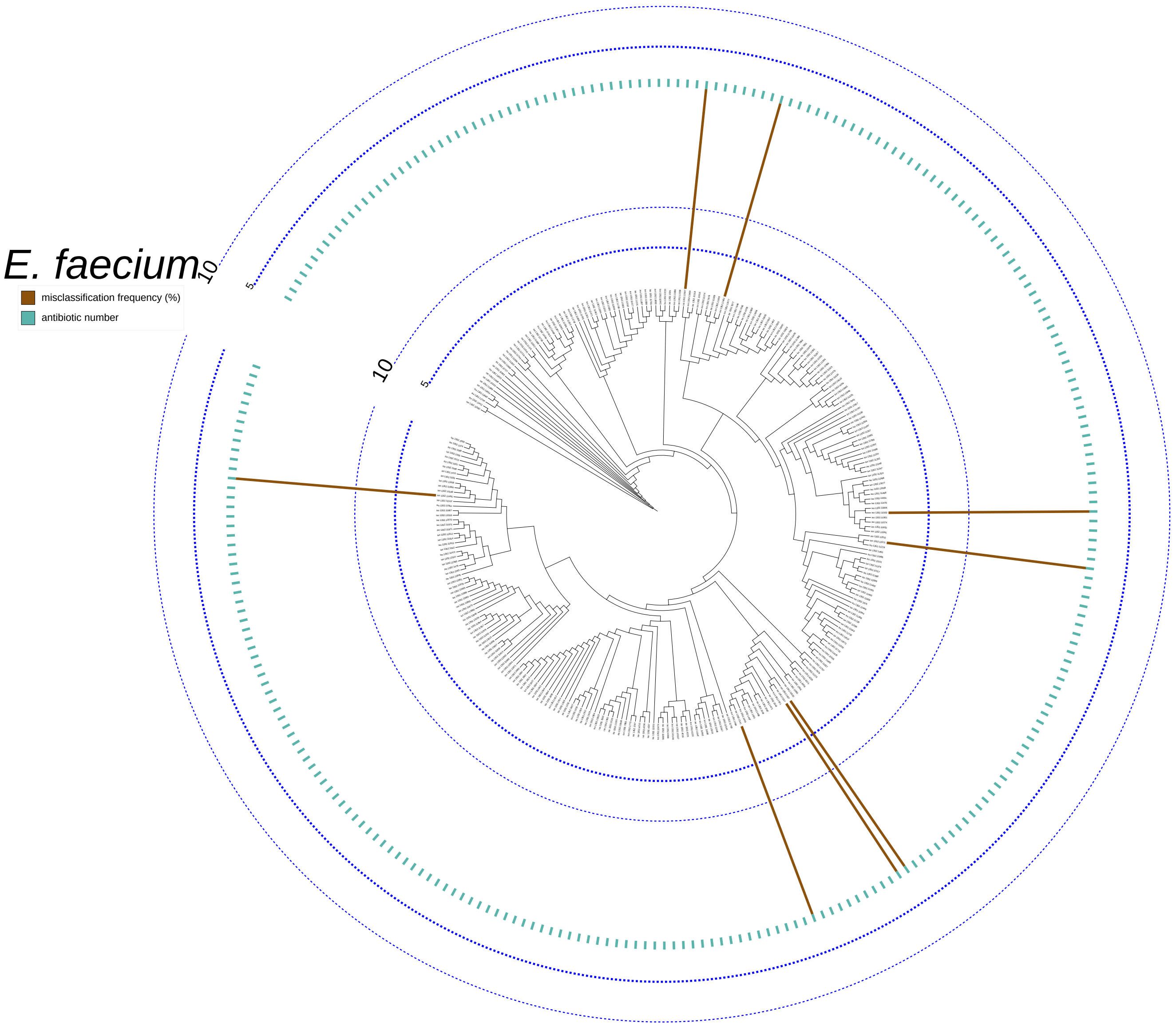


**Figure S22.** Distribution of misclassified genomes of *E. faecium* on the phylogeny tree. The height of the brown bar for each leave (i.e. a genome) stands for the misclassification frequency ratio that the genome was correctly predicted with random folds but misclassified with phylogeny-aware or homology-aware folds, normalized by the number of antibiotic combinations the genome was involved, which is 1, and by the number of ML-based AMR prediction methods, which is four. The height of the green bar for each leave (i.e. a genome) stands for the number of antibiotics for which the genome's phenotype was annotated. The inner dashed blue lines in both bar plots indicate an occurrence count of 5, while the outer dashed blue lines mark 10. This phylogeny tree was visualized and annotated using iTOL V6.6[4].

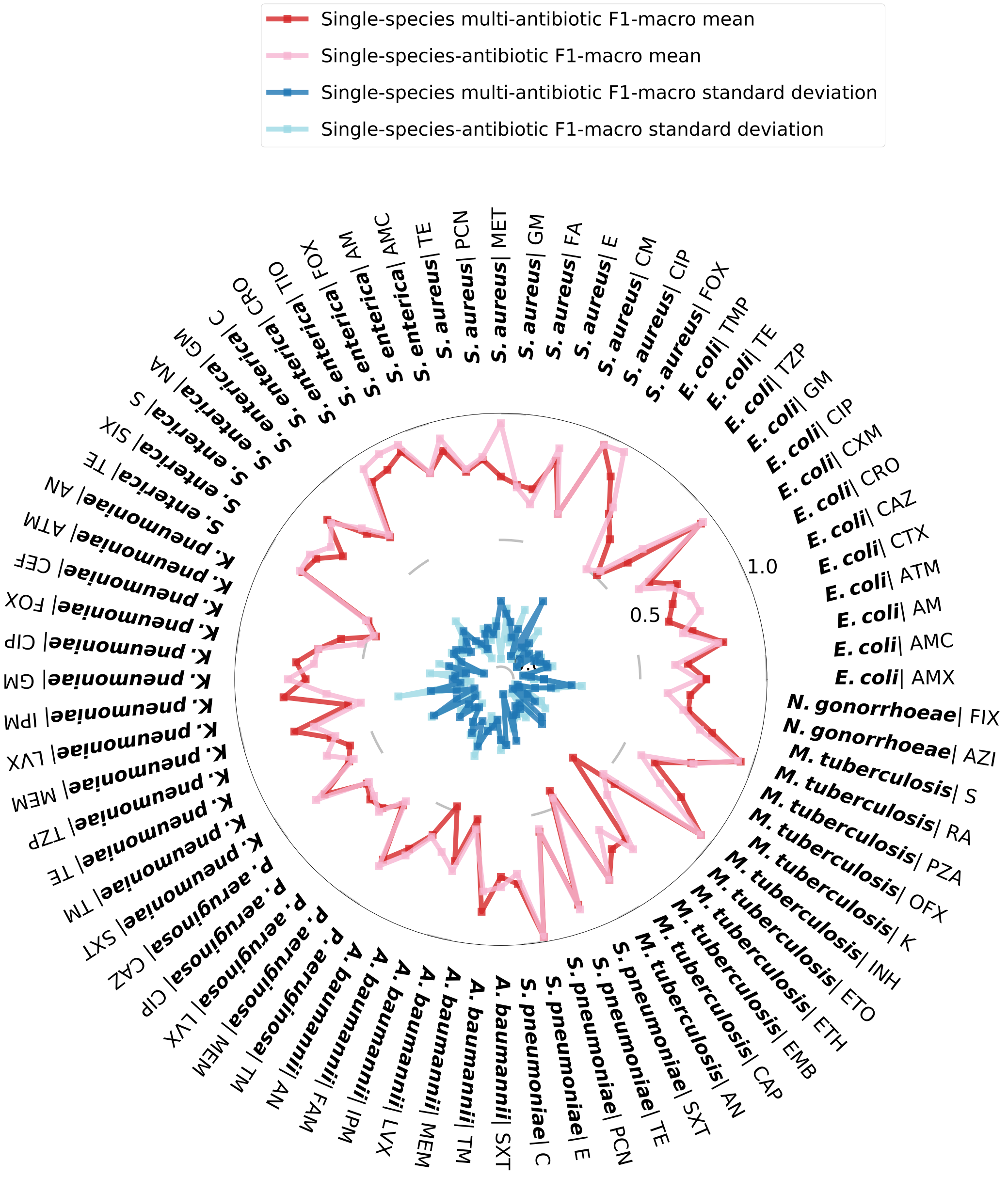


**Figure S23.** Comparison of the performance (F1-macro mean and standard deviation) between the Aytan-Aktug single-species models with single or multiple antibiotics. Both models were evaluated with homology-aware folds, whereas the samples in each fold differ. The mapping of the acronyms to antibiotics is in Supplemental Table S2.

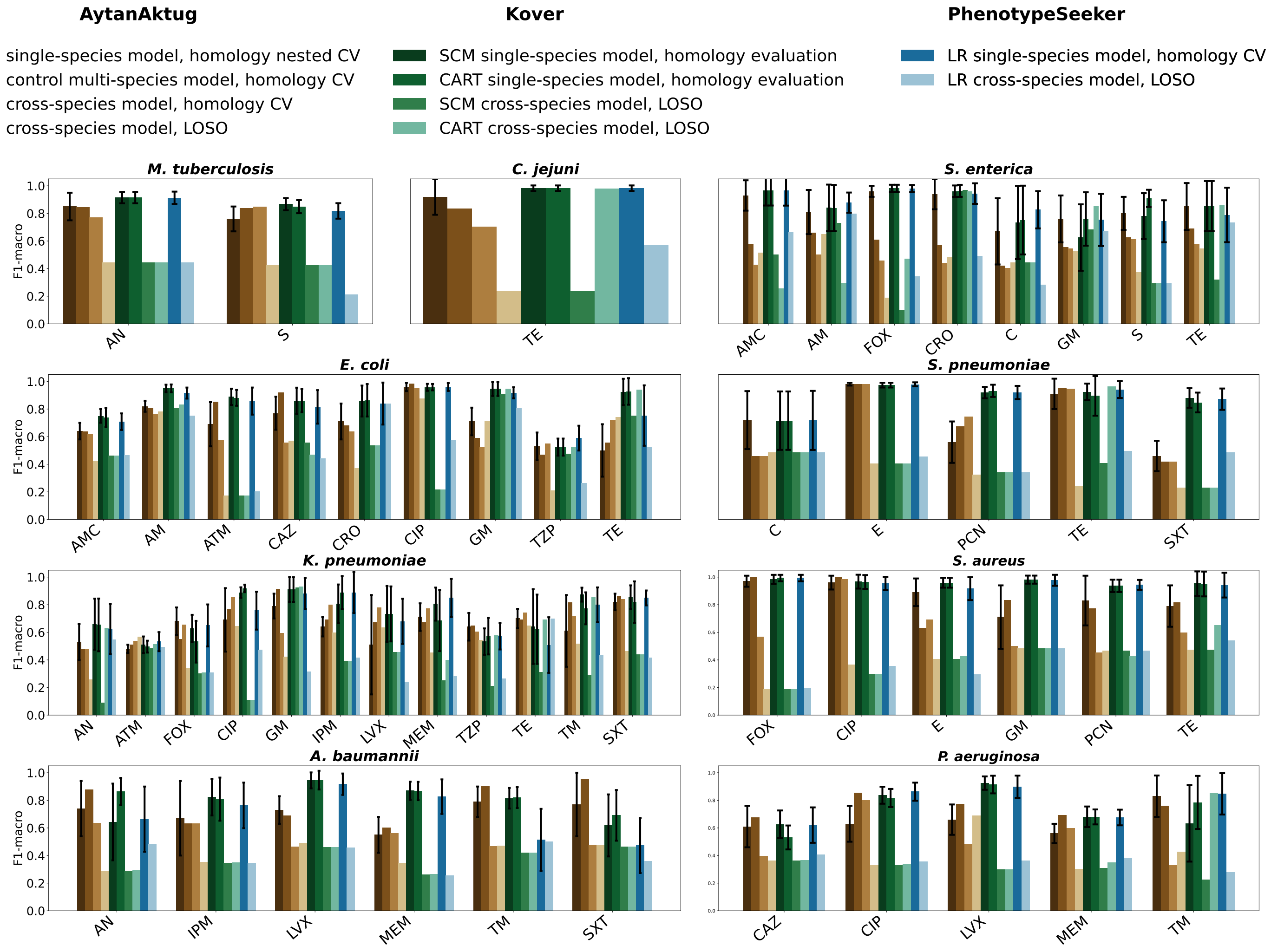


**Figure S24.** Performance (F1-macro) of the multi-species model using Aytan-Aktug, Kover, and PhenotypeSeeker, in comparison with the corresponding single-species model (F1-macro mean and standard deviation). The mapping of the acronyms to antibiotics is in Supplemental Table S2.

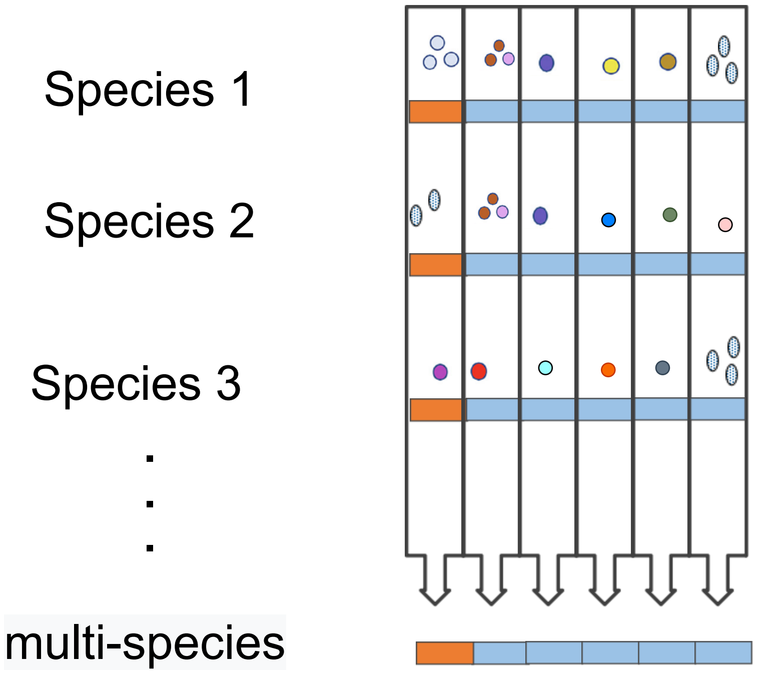


**Figure S25.** Sample partitioning method for the Aytan-Aktug control multi-species and cross-species models. We partitioned samples of each species into 6 folds, resulting in 54 folds (9 species $\times$ 6 folds). And then we picked a fold away from the fold set belonging to each of the 9 species to form a new fold, consisting of samples of 9 species. We conducted the picking procedure iteratively six times, generating a set of six folds. Each fold comprises samples from nine different species.

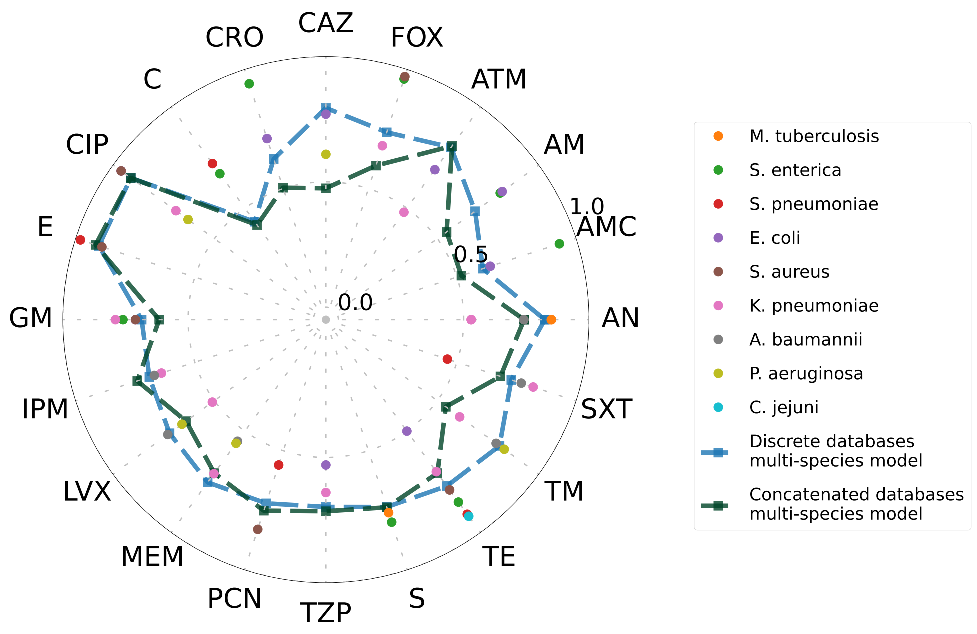


**Figure S26.** The performance (F1-macro) of the Aytan-Aktug control multi-species model, cross-species model, and single-species model compared by Aytan-Aktug original methods. The multi-species model's performance for each antibiotic was computed based on samples from multiple species. This prevented researchers from figuring out if there was any difference between single-species and multi-species models for a specific species-antibiotic combination. The mapping of the acronyms to antibiotics is in Supplemental Table S2.

4. Ivica Letunic and Peer Bork. Interactive tree of life (itol) v5: an online tool for phylogenetic tree display and annotation. *Nucleic acids research*, 49(W1):W293–W296, 2021.
