## Supplemental Table S1 for "Assessing computational predictions of antimicrobial resistance phenotypes from microbial genomes"

**Table S1.** Machine learning-based studies on AMR phenotypic determination from genome data.

| Method | Year | Pathogen (No.) | Feature [a] | Classifier[b] | Type[c] | comparison |
|---|---|---|---|---|---|---|
| Green et al. [1]* | 2022 | *M.tuberculosis*(23,049) | AMR loci | multi-drug CNN single-drug CNNs | phenotype | LR, wide and deep neural network[2], WHO catalog[3] |
| Ren et al. [4] * | 2022 | *E. coli*(987+1,509) | SNPs, | SVM,LR,RF,NN | phenotype | - |
| GenTB[5] * | 2021 | *M.tuberculosis* (train:1397+360, benchmark:20,408) | SNPs(AMR) | RF, wide and deep NN[2] | phenotype | TB-profiler[6], Mykrobe[7] |
| INGOT-DR[8] * | 2021 | *M.tuberculosis*(8,000) | SNPs | novel ML algorithm | phenotype | RF,LR,SVM, Kover[9] |
| Aytan-Aktug et al. [10] * | 2021 | *K. pneumoniae*(1,640) *M. tuberculosis*(2,497), *S. enterica*(1,981) | SNPs (partial & whole) | RF | phenotype | - |
| Yang et al. [11] | 2021 | *M.tuberculosis*(13,402) | SNPs(AMR) | heterogeneous graph attention network | phenotype | DA[12],SVM,LR, multi-lable RF[13] |
| Seq2Geno2Pheno [14] * | 2020 | *P. aeruginosa*(414) | SNPs, gene expression, pan-genome GPA, indels | SVM, RF, LG | phenotype | - |
| Van Camp et al. [15]+ | 2020 | *A. baumannii*(256+1) *E. coli*(330+11), *E. cloacae*(67+2), *K. aerogenes*(5+31), *K. pneumoniae*(211) | ARGs clusters | XGBoost | phenotype | LASSO and Ridge regression |
| Nguyen et al. [16](PATRIC)* | 2020 | *K. pneumoniae*(1,667), *M. tuberculosis*(5,353), *S. enterica*(1,999), *S. aureus*(1,274) | k-mers(core gene without AMR genes) | XGBoost | phenotype | full contig model |
| VAMPr[17]* | 2020 | 9 pathogens(3,393) | KEGG-ortholog -gene-based sequence variants | XGBoost | phenotype | PATRIC[18] (*K. pneumoniae*), with and without imbalance handling[19], elastic net,adaboost,svm, KNN,NN |
| Hyun et al. [20] | 2020 | *S. aureus*(288), *E. coli*(1,588), *P. aeruginosa*(456) | pan-genome GPA, SNPs(core gene) | SVM ensembles | phenotype | - |
| Aytan-Aktug et al. [21] * | 2020 | *M. tuberculosis*(3,528), *E. coli*(1,694), *S. enterica* (658), *S. aureus*(1,236 | SNPs(AMR), acquired ARGs | NN, RF | phenotype | Point-/Resfinder[22, 23] |
| Pataki et al. [24]* | 2020 | *E. coli*(704) | SNPs, ARGs | LR,RF | MIC | - |
| Liu et al. [25] | 2020 | *A. pleuropneumoniae*(96) | kmer(AMR), kmer | SVM, Kover[26] | phenotype | Kover[26] |
| Kouchaki et al. [13]* | 2020 | *M.tuberculosis*(13,402) | SNPs(AMR) | multi-label RF, single-label RF | phenotype | DA[12] |
| Kover[9]([26])* | 2019 (2016) | 12 pathogens and 56 antibiotics | k-mers | Classification and Regression Trees, Set Covering Machines | phenotype | LR,SVM,NB,Majority |
| Chen et al. [2]* | 2019 | *M. tuberculosis*(3,601) | SNPs(AMR), rare variants | wide and deep NN[27], MLP | phenotype | LR,RF, DA[12] |
| Nguyen et al. [28](PATRIC) + | 2019 | nontyphoidal Salmonella(5,278) | k-mers | XGBoost | MIC | - |
| Deelder et al. [29] | 2019 | *M.tuberculosis*(16,688) | SNPs, SNPs(AMR) | LR,DT,XGBoost | phenotype | TB-Profiler[6, 30], ML tools[2, 31, 32, 33] |
| Yang et al. [32]* | 2019 | *M.tuberculosis*(8,388) | SNPs(AMR), SNPs(AMR) subsets | multi-task deep denoising auto-encoder | phenotype | DA[12],RF,SVM, multi-label KNN, ensemble classification chain |
| Kouchaki et al. [33]** | 2019 | *M.tuberculosis*(13,402) | SNPs(AMR)[31] | RF, Adaboost, GBT,SVM, product-of-marginals | phenotype | DA[12] |
| PhenotypeSeeker [34]* | 2018 | *K. pneumoniae* (167), *P. aeruginosa*(200), *C. difficile*(459) | k-mers | linear regression, LR(RF,SVM) | phenotype | SEER[35], Kover[26] (compare on: AMR SNPs detection, running time) |

**Table S1.** (Continued) Machine learning-based studies on AMR phenotypic determination from genome data.

| Method | Year | Pathogen (No.) | Feature [a] | Classifier[b] | Type[c] | comparison |
|---|---|---|---|---|---|---|
| Nguyen et al. [18] (PATRIC)+ | 2018 | *K. pneumoniae*(1,668) | k-mers (whole, ARG-based, non-ARG-based) | XGBoost | MIC | AdaBoost, bagging, RF, ERT,SVM |
| Her and Wu [36] | 2018 | *E. coli*(59) | pan-genome GPA; CARD[37] gene clusters; (genetic algorithm -based selection) | SVM,NB,RF, Adaboost | phenotype | SVM based on genes from Tyson et al. [38], and gene clusters by Scoary[39] |
| Moradigaravand et al. [40]* | 2018 | *E. coli*(1936) | pan-genome GPA, SNPs(core genome ) population structure, year of isolation | LR, NN, RF gradient boosted DT | phenotype | srst2 (CARD) [41] Resdinder[22] |
| Yang et al. [31] * | 2018 | *M. tuberculosis*(1,839) | SNPs(AMR) | LR,SVM,RF, PM,CBMM | phenotype | DA[12] |
| Eyre et al. [42] | 2017 | *N. gonorrhoeae (680)* | AMR determinants | Multivariate linear regression | MIC | - |
| Davis et al. [43] | 2016 | *M. tuberculosis*, *S. pneumoniae*, *A. baumannii*, *S. aureus* | k-mers | AdaBoost | phenotype | - |
| Pesesky et al. [44] | 2016 | *Enterobacteriaceae* (78) | annotated ARGs | LR | phenotype | annotating database: Resdinder[22], CARD[45], HMMER3[46] |
| Drouin et al. [26] (Kover) | 2016 | *C. difficile*, *M.tuberculosis*, *P. aeruginosa*, *S. pneumoniae* | k-mers | Set Covering Machines | phenotype | $(\chi^2+)CART$, $(\chi^2+)SVM$, Majority |
| Li et al. [47] (Li et al. [48]) | 2016 (2017) | *S. pneumoniae*(2,528) *S. pneumoniae*(4,309) | penicillin-binding proteins types | mode, RF, elastic net (mode,RF) | MIC | - |
| Rishishwar et al. [49] | 2014 | *S. aureus*(25) | ARGs | LR | phenotype | J48,LR,RF,SVM,NN,NB |

[*] The * symbol denotes that the corresponding software is open-source.
[**] The ** symbol denotes that the corresponding software is claimed to be open-source, but the site was not accessible due to a broken URL during October and November 2023.
[+] The + symbol denotes that the corresponding software is only provided as a trained pipeline.
[a] Abbreviations: GPA: gene presence and absence; ARGs: AMR-associated genes or regions; SNPs: single nucleotide polymorphisms, including insertion/deletion. And SNPs or chromosomal mutation is used interchangeably here; SNPs(AMR): non-genome-wide SNPs, which can be either AMR-associated SNPs or SNPs identified from ARGs. And the AMR information can be based on either expert-judged published literatures or statistically associated published databases.
[b] Abbreviations: NB: Naive Bay, SVM: Support-vector machine; LR: logistic regression; RF: random forest; PM:product-of-marginals model; CBMM:class-conditional Bernoulli mixture model; NN: neural networks; CNN: convolutional neural networks; DT: decision tree; ERT: extremely randomized trees, XGBoost: extreme gradient boosting tree; GBT: gradient tree boosting; MLP: deep multilayer perceptron; KNN: k-nearest neighbor: DA: direct association based on a database.
[c] phenotype: phenotype prediction; MIC: minimum inhibitory concentration (MIC) prediction. Some studies also include AMR determinants analysis, which is beyond the focus of this table.
[d] https://blast.ncbi.nlm.nih.gov/Blast.cgi?PAGE=Proteins

3. Timothy M Walker, Paolo Miotto, Claudio U Köser, Philip W Fowler, Jeff Knaggs, Zamin Iqbal,

Martin Hunt, Leonid Chindelevitch, Maha R Farhat, Daniela Maria Cirillo, et al. The 2021 who catalogue of mycobacterium tuberculosis complex mutations associated with drug resistance: a genotypic analysis. *The Lancet Microbe*, 3(4):e265–e273, 2022.
