## Supplemental Table S11 for "Assessing computational predictions of antimicrobial resistance phenotypes from microbial genomes"

**Table S12.** Dataset folds potentially suffering from ill-defined F1-negative and F1-positive metrics under phylogeny-aware and homology-aware evaluation.

| | F1-positive | F1-negative |
|---|---|---|
| Homology-aware folds | *M. tuberculosis*: 1 fold in amikacin[a], pyrazinamide, rifampicin, respectively, 6 folds in capreomycin, 2 folds in ethionamide, streptomycin, ethiomide, respectively; *S. enterica*: 1 fold in cefoxitin, ceftiofur, ceftriaxone, chloramphenicol, gentamicin, nalidixic acid, respectively; *N. gonorrhoeae*: 1 fold in azithromycin, 5 folds in cefixime; *S. aureus*: 1 fold in gentamicin, tetracycline, respectively; *K. pneumoniae*: 1 fold in amikacin; *A. baumannii*: 1 fold in ampicillin/sulbactam. | *M. tuberculosis*: 2 folds in ethiomide; *S. aureus*: 1 fold in cefoxitin. |
| Phylogeny-aware folds | *E. coli*: 2 folds in cefotaxime, 1 fold in ceftazidime and ciprofloxacin; *N. gonorrhoeae*: 2 folds in cefixime; *S. pneumoniae*: 1 fold in chloramphenicol and tetracycline, respectively. | *S. aureus*: 4 folds in cefoxitin, 1 fold in ciprofloxacin; *K. pneumoniae*: 3 folds in ciprofloxacin, 1 fold in levofloxacin; *A. baumannii*: 2 folds in evofloxacin, 1 fold in meropenem and tobramycin, respectively. |

[a] It means F1-positive was ill-defined in one fold in the data set representing the combination of *M. tuberculosis* and amikacin.
